# Transposable element expression with variation in sex chromosome number: insights into a toxic Y effect on human longevity

**DOI:** 10.1101/2023.08.03.550779

**Authors:** Jordan Teoli, Miriam Merenciano, Marie Fablet, Anamaria Necsulea, Daniel Siqueira-de-Oliveira, Alessandro Brandulas-Cammarata, Audrey Labalme, Hervé Lejeune, Jean-François Lemaitre, François Gueyffier, Damien Sanlaville, Claire Bardel, Cristina Vieira, Gabriel AB Marais, Ingrid Plotton

## Abstract

Why women live longer than men is still an open question in human biology. Sex chromosomes have been proposed to play a role in the observed sex gap in longevity, and the Y male chromosome has been suspected of having a potential toxic genomic impact on male longevity. It has been hypothesized that in aging individuals, TE repression is diminished, which could lead to detrimental effects (e.g. somatic mutations, perturbed gene expression) and to an acceleration of the aging process. As the Y chromosome is typically enriched in transposable elements (TE), this could explain why the presence of a Y chromosome is associated with shorter longevity. Using transcriptomic data from humans with atypical karyotypes, we found an association between TE expression and the presence and number of Y chromosomes. These findings are consistent with the existence of a toxic Y effect on men's longevity.

## Introduction

Differences in longevity between sexes are prevalent across the tree of life, a phenomenon known as sex gap in longevity (SGL) (Lemaître et al. 2020). In both vertebrates and invertebrates, the heterogametic sex (males in XY, and females in ZW systems) usually has a reduced longevity compared to the homogametic sex (Lemaître et al. 2020). This SGL is also observed in humans, where on average, life expectancy at birth is five years longer for women than for men (World Health Organization 2023). The female-biased gap in longevity is consistently observed in nearly all human populations (Rochelle et al. 2015), and it explains the increased prevalence of women among supercentenarians (Willcox, Willcox, and Ferrucci 2008).

Although cultural factors favor the extended longevity in women, several biological hypotheses have also been proposed to underlie the SGL phenomenon, which are not mutually exclusive (Rochelle et al. 2015; Luy 2003). First, the sex-specific production of hormones (e.g. androgens) has been suggested to contribute to sexual dimorphism in longevity. In humans, it was found that the removal of sex-specific hormones increases male longevity (Maklakov and Lummaa 2013). Second, due to the exclusive maternal transmission of mitochondria, natural selection cannot target deleterious mutations in the mitochondrial genome that specifically impact male fitness. These mutations can thus freely accumulate and may reduce male longevity, a phenomenon called the mother’s curse (Maklakov and Lummaa 2013; Milot et al. 2017; Frank and Hurst 1996). Third, since the heterogametic sex carries one variant of each sex chromosome, it is consequently more susceptible to recessive X-linked (or Z-linked) deleterious genetic mutations, as proposed by the unguarded X hypothesis (Trivers 1985). Both theory and empirical data suggest that the latter mechanism explains a part of the sex gap in longevity (Connallon et al. 2022).

Finally, the toxic Y hypothesis has been recently proposed to explain the existence of SGL. This hypothesis relies on the high transposable element (TE) content in Y (or W) chromosomes (reviewed in (Marais et al. 2018; Marais, Lemaitre, and Vieira 2020)). Y (or W) chromosomes frequently exhibit lower rates of recombination, leading to an accumulation of repeated sequences (mainly TEs) on these chromosomes (Erlandsson, Wilson, and Pääbo 2000). TEs are DNA sequences with the ability to move within the genome, highly enriched in heterochromatin, and that contribute to an impressive 44-66% of the human genome sequence (Bennett et al. 2004; Guio and González 2019). In humans, it is estimated that approximately 100 insertions belonging to the long interspersed nuclear elements (LINE-1) family contribute to nearly all observed transposition activities, including the retrotransposition of nonautonomous short interspersed nuclear elements (SINEs) that rely on proteins encoded by LINE-1 elements (Deniz, Frost, and Branco 2019). While many TEs have a neutral impact on the host, certain insertions can interfere with gene function or result in detrimental chromosomal rearrangements (Deniz, Frost, and Branco 2019; Bourque et al. 2018). Epigenetic silencing mechanisms exist to prevent TE expression and transposition like DNA methylation, histone modifications and the production of small interfering RNAs (Slotkin and Martienssen 2007). However, aging has been shown to disrupt this epigenetic regulation at constitutive heterochromatin causing an increased TE activation in several organisms including humans (De Cecco, Criscione, Peterson, et al. 2013; De Cecco, Criscione, Peckham, et al. 2013; Van Meter et al. 2014; Dennis et al. 2012; Maxwell, Burhans, and Curcio 2011; Li et al. 2013; Chen et al. 2016; Brown, Nguyen, and Bachtrog 2020), thus enhancing the probability to induce detrimental effects through somatic mutations or through other mechanisms (e.g. accumulation of TE transcripts into the cytoplasm, interaction between TE transcripts and genes/proteins) (Mosaddeghi, Farahmandnejad, and Zarshenas 2023). Since the Y chromosome is rich in TEs, more TEs might become active in old males compared to old females, generating more detrimental effects, accelerating aging and likely reducing longevity in males.

The toxic Y hypothesis has already been investigated in the fruit fly *Drosophila melanogaster*, where TE expression was found to be higher in old males compared to old females (Brown, Nguyen, and Bachtrog 2020). This observation was associated with a loss of heterochromatin in repetitive elements during aging in male flies (Brown, Nguyen, and Bachtrog 2020). Moreover, flies with additional Y chromosomes showed decreased longevity, further suggesting that the number of Y chromosomes influences organismal survival in this species (Brown, Nguyen, and Bachtrog 2020). However, another recent study using flies with variable levels of heterochromatin in the Y chromosome pointed out that the presence, number, or size of the Y chromosome has no impact on sexual dimorphism in longevity (Delanoue et al. 2023). Hence, the harmful effects of the Y chromosome on longevity are still under debate.

In humans, the Y chromosome is relatively small (∼57 Mb) compared to the X chromosome (∼156 Mb), but it is also extremely rich in TE insertions (Skaletsky et al. 2003; “Human Genome Assembly GRCh38.P14,” n.d.). Furthermore, men with 47,XYY and 47,XXY abnormal karyotypes (with an extra Y or X chromosome, respectively) have strikingly different longevities: 47,XYY men were associated with a 10 year reduction in longevity while 47,XXY men were associated with a two year reduction compared to 46,XY individuals (Bojesen et al. 2004; Stochholm, Juul, and Gravholt 2010). This was observed despite the extra Y having smaller observable effects on 47,XYY individual’s biology and health than the extra X in 47,XXY individuals, which results in the Klinefelter syndrome (Bojesen et al. 2004; Stochholm, Juul, and Gravholt 2010). Yet, in humans, information is lacking about the contribution of TEs in longevity, and thus the possible toxic Y effect in this species.

In this work, we first studied whether TE expression is associated with the number of Y chromosomes. We analyzed transcriptomic data obtained from individuals of different karyotype compositions: 46,XX females (normal female karyotype), 46,XY males (normal male karyotype), as well as males with abnormal karyotypes, such as 47,XXY and 47,XYY. We found that the presence and number of Y chromosomes might be associated with increased TE expression. This tendency was observed for several TE subfamilies analyzed. We also tested whether there is an increased TE activation in old men compared to old women by using published transcriptomic data of males and females with normal karyotypes (46,XY for males and 46,XX for females, respectively). However, we did not find an increased TE expression in old males compared to old females probably due to the heterogeneity of the dataset. Overall, this work suggests an association between the Y chromosome and the increased TE activity in males and opens a new window to study the toxic effect of this particular chromosome in human longevity.

## Results

### Males with sex aneuploidies (XXY and XYY) have higher transcriptomic differences than XY males compared to XX females

The toxic Y hypothesis is based on the fact that increased TE expression in males is likely the result of the significantly higher insertion content in the Y chromosome. This implies a potential correlation between the number of Y chromosomes in a karyotype and the levels of TE expression. To test if an association between TE expression and the number of Y chromosomes exists in humans, we generated RNA-seq data from blood samples of males (46,XY) and females (46,XX) from the Lyon University Hospital. In addition, we also generated RNA-seq data from males with different sex chromosome aneuploidies: males with Klinefelter syndrome harboring an additional X chromosome (47,XXY), and males with Jacob syndrome harboring an additional Y chromosome (47,XYY). In total, we obtained blood samples from 24 individuals (six females 46,XX, six males 46,XY, eight males 47,XXY, and four males 47,XYY) (**Table S1**). Pairwise Wilcoxon tests revealed a significant difference in age distribution only between 46,XX and 47,XXY individuals (p-value = 0.033). This finding suggests that age could act as a confounding factor in the relationship between TE expression and karyotype. However, it is worth noting that any such conclusion would introduce a conservative bias when evaluating the correlation between TE expression and the number of Y chromosomes in a karyotype. Notably, individuals with 47,XXY (mean age 23.4 years) and 47,XYY (mean age 29.5 years) tend to be younger compared to those with 46,XX (mean age 39.2 years) and 46,XY (mean age 41.5 years) karyotypes.

We first analyzed gene expression to see whether different karyotypes have changes in their transcriptomic profile (**Fig. S1**) (see **Materials and Methods**). Principal component analysis (PCA) showed that the highest variation in gene expression was found when comparing females (46,XX) and males (with and without aneuploidies) (**Fig. S2**). Besides that, we found that the number of significantly differentially expressed genes (DEGs) varied depending on the pair of karyotypes compared (**Fig. S3, Table S2**). Comparing males 46,XY to females 46,XX samples, 19 (0.10%) and 18 (0.09%) genes were found to be upregulated and downregulated, respectively, among the 19,947 studied genes (**Table S2**). As expected, we found that while most of the upregulated genes (13/19, 68.42%) were Y-linked genes, most of the downregulated ones (14/18, 77.78%) were X-linked genes (**Fig. S4, Tables S2 to S4, Data S1 and S2**). Furthermore, we found an increased number of DEGs when comparing samples from individuals with sex chromosome aneuploidies (47,XXY and 47,XYY males) to female (46,XX) than to male (46,XY) samples. Indeed, 47,XXY and 47,XYY males showed 1,893 (9.49%) and 1,510 (7.57%) DEGs compared to 46,XX females, respectively. However, 47,XXY and 47,XYY males only showed 141 (0.71%) and 58 (0.29%) DEGs compared to 46,XY males, respectively (**Fig. S3, Table S2, Data S2**). We also found a small amount of DEGs when comparing 47,XYY and 47,XXY males (50, 0.25%) (**Fig. S3, Table S2**).

Finally, Gene Ontology (GO) and pathway enrichment analysis found the largest number of significant terms when 47,XXY and 47,XYY were compared to 46,XX but few considering the four other comparisons (**Data S3**). Overall, terms associated with demethylase activity, cell cycle, and in protein or DNA production were the most shared between the six pairwise karyotype comparisons and between the databases used (GO, KEGG or Reactome) (**Fig. S5 to S7**). Interestingly, biological pathways concerning senescence, like DNA damage/telomere stress induced senescence, senescence-associated secretory phenotype, and oxidative stress induced senescence, were found downregulated in 47,XXY karyotype compared to 46,XX, 46,XY and 47,XYY karyotypes (**Fig. S8, Data S3**). Furthermore, some of the DEGs were also previously found in other studies (see **Supplementary text**).

Altogether, we observed that males with 47,XYY and 47,XXY aneuploidies exhibit greater disparities in gene transcriptomic profiles compared to 46,XX females, in contrast to the comparison between 46,XY males and 46,XX females.

### The number of Y chromosomes tend to be associated with an increased TE expression

To check whether the presence of the Y chromosome is associated with overall increased levels of TE expression, we measured TE transcript amounts using the *TEcount* module of *TEtools* (Lerat et al. 2016) in all the RNA samples (**Fig. S1**). PCA showed that females (46,XX) had different TE expression profiles compared to males (46,XY, 47,XXY, or 47,XYY), similar to what we observed in gene expression (**Fig. S9**). Specifically, males (46,XY) tended to have an overall increased TE transcript amount compared to females (46,XX), although it was not statistically significant (Wilcoxon test: p-value = 0.310) (**Fig. 1A**). These results were in line with our expectations, since the Y chromosome is known to harbor many more TE insertions than the X chromosome. We then checked whether the addition of a Y chromosome accentuates our previous findings. Hence, we expected 47,XXY males having more TE transcripts than 46,XX females, and 47,XYY males having more TE transcripts than 46,XY males since the extra Y chromosome should bring more TE insertions to these individuals. In agreement with this hypothesis, we found a significant general increase of TE transcript amounts between 46,XX females and 47,XXY males (Wilcoxon test: p-value = 0.029) or 47,XYY (Wilcoxon test: p-value = 0.010), and a near significant increased between 46,XY and 47,XYY males (Wilcoxon test: p-value = 0.067) (**Fig. 1A**). Differences in TE transcript amounts between 47,XXY and 46,XY, and between 47,XYY and 47,XXY were not significant (Wilcoxon test: p-values = 0.345 and 0.283, respectively). Moreover, we saw that the addition of a sex chromosome, either X or Y, tended to increase the overall amounts of TE transcripts probably due to an increase in genomic material and thus in TE load. However, the addition of a Y chromosome seemed to increase the expression of TEs even more than the addition of a X chromosome (**Fig. 1A**). These results suggested that the presence of the Y chromosome might be associated with an increase in TE transcripts, as postulated in the toxic Y hypothesis, and could contribute to a global deregulation of TEs.

**Figure 1.**
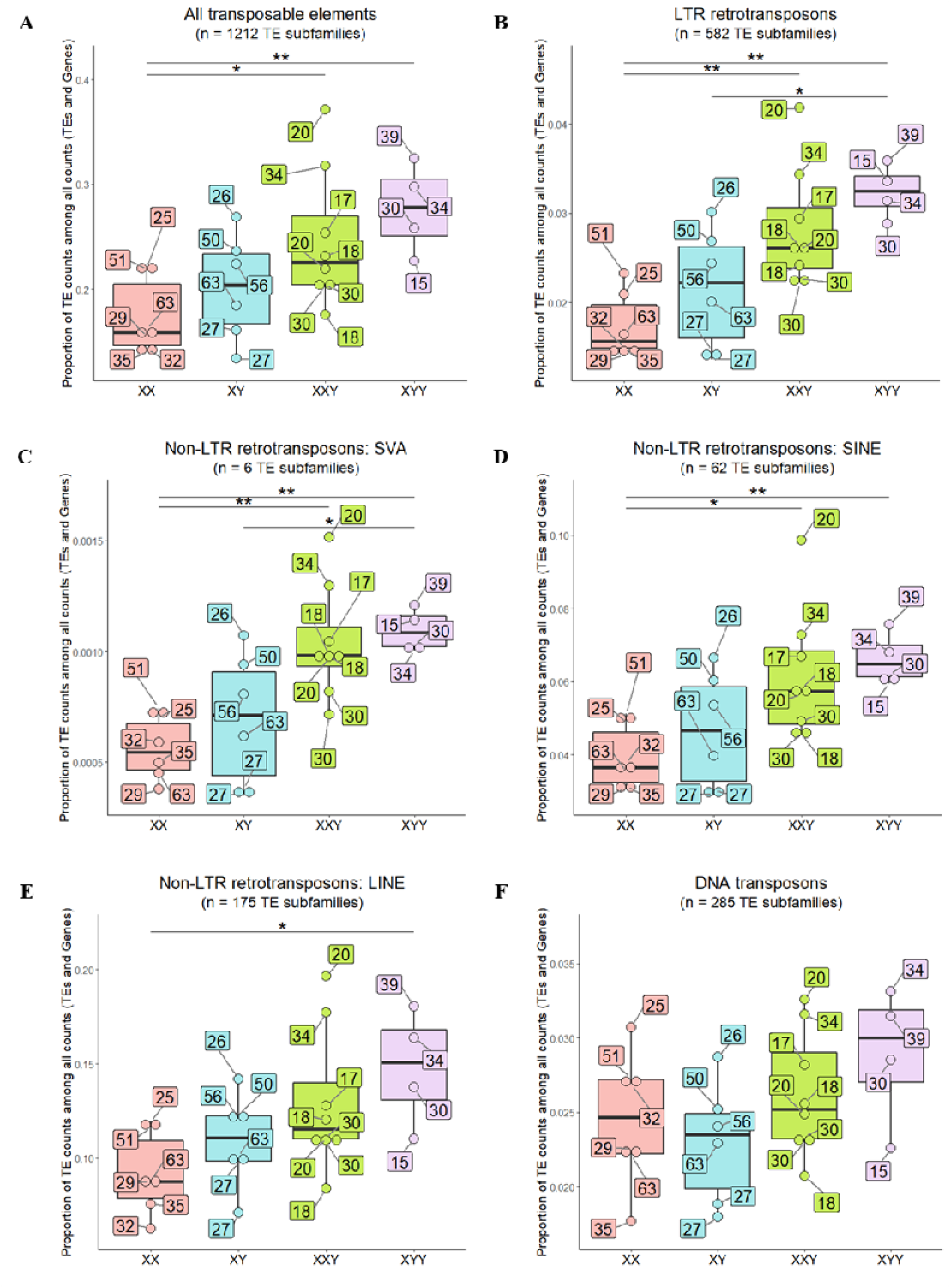
TE expression in the different karyotypes after removing batch effect, considering (A) all TE subfamilies (1,212 TE subfamilies) and (B) TE subfamilies belonging to LTR retrotransposons (582 TE subfamilies), (C) SVA non-LTR retrotransposons (6 TE subfamilies), (D) SINE non-LTR retrotransposons (62 TE subfamilies), (E) LINE non-LTR retrotransposons (175 TE subfamilies), or (F) DNA elements (285 TE subfamilies). _Global TE expression was measured in each karyotype (x-axis) as the proportion of TE read counts among all read counts (TEs and genes) (y-axis). In this calculation, read counts were used after DESeq2 normalization (“normalized counts”) to remove any depth sequencing bias. Each dot represents one individual. Dots are colored according to karyotype (four levels: XX, XY, XXY, XYY). The age of each individual is indicated in a labeled box linked to the corresponding dot. Batch effect was removed before graphical display. P-values from Wilcoxon test comparing pairwise karyotypes: (*) < 0.05 and (**) < 0.01._

We also performed the same analysis as before focusing on TE classes/subclasses (LTR, SVA, SINE, LINE, and DNA elements) (**Fig. 1B to F**). Considering LTR, SVA, and SINEs: both 47,XXY and 47,XYY individuals had a statistically significant increased TE expression compared to 46,XX females (Wilcoxon test: p-values = 0.003 and 0.010 for LTR respectively, p-values = 0.001 and 0.010 for SVA respectively, and p-values = 0.013 and 0.010 for SINE respectively) (**Fig. 1B to D**). There were also significant differences in TE transcript amounts between 46,XY and 47,XYY males for LTR (Wilcoxon test: p-value = 0.019) (**Fig. 1B**) and SVA elements (Wilcoxon test: p-value = 0.038) (**Fig. 1C**). Finally, for LINE elements we only found significant differences between 46,XX females and 47,XYY males (Wilcoxon test: p-value = 0.038) (**Fig. 1E**), and no significant differences in TE expression between any karyotypes for DNA elements (**Fig. 1F**).

In humans, only SVA, HERVK, AluS, L1, and AluY are known to be transcriptionally active (Kojima 2018). We found that the proportion of TE transcripts belonging to these groups was significantly increased in 47,XYY males compared to 46,XX females (Wilcoxon test: p-values = 0.010, 0.010, 0.010, and 0.038 respectively for SVA, HERVK, AluS and L1 elements) (**Fig. S10A to D**), except for AluY elements (**Fig. S10E**). The proportion of TE transcripts was also significantly increased in 47,XXY males compared to 46,XX females in SVA (Wilcoxon test: p-value = 0.001) (**Fig. S10A**), HERVK (Wilcoxon test: p-value = 0.003) (**Fig. S10B**), and AluS families (Wilcoxon test: p-value = 0.020) (**Fig. S10C**). Moreover, 47,XYY males also showed a significant increase in TE transcripts in comparison with 46,XY males for SVA (Wilcoxon test: p-value = 0.038) (**Fig. S10A**) and HERVK families (Wilcoxon test: p-value = 0.019) (**Fig. S10B**). No significant differences in TE expression between any karyotypes considering AluY elements was found (**Fig. S10E**).

A similar tendency in TE expression was found when analyzing differentially expressed TE subfamilies between the karyotypes (**Fig. S11**, **Table S5**). For the vast majority of them, we found a significant increased TE expression in both 47,XXY and 47,XYY individuals compared to 46,XX, with a tendency to find 47,XYY individuals being the ones with the highest TE expression (**Fig. 2 and Fig. S12**). For some TE subfamilies, TE transcripts also significantly increased in 46,XY compared to 46,XX (**Fig. 2**). Indeed, with the exception of some TE subfamilies, female samples (46,XX) were the ones showing less reads mapping to TEs and 47,XYY samples the ones with more reads (**Fig. 2 and Fig. S12**).

**Figure 2.**
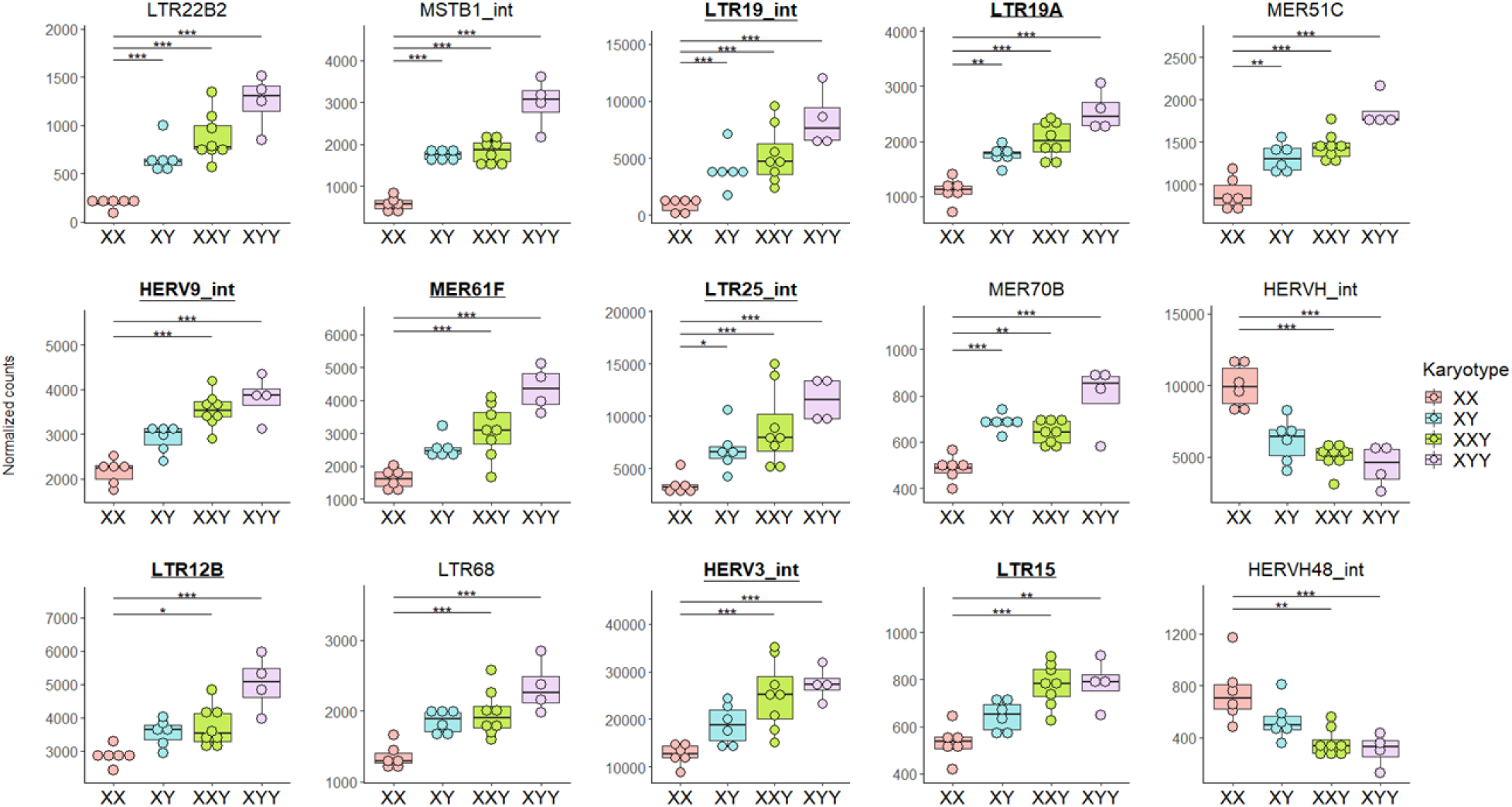
Boxplots for the 15 most significantly differentially expressed TE subfamilies according to karyotype using the likelihood-ratio test, after removing batch effect. Specific TE subfamily expression was estimated using the number of read counts after DESeq2 normalization (“normalized counts”) to remove any depth sequencing bias. Each dot represents one individual. Dots are colored according to karyotype (four levels: XX, XY, XXY, XYY). Batch effect was removed before graphical display. Adjusted p-values from DESeq2: (*) < 0.05, (**) < 0.01, (***) < 0.001. All of these 15 subfamilies had at least one copy on the Y chromosome, except for HERVH48-int, in the TE sequence reference file. All of these 15 subfamilies had at least one copy on the X chromosome in the TE sequence reference file. Underlined bold TE subfamilies are enriched in the Y chromosome (binomial test adjusted p-value < 0.05 and number of observed copies located on Y chromosome > expected). See **Fig. S12** for the other significantly differentially expressed TE subfamilies according to karyotype.

We finally estimated which TE subfamilies were enriched in the Y chromosome among the 1,246 TE subfamilies studied herein (**Data S4**). We found that 9.07% (113/1,246) of all TE subfamilies were enriched in the Y chromosome (binomial test adjusted p-value < 0.05 and number of observed copies located on Y chromosome > expected). Eight TE subfamilies out of the 15 most significant ones according to karyotype (53.33%), were enriched in the Y chromosome and that was greater than expected (p-value < 0.001) (**Fig. 2**). Among the upregulated TE subfamilies found when we compared 46,XY, 47,XXY, or 47,XYY to 46,XX individuals, the proportion of TE subfamilies enriched in the Y chromosome (42.86% (3/7), 26.58% (21/79), and 37.21% (16/43) respectively) were also greater than expected (p-values = 0.020, < 0.001, and < 0.001, respectively) (**Table S5**). Considering downregulated TE subfamilies, we found 12.50% (2/16) of them enriched in the Y chromosome when we compared 47,XXY to 46,XX individuals (p-value = 0.651) and 29.41% (5/17) when we compared 47,XYY to 46,XX individuals greater than expected (p-value = 0.015) (**Table S5**). The increase of TE subfamily expression when the number of copies of this TE subfamily carried by the Y chromosome increased was more pronounced in the Y-enriched TE subfamily group than in the not Y-enriched TE subfamily group (**Fig. S13A**). However, the decrease of TE subfamily expression when the proportion of copies of this TE subfamily carried by the Y chromosome increased, was also more pronounced in the Y-enriched TE subfamily group than in the not Y-enriched TE subfamily group (**Fig. S13B**). Again, we found a significant general increased proportion of TE transcripts in 47,XXY and 47,XYY males compared to 46,XX females and sometimes in 47,XYY males compared to 46,XY males considering separately expression of Y-enriched, Y-depleted, neither Y-enriched nor Y-depleted TE subfamilies, or TE subfamilies with no copies carried by the Y chromosome, the X chromosome or neither (**Fig. S14**).

In summary, these results suggested that TE expression is dependent on the sex chromosomes contained in the karyotype. Particularly, the presence and number of Y chromosomes might be associated with increase TE expression.

### 47,XYY individuals have an enrichment of upregulated TE subfamilies in upstream region of upregulated genes

To see if there was any correlation between differentially expressed TE subfamilies and DEGs, we tested whether insertions from differentially expressed TE subfamilies are enriched or depleted in DEGs upstream regions. To that end, we performed permutation tests for each pairwise karyotype comparison in both upregulated and downregulated genes separately (see **Materials and Methods**). Results revealed an enrichment of TEs from upregulated subfamilies in upstream regions of upregulated genes exclusively in 47,XYY compared to 46,XX karyotype (p-value = 0.017). In contrast, we found a depletion of TE copies from upregulated TE subfamilies in the upstream region of downregulated genes when comparing 47,XXY to 46,XX karyotype (p-value = 0.009). We then used the same approach focusing on TE copies from downregulated TE subfamilies. This time, we only found an enrichment of TE copies in the upstream region of downregulated genes in 47,XYY compared to 46,XX karyotype (p-value = 0.021).

### Some TE transcripts could come from passive co-transcription with genes

In addition to originating from autonomous expression, TE transcripts could also arise from passive co-transcription with genes, such as through intron retention or pervasive intragenic transcription (i.e. non-coding transcription) (Lanciano and Cristofari 2020). In fact, the majority of TE-derived RNA-seq reads typically stem from co-transcription or pervasive transcription (Deininger et al. 2017; Navarro et al. 2019). Hence, we checked whether there were TE transcripts that come from gene transcription rather than autonomous transcription and which could contribute to TE subfamily being differentially expressed. To do that, we tested whether TE copies from upregulated subfamilies were enriched within upregulated genes for each pairwise karyotype comparison. Then, we tested whether TE copies from downregulated subfamilies were enriched within downregulated genes for each pairwise karyotype comparison. When we compared 46,XY males with 46,XX females, we found 2 upregulated genes containing one or more TE insertions from upregulated TE subfamilies, but 0 overlap between downregulated genes and TE copies from downregulated TE subfamilies (**Data S5**). Considering the comparison 47,XXY versus 46,XX, these numbers rose to 228 and 88 respectively. There were 71 and 123 considering the comparison 47,XYY versus 46,XX (**Data S5**). Except when the number of overlap between genes and TE was equal to 0, we found a higher proportion of upregulated genes containing at least one copy of one of the upregulated TE subfamilies and a higher proportion of downregulated genes containing at least one copy of one of the downregulated TE subfamilies in 46,XY versus 46,XX, 47,XXY versus 46,XX, and 47,XYY versus 46,XX comparisons (**Table S6**).

### No significant association of TE expression in blood cells with age

The toxic Y hypothesis also relies on the fact that aging disrupts the mechanisms of epigenetic regulation at constitutive heterochromatin, consequently leading to heightened activation of TE (De Cecco, Criscione, Peterson, et al. 2013; De Cecco, Criscione, Peckham, et al. 2013; Van Meter et al. 2014; Dennis et al. 2012; Maxwell, Burhans, and Curcio 2011; Li et al. 2013; Chen et al. 2016; Brown, Nguyen, and Bachtrog 2020). Given the substantial abundance of TE insertions on the human Y chromosome relative to the X chromosome, we investigated whether age impacts TE expression differently between males and females hypothesizing that old males exhibit more TE expression than old females. As such, we used published RNA-seq data from the Genotype-Tissue Expression (GTEx) project. To minimize data heterogeneity and to maintain consistency with our previous analysis, we selected blood samples exclusively from non-Latino and non-Hispanic individuals without any history of persistent viral infection or dementia (**Fig. S15**) (see **Material and Methods**). We thus compared TE expression in 46,XY males and 46,XX females belonging to five different age groups from 20 to 70 years old. We divided the dataset in two groups: one excluding data obtained from patients with current or prior histories of cancer or cardiovascular diseases (**Table S7**) and the other including them (**Table S8**).

In the no disease dataset, there were no significant differences observed in TE expression levels between the oldest males (]50-70] years, n = 17) and the youngest males ([20-50] years, n = 28) (Wilcoxon test: p-value = 0.371). Similarly, no significant differences were found between the oldest females (]50-70] years, n = 14) and the youngest females ([20-50] years, n = 21) (Wilcoxon test: p-value = 0.454). Moreover, we observed no difference in TE expression between the oldest males and the oldest females (Wilcoxon test: p-value = 0.399). We then studied more precisely the effect of age in TE expression using 10-year ranges (age groups: [20-30], ]30-40], ]40-50], ]50-60], and ]60-70]). Again, global expression of TEs was not significantly associated with sex, age in females, or age in males (p-values = 0.788, 0.585, and 0.471, respectively) (**Fig. 3**). Focusing on TE classes/subclasses (LTR, SVA, SINE, LINE, or DNA elements), active TEs (SVA, HERVK, AluS, L1, or AluY), Y-enriched, Y-depleted, or TEs with no copies in the Y chromosome, we found no significant association between TE expression and both sex and age group. Regarding TE subfamilies, none of them were upregulated in oldest males compared to youngest males or when comparing oldest females with youngest females (**Fig. S16A-B**). However, we found several TE subfamilies significantly differentially expressed between some age groups, particularly in males (**Fig S16B and S17, Table S9**). Interestingly, the 30-40 years old group shows the highest number of significantly differentially expressed TE subfamilies when compared with other age groups in males (**Fig. S16B, Table S9**). In addition, upon comparing TE expression between males and females in every age group, we did not observe an elevated number of TE subfamilies upregulated in males associated with age (**Fig. S16C-G, Table S10**). Only three TE subfamilies were upregulated in males compared to females adjusted on age group, none were upregulated in females compared to males (**Fig. 4, Table S10**).

**Figure 3.**
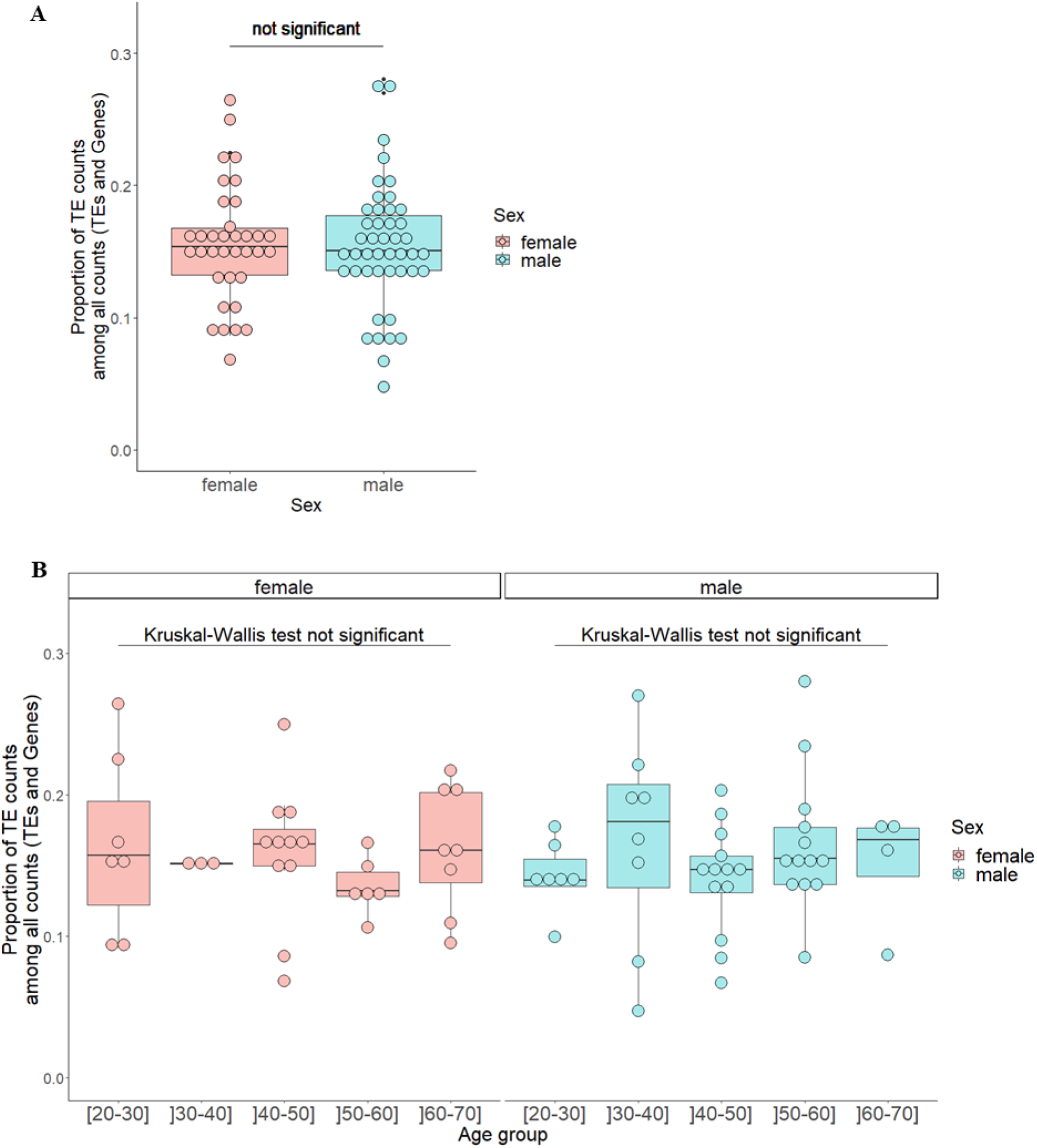
TE expression in the filtered GTEx dataset (no disease group) according to sex (A) regardless of age group or (B) also according to age group. Global TE expression was measured in each karyotype (x-axis) as the proportion of TE read counts among all read counts (TEs and genes) (y-axis). In this calculation, read counts were used after DESeq2 normalization (“normalized counts”) to remove any depth sequencing bias. Each dot represents one individual. Dots are colored according to sex. Sex variable has two levels: female, male. Age group has five levels: [20-30], ]30-40], ]40-50], ]50-60], and ]60-70]. **(A)** Proportion of TE read counts among all read counts was modeled using a linear model with sex and age group as independent variables. From this model, the p-value testing the effect of sex on proportion of TE read counts among all read counts adjusted on age group was not significant (p-value = 0.788). **(B)** A Kruskal-Wallis test comparing proportion of TE read counts among all read counts across age groups was used in females and then in males. This test was not significant in females (p-value = 0.585) as well as in males (p-value = 0.471).

**Figure 4.**
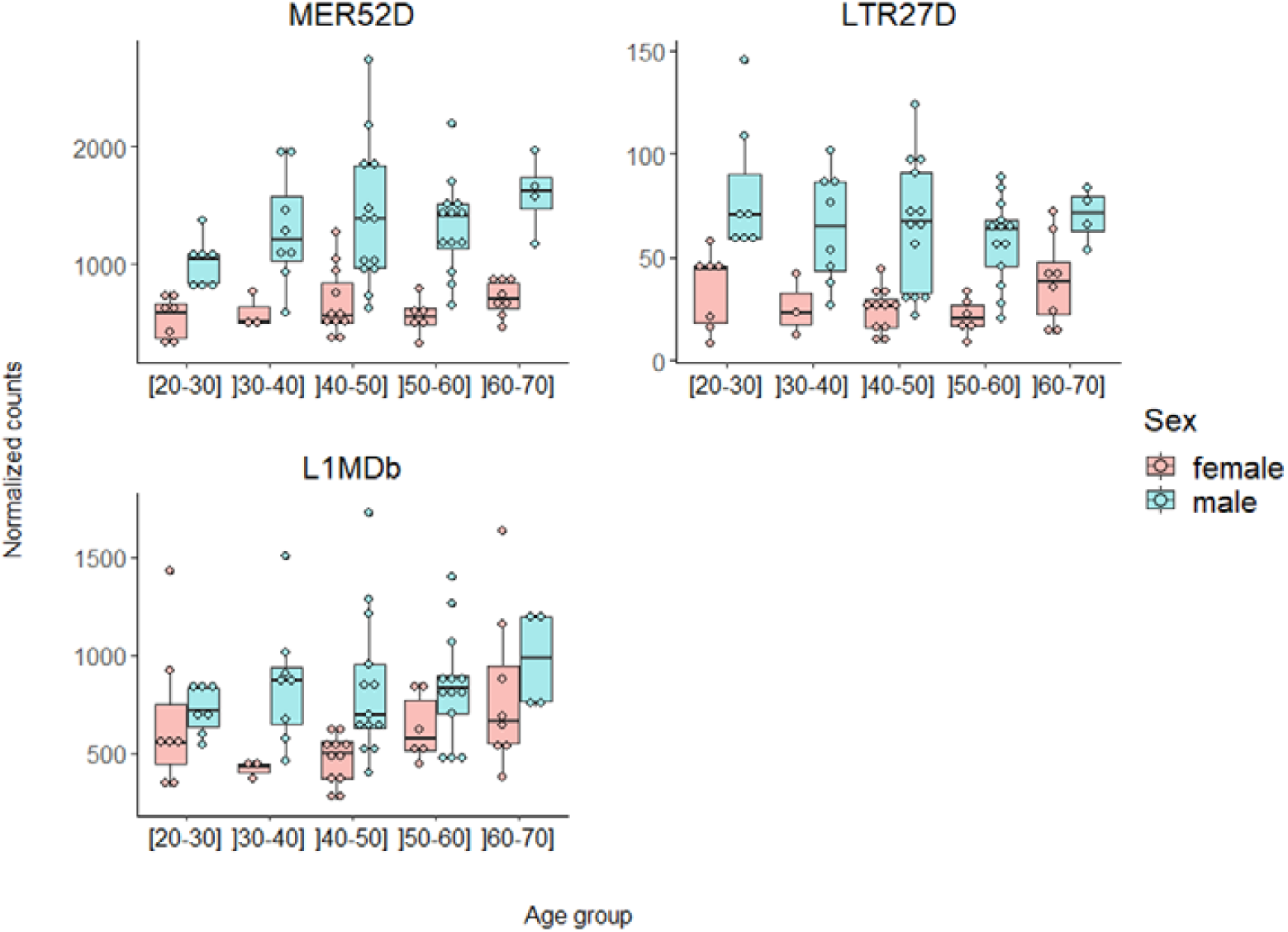
Normalized counts according to sex and age group for the 3 TE subfamilies that are differentially expressed between males and females adjusted on age group in the filtered GTEx dataset (no disease group) Specific TE subfamily expression was estimated using the number of read counts after DESeq2 normalization (“normalized counts”) to remove any depth sequencing bias. All of these 3 TE subfamilies had at least one copy on the Y chromosome and one copy on the X chromosome, but none are enriched on the Y chromosome. For boxplots of normalized counts according to sex and age group for the significantly differentially expressed TE subfamilies according to age group see **Fig. S17.**

Including patients who presented or had presented cancers and/or cardiovascular diseases provided similar results regarding overall TE expression and upregulated TE subfamilies (**Fig. S18 to S20**, **Tables S11 and S12**).

Given that filtering the GTEx dataset is likely to diminish the statistical power of our analysis, we repeated the same analysis without filtering based on the individual’s ethnicity (large GTEx dataset) (**Table S13**). We identified the variable “COHORT” (2 levels: Organ donor (OPO) individuals, Postmortem individuals), referring to the condition in which the organ donation was held, as a potential confounding factor that we have considered in the subsequent analyses. We found no effect of sex on TE expression after adjustment on age group in Organ donor (OPO) individuals nor in Postmortem individuals (**Fig S21A**). Once again, we found no relation between age group and TE transcript amounts in the different levels of the variables COHORT (Organ donor (OPO) individuals, Postmortem individuals) and sex (females, males) (**Fig. S21B**).

Overall, using published transcriptomic data of males and females with normal karyotypes (46,XY for males and 46,XX for females), we did not find a clear association between general TE expression and age. Particularly, we did not find an overall increase of TE expression in old males compared to old females contrary to our initial hypothesis. Nevertheless, we identified several differentially expressed TE subfamilies between 46,XY males and 46,XX females. Additionally, our results showed a higher number of differentially expressed TE subfamilies associated with age in males compared to females.

## Discussion

In this study, we investigated the relationship between TE expression and sex chromosome karyotype composition using transcriptomic data obtained from individuals with different karyotypes (46,XX, 46,XY, 47,XXY and 47,XYY). We observed a significant overall increase in TE transcript levels in 47,XXY males or 47,XYY males compared to 46,XX females, with a near-significant increase observed between 46,XY and 47,XYY males. Additionally, it seemed that the presence of an additional sex chromosome, either X or Y, tended to result in an elevated overall abundance of TE transcripts, likely due to increased genomic material and, consequently, TE load. Notably, consistent with the toxic Y hypothesis, the addition of a Y chromosome appeared to amplify TE expression even more than the addition of an X chromosome.

Our findings suggest a potential association between the presence of the Y chromosome and elevated TE transcripts, aligning with the toxic Y hypothesis. Interestingly, similar observations were found in *D. melanogaster,* where the presence of an additional Y chromosome (in XXY females and in XYY males) was associated with an increased deregulation of Y-linked TE insertions (Brown, Nguyen, and Bachtrog 2020). Moreover, the authors also found that TE silencing mechanisms were compromised even in young flies with a reduction of H3K9me2 histone marks at TE insertions (Brown, Nguyen, and Bachtrog 2020). In those flies, the absence/presence of a Y chromosome correlated with increased/reduced lifespan (Brown, Nguyen, and Bachtrog 2020). Another recent study in the same species proposed that Y chromosomes might affect heterochromatin formation in a size-dependent manner, thereby impacting gene silencing in other chromosomes through the titration of heterochromatin factors (Delanoue et al. 2023). However, when truncating the Y chromosome in different sizes, they did not observe significant differences in male lifespan, suggesting that both the presence and the size of the Y chromosome might not be associated with longevity (Delanoue et al. 2023). Additionally, the authors showed that X0, XY and XYY males displayed identical longevity patterns (Delanoue et al. 2023). In humans, differential TE expression according to sex has already been studied using the GTEx dataset, showing that the number of differentially expressed TE subfamilies between males and females varies significantly across different tissues (Bogu et al. 2019). Nevertheless, in most tissues, including whole blood, differential expression analysis of TE subfamilies between males and females revealed that there was an increased number of upregulated TEs in males compared to downregulated ones (Bogu et al. 2019). The opposite pattern was observed in few tissues such as breast mammary, muscle skeletal, liver, and thyroid (Bogu et al. 2019). Hence, our results with blood samples of 46,XY males and 46,XX females are in agreement with those reported using whole blood GTEx samples, even though different bioinformatic approaches were used.

In several animal species, the association between TE expression and aging has already been studied. It is known that L1 elements are upregulated in aging mouse liver (De Cecco, Criscione, Peterson, et al. 2013) and that several TE families are upregulated in old flies compared to young ones (Brown, Nguyen, and Bachtrog 2020; Li et al. 2013). Specifically, LINE elements were found to be activated in the aging fly brain leading to an age-dependent memory impairment and shortened lifespan (Li et al. 2013). In humans, differential TE expression with respect to age was investigated in the GTEx dataset (Bogu et al. 2019). Considering all tissues, authors found that there were more TE subfamilies that increased expression (679) compared to the ones that decreased expression (63) with age (Bogu et al. 2019). Again, the number of differentially expressed TE subfamilies is strongly dependent on the tissue considered. In whole blood, Bogu et al. found a higher number of TE subfamilies showing a decreased expression with age than an increased expression (Bogu et al. 2019). She et al. drew the same conclusion focusing on differential HERV loci expression according to age in whole blood tissue using GTEx data (She et al. 2022). However, Bogu et al. as well as She et al. did not test the expression of TE subfamilies against interaction between age and sex. Then, and in line with the toxic Y effect, we also explored the association between TE expression and age in males and in females using the GTEx dataset, hypothesizing a greater activation of TEs with aging in males compared to females. However, we could not find any differences in global TE expression levels between old males and old females, or between the oldest individuals and the youngest ones. We suggest that a more homogeneous dataset would be needed to further investigate the relationship between TE expression and age. First, the inclusion of numerous donors with varying conditions and ethnic backgrounds together with factors such as the absence of longitudinal tracking of individuals or survival bias, might introduce biological noise. Secondly, the many different sequencing runs in GTEx data could contribute to technical noise. Furthermore, in humans, it has been recently shown that there is no correlation between retrotransposon expression and chronological age using different human blood tissues (Tsai et al. 2024). However, LTR and LINE expression seem to contribute to biological aging being related with events like cellular senescence, inflammation, and type I interferon response (Tsai et al. 2024).

Regardless, it would be of great interest to explore the association between the Y chromosome, aging, and TE expression in other human tissues using a larger dataset and taking advantage of the recent long read sequencing and specific bioinformatic tools (Schwarz et al. 2022; She et al. 2022; Bogu et al. 2019).

Sex gap in longevity is a multifactorial trait and the toxic Y effect is just one of the possible causes underlying this phenomenon. The present results suggest an association between the Y chromosome and an increase in TE expression. The primary consequence of TE expression as individuals age is the potential for inducing genomic instability through insertional mutagenesis and/or the generation of insertions or deletions. This is facilitated by the occurrence of double-stranded DNA breaks required for TE reinsertion (Levin and Moran 2011; Pray 2008). However, the generation of new somatic mutations with age and the impact of these new TE insertions in the aging process is still to be fully determined in humans (Pabis et al. 2024). Additionally, the sole expression of TEs can also modify in many different ways the expression and structure of genes putatively causing physiological changes that can in turn reduce longevity (Casacuberta and González 2013). Indeed, a recent study in *D. melanogaster* suggested that the number of TE insertions does not increase with age (Schneider et al. 2023). Nonetheless, diminishing the expression of two specific TEs, *412* and *roo*, resulted in an extension of longevity (Schneider et al. 2023). This implies that the expression of TEs, rather than their insertion, play a role in influencing longevity (Schneider et al. 2023). To emphasize the potential impact of TE expression on aging, a recent study demonstrated correlations between LTR and LINE expression in human blood tissues and inflammatory response, while SINE expression was associated with DNA repair processes (Tsai et al. 2024). The latter association suggests a potential link to DNA damage and genome instability (Tsai et al. 2024). Indeed, the detrimental effects of TEs can arise through diverse mechanisms including the accumulation of TE transcripts into the cytoplasm, translation of TE transcripts into proteins, and their interaction of TE transcripts with other genes or proteins (such as the inhibition of tumor suppressor protein). These interactions may activate the immune system, contributing to the “inflammaging” process and potentially promoting cancer development (reviewed in (Mosaddeghi, Farahmandnejad, and Zarshenas 2023)).

In summary, the findings herein are based on a cohort of a small sample size but are also very promising. They open a new window to study the toxic effect of the Y chromosome in human longevity with a particular emphasis on TEs.

## Materials and Methods

### Constitution of the datasets used

#### Gonosome aneuploidy dataset: biological sample collection for 46,XX; 46,XY; 47,XXY; 47,XYY individuals

We obtained blood samples from 24 individuals: six females (46,XX), six males (46,XY), and 12 males with sex chromosome aneuploidies (eight patients with a 47,XXY karyotype and four with a 47,XYY karyotype). All 47,XXY males and 47,XYY males had homogenous 47,XXY or 47,XYY karyotype respectively except for one 47,XXY male (95% of mitoses were XXY, 3% XY, and 2% XXXY) and one 47,XYY male (97% of mitoses were XXY and 3% XY). The median age of the individuals was 30 years old (range: 15 – 63). All individuals signed a consent form for genetic analysis and the study was approved by the ethics committee of the Lyon university hospital (number of the ethics committee: 22_385, number in the register of the *Commission nationale de l’informatique et des libertés*: 22_5385).

Blood samples were collected on a PAXgene tube, kept two hours at room temperature, and then frozen at −80°C until extraction. Blood samples were thawed and then RNA was extracted on a Maxwell RSC system (Promega, Wisconsin, USA) with the Maxwell RSC SimplyRNA Blood kit. Quantity (fluorometric concentration) and quality (RNA Integrity Number, RIN) of the extract were assessed by a fluorometric measurement (Quant-iT RNA Assay Kit, Broad Range from ThermoFisher Scientific, Massachusetts, USA) on a Spark reader (Tecan, Männedorf, Switzerland) and by electrophoresis (TapeStation 4200 System from Agilent Technologies, California, USA), respectively. Each sample was of acceptable RNA quantity (fluorometric concentration of nucleic acids between 26.9 and 109.3 ng/µl) and quality (RIN between 5.8 and 9.3).

Libraries were prepared using the Illumina TruSeq Stranded Total RNA Lp Gold kit (Illumina Inc, California, USA). Size homogeneity of the fragments was verified using the TapeStation 4200 System. Average length of fragments was between 405 and 498 bp (including adapters). Library concentrations were assessed by fluorometric measurement (Quant-iT 1X dsDNA HS Assay Kit from ThermoFisher) on a Spark reader. Concentrations were between 19.3 and 133.0 nmol/L.

Sequencing was performed on a NovaSeq 6000 system (Illumina Inc, California, USA) in paired-end mode (2 x 151 cycles) with a median of 174 million (Min: 132 – Max: 206 million) pairs of reads per sample (after trimming, see below). Demultiplexing was performed with bcl2fastq (v2.20) to obtain FASTQ files. The 24 individuals were divided into two sequencing runs (run 1: 13 individuals, run 2: 11 individuals) (**Table S1**).

#### GTEx datasets

The Genotype-Tissue Expression (GTEx) project regroups samples from a multitude of post-mortem organ donors and from a multitude of tissues. We focused on whole blood tissue to constitute two datasets: one with selection filters to study the most similar individuals as the gonosome aneuploidy dataset generated in this study (one filtered GTEx dataset excluding individuals with current or antecedents of cancer or cardiovascular diseases and one filtered GTEx dataset including them, see below) and one with fewer selection filters (large GTEx dataset).

Filtered GTEx dataset (no disease group): to constitute this dataset, we extracted raw expression data (FASTQ containing Illumina reads) from the human expression atlas database GTEx via the dbGaP portal (dbGaP accession number phs000424.v8.p2). We considered only data of RNA-seq (“Assay.Type” variable) for which a file with sra extension was available (“DATASTORE.filetype” variable) and generated from RNA extracted from whole blood (“body_site” variable). To reduce data heterogeneity and to get closer to the individuals we sampled in parallel to constitute the gonosome aneuploidy dataset generated herein (see above), we focused on: data collected in non-latino nor hispanic white individuals (“RACE” and “ETHNCTY” variables) and excluded data from individuals with persistent infection by the human immunodeficiency virus (HIV), the hepatitis C virus (HCV) or the hepatitis B virus (HBV) (“LBHCV1NT”,“LBHBHCVAB”, “LBHIV1NT”, “LBHIVAB”, “LBHIVO”, and “LBHBSAG” variables), with dementia (“MHALZDMT” and “MHALZHMR” variables), with current or antecedents of cancer (“MHCANCER5”, “MHCANCERC”, and “MHCANCERNM” variables), or cardiovascular diseases (“MHHRTATT”, “MHHRTDIS”, and “MHHRTDISB” variables). At this step, we removed samples identified by the GTEx analysis working group as suboptimal for use in analysis (“SMTORMVE” variable) and we kept only samples for which the material type experiment was described as RNA:Total RNA (“material type exp” variable). Then, we downloaded the remaining FASTQ files (n = 104). For all subsequent analyses, we generated an “age group” variable from the individuals’ age by using five classes of 10-year ranges: [20-30], ]30-40], ]40-50], ]50-60], ]60-70]. Then, using PCA on gene expression or TE expression in the whole dataset obtained and in subsets according to sex or by age group, we identified a strong clustering of individuals who were organ donors OPO (Organ Procurement Organization), who were distinct from individuals who were not (Postmortem individuals) from the “COHORT” variable (**Fig. S22**). We also found that the “COHORT” variable had a statistically significant relation with the age group (Fisher test: p-value < 0.001) but not with the sex (Fisher test: p-value = 0.185). Therefore, we considered the “COHORT” variable as acting as a confounding factor and we excluded the smallest group of individuals, i.e. individuals who were not organ donors OPO (n = 24). Finally, we obtained 80 files suitable for analysis (**Fig. S15**) (see **Table S7** for repartition of samples according to age and sex). The median number of paired-reads (after trimming, see below) was about 44.6 million (Min: 27.0 – Max: 141.1 million) for the 80 FASTQ.

Filtered GTEx dataset (group including individuals with current or antecedents of cancer or cardiovascular diseases): we constituted the same cohort of individuals as before but including 22 individuals with current or antecedents of cancer or cardiovascular diseases (**Table S8**). Therefore, this dataset was composed of 102 individuals. The median number of paired-reads (after trimming, see below) was about 44.3 million (Min: 27.0 – Max: 141.1 million) for the 102 FASTQ.

Large GTEx dataset: to constitute this dataset, we used the 318 curated files containing raw human blood expression data (FASTQ containing Illumina reads) of the Bgee project (see supplementary data of (Bastian et al. 2021)) originating from dbGaP (dbGaP accession number phs000424.v6.p1). We excluded one male suspected of having sex chromosome aneuploidy (such as 47,XXY) because of the expression of *XIST* like individuals in possession of two X chromosomes and the expression of *USP9Y* like individuals in possession of one Y chromosome. We generated an “age group” variable from the individuals’ age by using five classes of 10-year ranges: [20-30], ]30-40], ]40-50], ]50-60], ]60-70] and we found a significant statistical relation between sex and age group variables (Pearson’s Chi-squared test: p-value = 0.013) (**Table S13**). Again, using PCA on gene expression or TE expression, we identified a strong clustering of individuals who were organ donors OPO (Organ Procurement Organization), who were distinct from individuals who were not (Postmortem and Surgical individuals) from the “COHORT” variable. We also found that the “COHORT” variable had a significant statistical relation with the age group (Pearson’s Chi-squared test: p-value < 0.001, excluding the 3 individuals with “COHORT” = Surgical) and with the sex group (Pearson’s Chi-squared test: p-value = 0.017, excluding the 3 individuals with “COHORT” = Surgical). Then we considered the variable “COHORT” as a potential confounding factor and added it in the DESeq2 statistical model (see below). Therefore, we excluded 3 more individuals for whom “COHORT = Surgical” to guarantee a sufficient number of individuals in each COHORT, sex, and age group levels. Thus, 314 files were used in downstream analysis.

### Bioinformatic analysis

See pipeline in **Fig. S1**.

#### Input files

FASTQ files were used after a quality control step with FastQC (v0.11.9) and Trimmomatic (v0.33) softwares (Bolger, Lohse, and Usadel 2014). The whole human reference transcriptome (Human Release 32 GRCh38/hg38, gencode v32) was downloaded from the University of California Santa Cruz (UCSC) table browser (https://genome.ucsc.edu/cgi-bin/hgTables?hgsid=814613547_qUmIEGan4K2f39cHvb9AgrKdRVdK&clade=mammal&org=Human&db=hg38&hgta_group=genes&hgta_track=knownGene&hgta_table=0&hgta_regionType=genome&position=chr1%3A11%2C102%2C837-11%2C267%2C747&hgta_outputType=primaryTable&hgta_outFileName=) with output format = “sequence” (fasta file) and used for the differential gene expression analysis. The downloaded file contained 247,541 transcript sequences (5’ and 3’ untranslated regions and coding DNA sequences of each gene were included, introns were discarded, and repeated regions were masked). The GTF file corresponding to release 32 GRCh38/hg38 of the human reference genome was downloaded from the gencodegenes website https://www.gencodegenes.org/human/ (content = “comprehensive gene annotation”, regions = “CHR”). This file provided a correspondence between transcripts (227,463 transcript names) and genes (60,609 genes, including 19,947 protein coding genes). The human reference sequence file (including TE insertion sequences) was downloaded as a fasta from the UCSC table browser (https://genome.ucsc.edu/cgi-bin/hgTables?hgsid=791366369_Zf0cNT7ykVM0ZQ0zErZRArSgvMEO&clade=mammal&org=Human&db=hg38&hgta_group=allTracks&hgta_track=rmsk&hgta_table=0&hgta_regionType=genome&position=chr1%3A11%2C102%2C837-11%2C267%2C747&hgta_outputType=primaryTable&hgta_outFileName=) including the RepeatMasker track from the “Dec. 2013 GRCh38/hg38” assembly of the human reference genome. This file contains sequences of 4,886,205 insertion copies, after excluding transfer RNAs, ribosomal RNAs, small RNAs, repeats U1 to U13, and satellites (microsatellites, GSATX, HSAT5). From this file, we built the “rosetta” file (**Data S6**), which provided a correspondence between the 4,886,205 insertion sequences and insertion subfamily names (1,270 repeat subfamilies). Since we did not have access to the genome of the participants, we could not determine the location of each repeat copy in each individual, which may vary from one to another. Therefore, the rosetta file allowed us to regroup read counts per repeat copy into read counts per repeat subfamily, independently of their location in the genome.

#### Alignment and read counting

Gene expression was quantified with Kallisto software (v0.46.1) (Bray et al. 2016) using the transcript sequences from the whole human reference transcriptome. The --rf-stranded option was added to deal with reverse stranded reads when analyzing RNAseq data of the gonosome aneuploidy dataset (contrary to GTEx datasets as GTEx data are non-stranded). Alignments to the reference transcriptome resulted in high mapping efficiency (**Table S1**). Repeat expression (alignment and read count) was analyzed with the TEcount (v1.0.0) software from TEtools (Lerat et al. 2016). This tool uses Bowtie2 (Langmead and Salzberg 2012) as aligner, and we added the --nofw option to deal with reverse stranded RNAseq data when analyzing RNAseq data of the gonosome aneuploidy dataset (contrary to GTEx datasets).

### Statistical analysis

Statistical analysis was performed using the R software (v3.6.3 to 4.3.2). The significance threshold was set at 0.05 for all statistical tests performed. The GTEx datasets and the gonosome aneuploidy dataset were computed separately using the same procedure described below. We considered as gene or TE “expression” the number of reads aligned (i.e. counts) on each reference transcript or TE sequence and as “differential expression analysis” the statistical procedure to compare gene or TE expression between several conditions (i.e. sex, age group, or karyotype).

#### Data preparation

Differential expression analysis was performed separately for genes and for TEs. Two files were prepared for each purpose. Since one gene can have several transcripts, the tximport R package (v1.14.2, Bioconductor) was used to sum the read counts per gene transcript into read counts per gene using the GTF file mentioned above. Then, a txi file containing the gene identifier (Ensembl nomenclature), the average gene length, and the estimated counts for all genes per sample was obtained. For DE analysis on TEs, we normalized the repeat subfamily read counts based on the normalization factor previously calculated using gene read counts to avoid masking relevant biological information (Lerat et al. 2016). To estimate the normalization factor, the gene identifiers and read counts per gene were extracted from the txi file and merged with the TEcount file containing the repeat subfamily names and read counts per repeat subfamily. The length could not be included in the merged table since this variable was not available from the TEcount table (each repeat subfamily is a collection of copies displaying a diversity of lengths, it is thus difficult to estimate the length of a TE subfamily). This merged table was then imported into the DESeq2 tool (R package DESeq2 v1.26.0, Bioconductor) (Anders and Huber 2010; Love, Huber, and Anders 2014), using the DESeqDataSetFromMatrix function, which normalized the read counts so as to remove any depth sequencing bias. From this step on, we removed read counts on the following 24 snRNA, snpRNA, scRNA, or remaining satellites and rRNA repeat subfamilies: 7SLRNA, 7SK, LSAU, D20S16, REP522, SATR1, SATR2, ACRO1, ALR/Alpha, BSR/Beta, CER, 6kbHsap, TAR1, SST1, MSR1, SAR, GA-rich, G-rich, A-rich, HY1, HY3, HY4, HY5,5S. This step allowed us to only consider the 1,246 subfamilies corresponding to TE subfamilies out of the 1,270 repeat subfamilies contained in the rosetta file. Then, genes and TE subfamilies with zero read counts in all individuals were removed leaving 1,212 TE subfamilies to study.

For the DE analysis on genes, as the length was available for genes, a more reliable procedure was performed. The raw txi file was directly imported in DESeq2 (DESeqDataSetFromTximport function) and the average length of each gene could be used as a correction term (offset) in the DESeq2 statistical model described below to consider the length bias (Anders and Huber 2010; Love, Huber, and Anders 2014). This procedure normalized the read counts in order to remove any depth sequencing bias and also bias related to sequence length.

#### Comparison of the global expression of TE between sex, age group or karyotypes

We first looked at the global TE expression rather than the TE subfamily expression separately. To do that, using the merge table imported into DESeq2, we summed all the normalized counts of each TE subfamily per individual and then we divided by the total number of normalized counts on genes and TE subfamilies per individual. A pairwise Wilcoxon test was used to compare global TE expression between karyotypes in the gonosome aneuploidy dataset. In GTEx datasets, linear models or Wilcoxon tests were used, as appropriate, to compare global TE expression between females and males (adjusted on age group and COHORT variables when necessary), and between the youngest individuals ([20-50] years, obtained by combining individuals from [20-30], ]30-40], and ]40-50] age groups) and the oldest individuals (]50-70] years, obtained by combining individuals from ]50-60], and ]60-70] age groups). Kruskall-wallis tests or ANOVA were also used, as appropriate, to test the relation between global TE expression and age group in the GTEx datasets in males and females separately. The same procedure was made to looked at expression of TE classes/subclasses (LTR-retrotransposons, non-LTR retrotransposons: SVA, non-LTR retrotransposons: SINE, non-LTR retrotransposons: LINE, DNA transposons) or some TE groups (SVA, HERVK, AluS, L1, AluY) by summing only normalized counts of TE subfamilies belonging to this TE class/subclass or this TE group according to the “class/family” column of the hg38.fa.out.gz file downloaded from https://www.repeatmasker.org/species/hg.html (track hg38 RepeatMasker open-4.0.6 - Dfam 2.0) or according to the name of TE subfamilies when necessary.

#### Differential expression analysis on genes and TE subfamilies

Then, we performed a differential expression analysis for each gene and each TE subfamily separately.

The R package DESeq2 (v1.26.0, Bioconductor) (Anders and Huber 2010; Love, Huber, and Anders 2014) was used to fit a negative binomial model on the log2 of the normalized counts (dependent variable) according to two independent variables: sex (factor with 2 levels: female, male) and age group (factor with 5 levels: [20,30], ]30,40], ]40,50], ]50,60], ]60,70]) and the interaction of the two for the GTEx datasets, or karyotype (factor with 4 levels: XX, XY, XXY, XYY) and sequencing batch (factor with 2 levels: first batch, second batch) for the gonosome aneuploidy dataset. The statistical model provided variation of expression of a gene or a TE subfamily between two conditions expressed as log2 fold-change (log2FC) (i.e. log base 2 of the ratio of predicted normalized counts between the two conditions). Log2FC were compared to 0 with a Wald test. A likelihood-ratio test (LR test) was also performed to assess whether adding an independent variable in the model provided relevant information compared to the null model, i.e. to test whether gene or TE subfamily expression was globally impacted by an independent variable having more than two levels (age group or karyotype variables). A 0.05 False-Discovery Rate (FDR) threshold value was used for significance and p-values were adjusted for multiple-testing using the Benjamini & Hochberg procedure (Benjamini and Hochberg 1995). The thresholds for the outlier and the low mean normalized read count filters were set as default. Graphics were generated after the data transformations proposed by the DESeq2 package authors: variance stabilizing transformation for PCA, adaptive shrinkage for Minus-Average plots (MA plots), log2 transformation of the normalized counts for heatmaps (Anders and Huber 2010; Stephens 2016). When specified, the command removeBatchEffect function (for unbalanced batches) from the limma R package (v3.46.0) was used before PCA representations, heatmap representations and before building the different boxplots of proportion of TE read counts or normalized counts to deal with batch effect in the gonosome aneuploidy dataset or COHORT effect in the large GTEx dataset. When necessary, gene name, gene description, and gene localization were determined using the GTF file and the command gconvert from the R package gprofiler2 (v0.1.9) (Raudvere et al. 2019; Kolberg et al. 2020) from the Ensembl gene (ENSG) identifier.

#### Overlap between the DEGs generated in our study and in published studies

The DEGs generated in our study in the XXY karyotype compared to the XY karyotype was compared to those found in the supplementary materials of Zhang et al. study (Zhang et al. 2020) and to confirmed XCI-escapees in supplementary materials of Wainer-Katsir et al. study (Wainer Katsir and Linial 2019).

#### GO term enrichment analysis

A gene ontology (GO) analysis was performed using the lists of significantly upregulated genes (adjusted p-value < 0.05 and log2-fold change > 0) and significantly downregulated genes (adjusted p-value < 0.05 and log2-fold change < 0) obtained for each of the six pairwise karyotype comparisons (47,XYY vs 46,XX, 47,XXY, 46,XX, 46,XY vs 46,XX, 47,XYY vs 46,XY, 47,XXY vs 46,XY, 47,XYY vs 47,XXY) after adjustment on the batch effect. To generate lists of significant GO terms (**Data S3**), we used the gost function from the R package gprofiler2 (v0.1.9) (Raudvere et al. 2019; Kolberg et al. 2020) with the g:SCS algorithm for multiple testing correction and a significance threshold set at 0.05. Other parameters were set to default. To generate plots (**Fig. S5 to S7**), we used the compareCluster function from the R package clusterProfiler (v4.10.0) (Wu et al. 2021) with the Benjamini & Hochberg procedure (Benjamini and Hochberg 1995) for multiple testing correction and a significance threshold set at 0.05. Other parameters were set to default. We generated a plot concerning terms in the GO database, KEGG database and Reactome database. Then, we plotted specifically the significant terms from the Reactome database including the word “senescence” in their description (**Fig. S8**).

#### Y enrichment of TE copies

For all of the 1,246 TE subfamilies, an exact Binomial test was performed to test if the observed proportion of TE copies for one TE subfamily carried by the Y chromosome was equal to the expected proportion of 1.85%. The expected proportion of 1.85% (57200/3088200) was calculated according to the length of Y chromosome (57200 kb) compared to the sum of the length of all chromosomes (3088200 kb) (“Human Genome Assembly GRCh38.P14,” n.d.). The observed proportion of TE copies corresponded to the ratio between the observed number of TE copies from a TE subfamily carried by the Y chromosome and the observed total number of TE copies from the same TE subfamily in the human reference sequence file including TE insertion sequences (see above). The expected number of TE copies from a TE subfamily carried by the Y chromosome was determined as the total number of TE copies multiplied by 1.85%. The exact Binomial test was two-sided. A 0.05 False-Discovery Rate (FDR) threshold value was used for significance and p-values were adjusted for multiple-testing using the Benjamini & Hochberg procedure. **Data S4** regroups the observed and expected numbers of TE copy carried by the Y or the X chromosome and the adjusted p-values of the exact Binomial test obtained for each of the 1246 TE subfamilies. Then, using **Data S4**, we made 6 groups of TE subfamilies: one concerning Y-enriched TE subfamilies (binomial test adjusted p-value < 0.05 and number of observed copies located on Y chromosome > expected), one concerning Y-depleted TE subfamilies (binomial test adjusted p-value < 0.05 and number of observed copies located on Y chromosome < expected), one concerning neither Y-enriched nor Y-depleted TE subfamilies (TE subfamilies that did not meet the conditions for Y-enriched nor Y-depleted TE subfamilies), one concerning TE subfamilies with no copies carried by the Y chromosome, one concerning TE subfamilies with no copies carried by the X chromosome, and one concerning TE subfamilies with no copies carried by either the Y or X chromosomes. Finally, a pairwise Wilcoxon test was used to compare TE expression between karyotypes in these 6 TE groups in the gonosome aneuploidy dataset. In GTEx datasets, linear model was used to compare TE expression between females and males, adjusted on age group, and kruskall-wallis test was used to test the relation betweenTE expression and age group in males and females separately.

#### Overlap between 3 kb upstream regions of differentially expressed genes and differentially expressed TE subfamilies

First, we downloaded a bed file for repeats from https://hgdownload.soe.ucsc.edu/goldenPath/hg38/bigZips/ (hg38.fa.out.gz file, last update 2014-01-15 20:56) and we generated a bed file for genes from the GTF file Release 32 GRCh38/hg38 previously downloaded. From the gene bed file, we generated a bed file containing the 3 kb upstream region of all genes. Then, we filtered this bed file using the list of differentially upregulated (downregulated, respectively) genes obtained for each pairwise karyotype comparison. We filtered the repeat bed file using the list of differentially upregulated (downregulated, respectively) TE subfamilies obtained for each pairwise karyotype comparison. Then, we used the R package RegioneR v1.34.0 (Gel et al. 2016) (toGRanges and numOverlaps functions) to obtain the number of TE copies whose sequence overlaps the 3 kb upstream sequence of any gene using the filtered bed files previously created. A permutation test was performed to test whether the frequency of genes with TE copies overlapping their 3 kb upstream region was significantly greater or lower than expected by chance. We used the permTest function with the following options:ntimes=5000, randomize.function=resampleRegions, evaluate.function=numOverlaps, count.once=TRUE, alternative = “auto”, and the 3 kb upstream region of all genes as universe. We made this procedure four times for each pairwise karyotype comparisons: one considering TE copies from upregulated TE subfamilies and upregulated genes, one considering TE copies from upregulated TE subfamilies and downregulated genes, one considering TE copies from downregulated TE subfamilies and upregulated genes, and one considering TE copies from downregulated TE subfamilies and downregulated genes.

#### Overlap between intragenic regions of differentially expressed genes and differentially expressed TE subfamilies

We also looked for overlap between TE copy sequences of upregulated TE subfamilies and sequences of upregulated genes using bedtools intersect (v2.30.0) with -s option to consider the strandness. Then, we made the same procedure considering TE copy sequences of downregulated TE subfamilies and sequences of downregulated genes. Then, we performed an exact binomial test for each pairwise karyotype comparison which compared the observed proportion of upregulated genes containing at least one TE copy from one of the upregulated TE subfamilies among all the upregulated genes to the expected proportion of genes containing at least one TE copy from one of the upregulated TE subfamilies among all genes. The same procedure was also applied considering then downregulated genes and downregulated TE subfamilies.

## Supporting information

DataS1

DataS2

DataS3

DataS4

DataS5

DataS6

## Appendices

Supplementary material including supplementary Fig. S1 to S22 and supplementary Tables S1 to S13.

Data S1. List of all differentially expressed genes on X chromosome or Y chromosome according to karyotype using the likelihood-ratio test in the gonosome aneuploidy dataset, first considering all genes and then considering protein-coding genes only.

Data S2. List of all differentially expressed genes (regardless if they are protein-coding or not) in each pairwise karyotype comparison after batch adjustment in the gonosome aneuploidy dataset.

Data S3. GO analysis in the gonosome aneuploidy dataset. Data S4. Y chromosome enrichment test for TE subfamilies.

Data S5. Overlap between genomic positions of upregulated (downregulated, respectively) genes and positions of upregulated (downregulated, respectively) TEs for each pairwise karyotype comparison (when applicable).

Data S6. Rosetta file.

## Acknowledgements

We thank Yasmine Zerdoumi and Maxime Vallée for their help in the data generation and data analysis, and Véréna Landel (DRS, Hospices Civils de Lyon) for reading the manuscript. We thank Marc Robinson-Rechavi for access to the Bgee database and useful comments on the GTEx analysis and on the manuscript. Part of this work was performed using the computing facilities of the CC LBBE/PRABI-AMSB. The Genotype-Tissue Expression (GTEx) Project was supported by the Common Fund of the Office of the Director of the National Institutes of Health, and by NCI, NHGRI, NHLBI, NIDA, NIMH, and NINDS. Filtered GTEx datasets were generated as part of dbGaP Project #28282: “Studying transposable elements activity in human tissues”. We gratefully thank the GTEx group for the data provided.

A preprint version of this article has been peer-reviewed and recommended by PCI Genomics (https://doi.org/10.24072/pci.genomics.100293).

## Data, scripts, code, and supplementary information availability

Fastq files of the gonosome aneuploidy dataset generated in this study were submitted to the European Genome-phenome Archive (EGA ID of the study: EGAS00001007462).

Scripts and code were described in detail in the Materials and methods section. The code of the pipeline used to generate gene and TE count files from FastQ files is available here: https://github.com/teolijo/longevitY/blob/main/pipeline. More details can be supplied upon request.

## Authors’ contributions

Conceptualization: GABM, IP, CV; Methodology: MF, DS, CV, GABM, IP; Software: JT, MF, AN, DS-O, AB-C, CB; Formal analysis: JT, DS-O; Investigation: JT, AN, AL, HL, DS, CB, GABM, IP; Resources: JT, MF, AN, AB-C, AL, HL, DS, CB, GABM, IP; Data Curation: JT, MF, AN; Writing - Original Draft: JT, GABM; Writing - Review & Editing: JT, MM and all authors; Visualization: JT, MM, CV, GABM, IP; Supervision: IP, GABM, CV; Project administration: IP, GABM, CV; Funding acquisition: GABM, IP, CV, FG, DS.

