## Supplementary materials for "Transposable element expression with variation in sex chromosome number: insights into a toxic Y effect on human longevity"

**This PDF file includes:**

Supplementary text

Figs. S1 to S22

Tables S1 to S13

**Other Supplementary Materials for this manuscript include the following:**

Data S1 to S6

#### Supplementary text

The differential expression study on genes found expected differentially expressed genes like those located on the Y chromosome to validate our protocol. Furthermore, we found *XIST* and *TSIX* genes to be expressed only in karyotypes containing two X chromosomes (47,XXY and 46,XX), which was also expected as these genes are implicated in the X inactivation occurring only in the presence of two or more X chromosomes (Loda and Heard 2019; Hassan et al. 2019; Skakkebæk et al. 2018). In the karyotype with two or more X chromosomes (normal 46,XX karyotype or abnormal karyotype such as 47,XXY, 47,XXX…), the extra-X chromosomes are inactivated and that; only the X-linked genes escaping inactivation exhibit expression changes according to X chromosome dosage (Monkhorst et al. 2008; Zhang et al. 2020). We found 6 of the 31 (19.4%) confirmed XCI escapees in the study of Wainer-Katsir et al. (Wainer Katsir and Linial 2019) among the upregulated genes retrieved when in 47,XXY karyotype was compared to 46,XY karyotype. We also found differentially expressed genes when we compared 47,XXY to 46,XY or 46,XX karyotypes that were described elsewhere. Indeed, we shared 22 of the 130 (16.9%) upregulated genes (regardless if they are protein-coding or not) in 47,XXY compared to 46,XY reported in the study of Zhang et al. and 5 of the 111 (4.5%) downregulated genes in 47,XXY compared to 46,XY (Zhang et al. 2020).

#### Supplementary figures

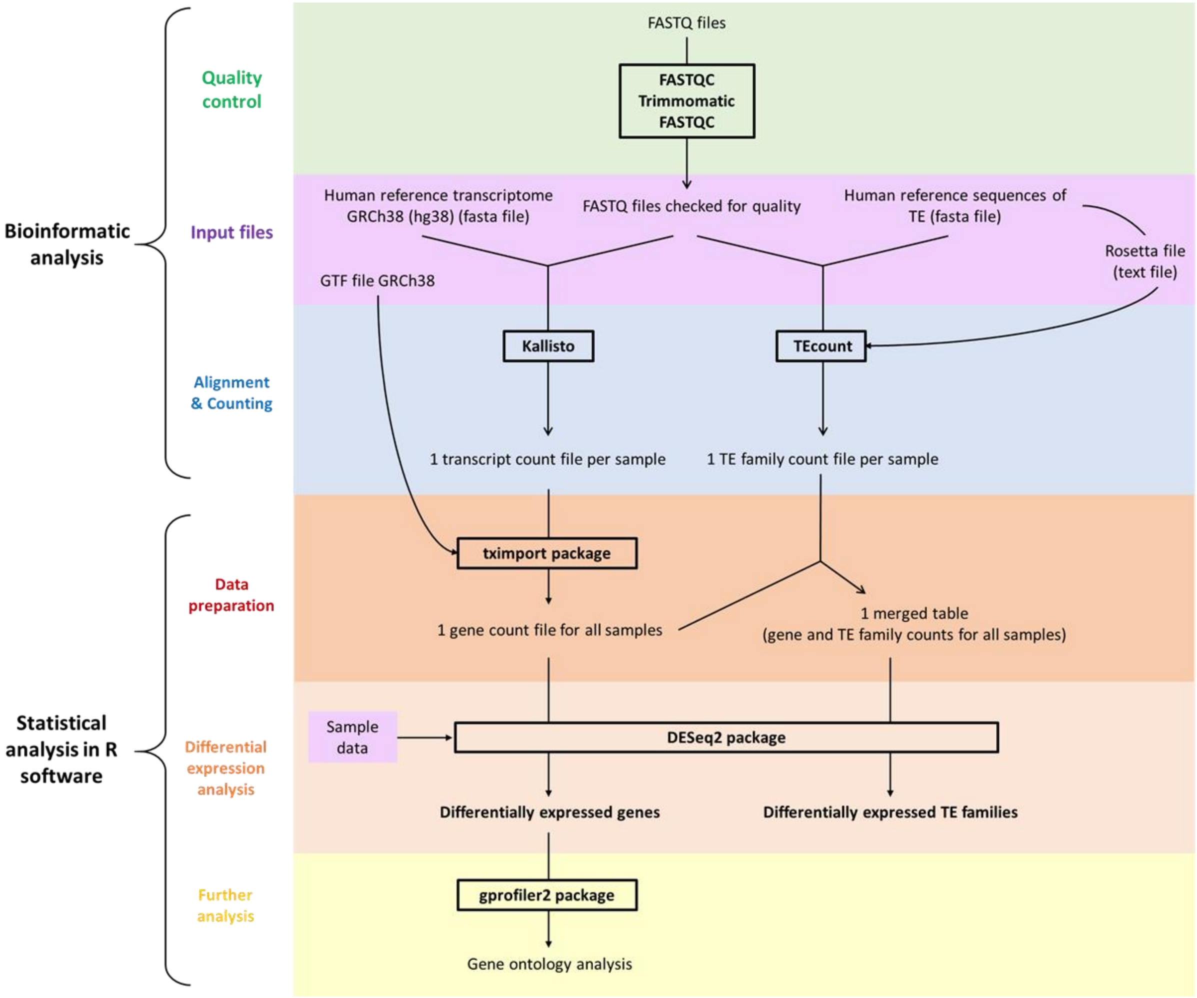


**Fig. S1. Pipeline for gene and TE differential expression analysis.**

GTF file: Gene transfer format file, TE: Transposable element.

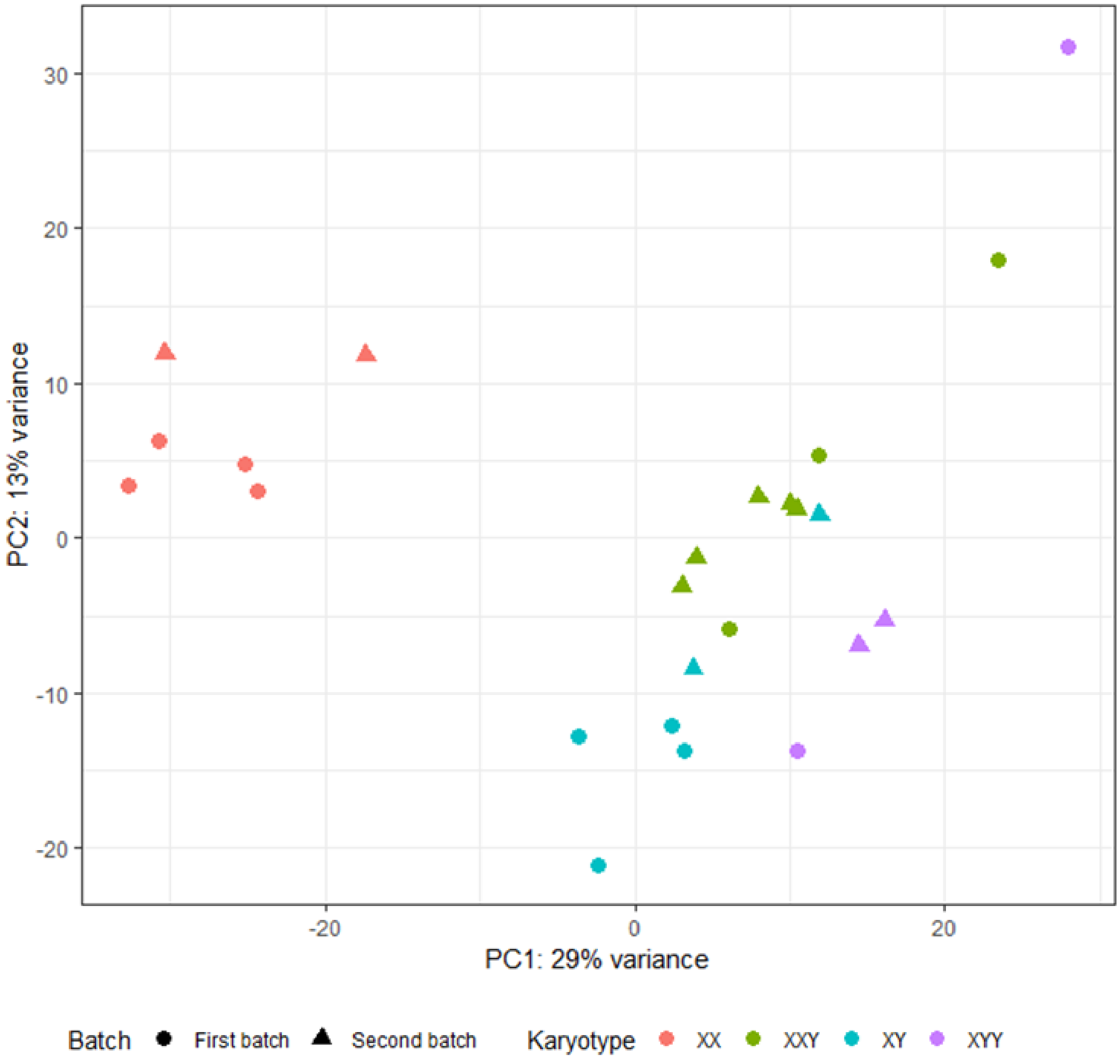


**Fig. S2. Principal component analysis on the 500 genes explaining the most variance in the gonosome aneuploidy dataset after batch effect was removed.**

Each dot represents one individual. Dots are colored according to karyotype (XX, XY, XXY, XYY) and shaped according to batch (first batch, second batch). Batch effect was removed before graphical display. PC: Principal Component.

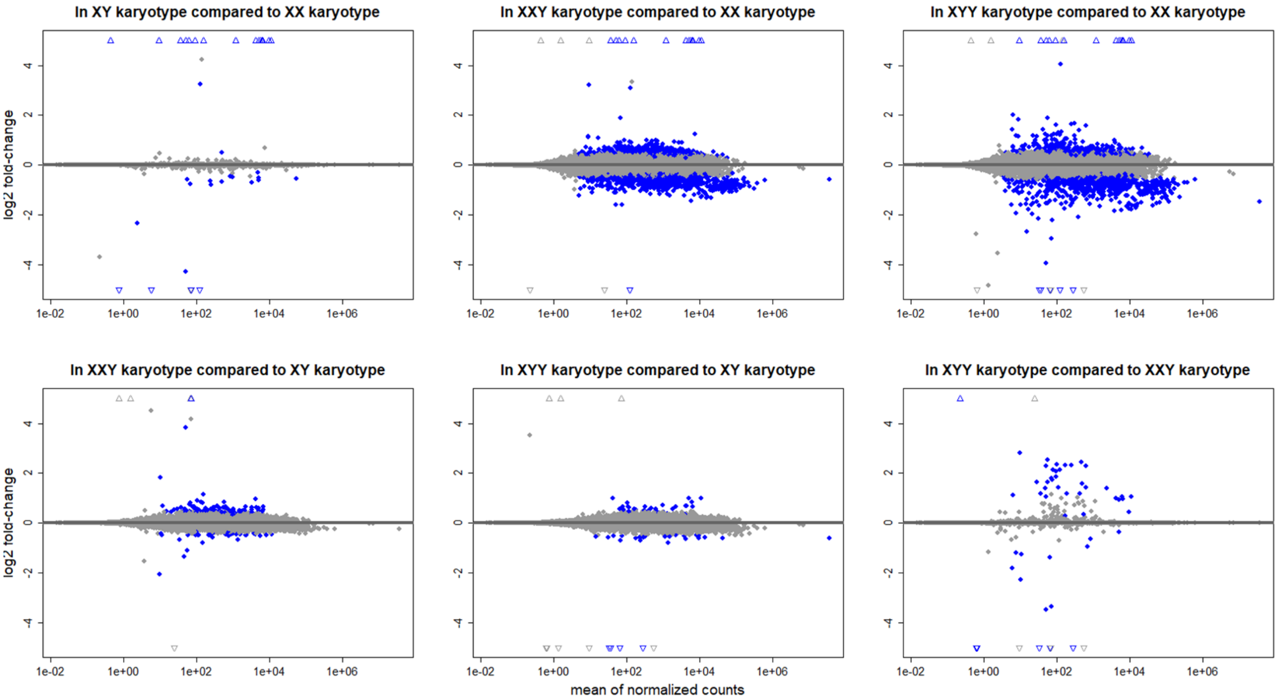


**Fig. S3. MA plots on protein-coding genes in each pairwise karyotype comparison after batch adjustment in the gonosome aneuploidy dataset.**

Minus-Average plots (MA plots) represent, for each protein-coding gene, the $\log_2$(normalized counts in group 1) – $\log_2$(normalized counts in group 2) on the y-axis and the mean expression (i.e. mean of normalized counts) across all the samples on the x-axis. Each dot or triangle represents one protein-coding gene. Blue dots show genes with an adjusted p-value $< 0.05$. Genes for which log2 fold-change is greater than 4 or less than -4 are represented by an empty triangle. Positive log2 fold-change means that the gene is upregulated in the first karyotype (e.g. XYY in the “XYY karyotype vs XX karyotype” comparison) compared to the second (e.g. XX in the “XYY karyotype vs XX karyotype” comparison). Negative log2 fold-change means that the gene is downregulated in the first karyotype (e.g. XYY in the “XYY karyotype vs XX karyotype” comparison) compared to the second (e.g. XX in the “XYY karyotype vs XX karyotype” comparison).

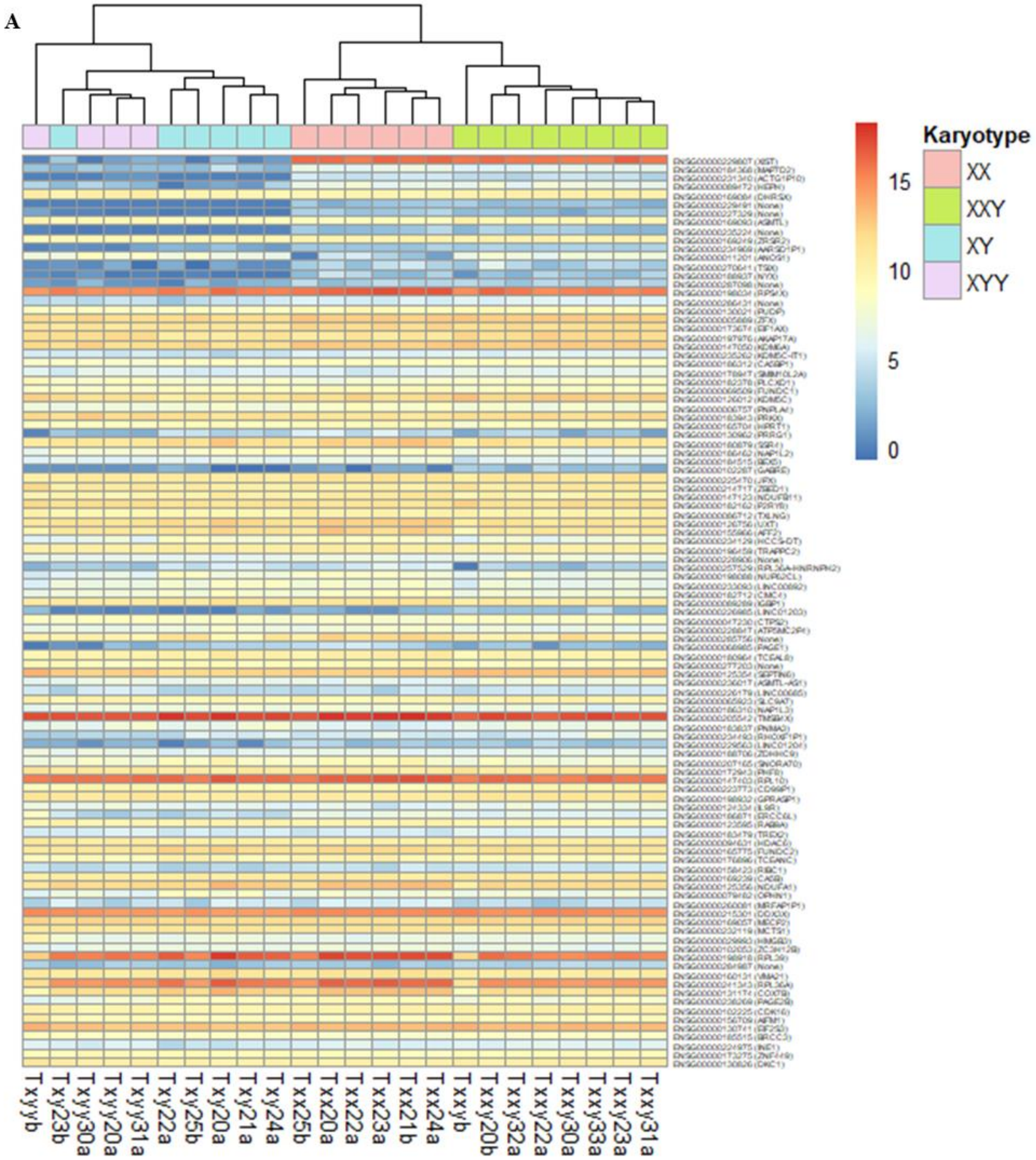
A
Karyotype
XX
XXY
XY
XYY
15
10
5
0
Txyyb
Txy23b
Txyy30a
Txyy20a
Txyy31a
Txy22a
Txy25b
Txy20a
Txy21a
Txy24a
Txx25b
Txx20a
Txx22a
Txx23a
Txx21b
Txx24a
Txxyb
Txxy20b
Txxy32a
Txxy22a
Txxy30a
Txxy33a
Txxy23a
Txxy31a

**B**

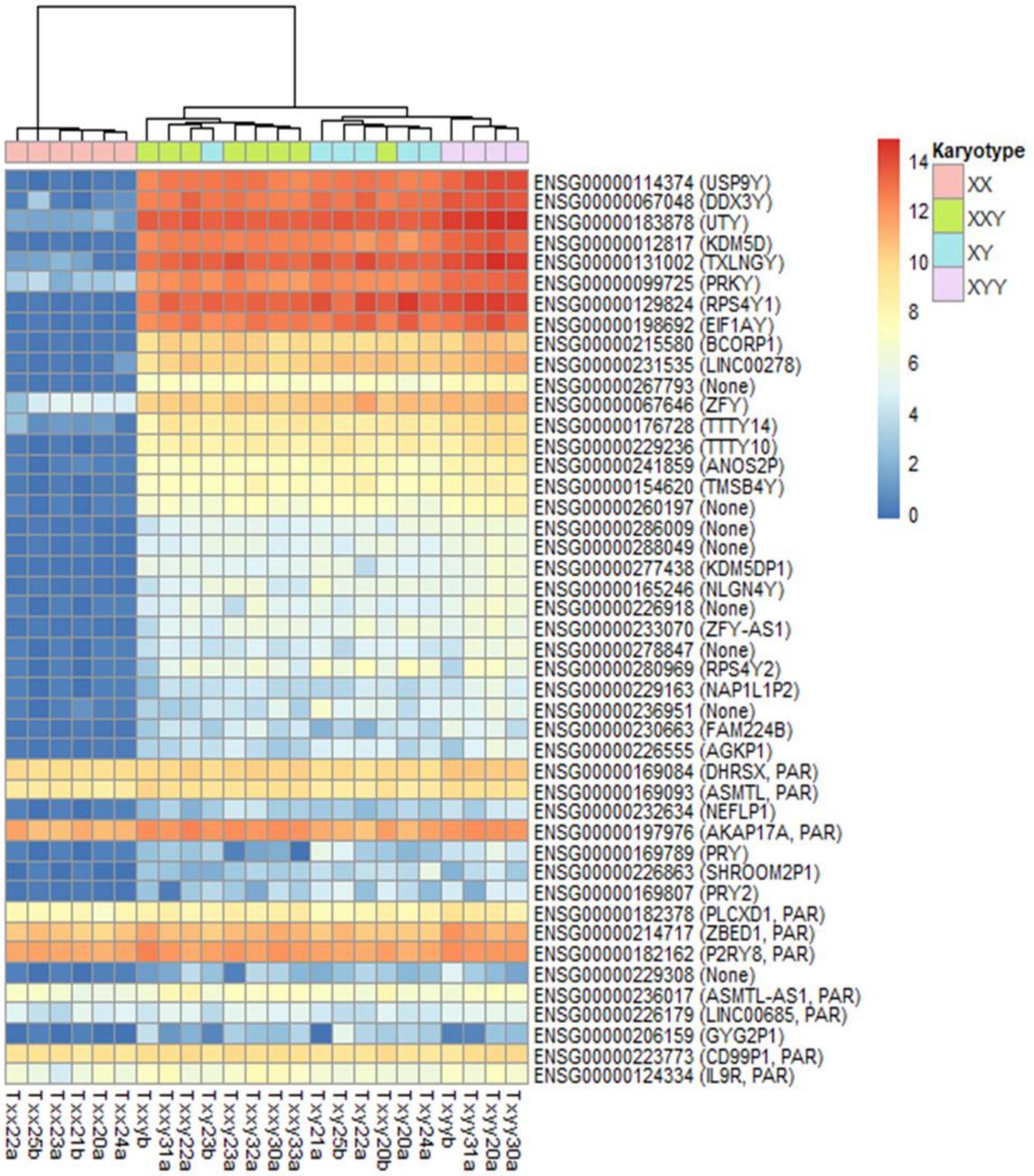
Karyotype
XX
XXY
XY
XYY
14
12
10
8
6
4
2
0
ENSG00000114374 (USP9Y)
ENSG00000067048 (DDX3Y)
ENSG00000183878 (UTY)
ENSG00000012817 (KDM5D)
ENSG00000131002 (TXLNGY)
ENSG00000099725 (PRKY)
ENSG00000129824 (RPS4Y1)
ENSG00000198692 (EIF1AY)
ENSG00000215580 (BCORP1)
ENSG00000231535 (LINC00278)
ENSG00000267793 (None)
ENSG00000067646 (ZFY)
ENSG00000176728 (TTTY14)
ENSG00000229236 (TTTY10)
ENSG00000241859 (ANOS2P)
ENSG00000154620 (TMSB4Y)
ENSG00000260197 (None)
ENSG00000286009 (None)
ENSG00000288049 (None)
ENSG00000277438 (KDM5DP1)
ENSG00000165246 (NLGN4Y)
ENSG00000226918 (None)
ENSG00000233070 (ZFY-AS1)
ENSG00000278847 (None)
ENSG00000280969 (RPS4Y2)
ENSG00000229163 (NAP1L1P2)
ENSG00000236951 (None)
ENSG00000230663 (FAM224B)
ENSG00000226555 (AGKP1)
ENSG00000169084 (DHRSX, PAR)
ENSG00000169093 (ASMTL, PAR)
ENSG00000232634 (NEFLP1)
ENSG00000197976 (AKAP17A, PAR)
ENSG00000169789 (PRY)
ENSG00000226863 (SHROOM2P1)
ENSG00000169807 (PRY2)
ENSG00000182378 (PLCXD1, PAR)
ENSG00000214717 (ZBED1, PAR)
ENSG00000182162 (P2RY8, PAR)
ENSG00000229308 (None)
ENSG00000236017 (ASMTL-AS1, PAR)
ENSG00000226179 (LINC00685, PAR)
ENSG00000206159 (GYG2P1)
ENSG00000223773 (CD99P1, PAR)
ENSG00000124334 (IL9R, PAR)
Txx22a
Txx25b
Txx23a
Txx21b
Txx20a
Txx24a
Txxyb
Txxy31a
Txxy22a
Txy23b
Txxy23a
Txxy32a
Txxy30a
Txxy33a
Txy21a
Txy25b
Txy22a
Txxy20b
Txy20a
Txy24a
Txyyb
Txyy31a
Txyy20a
Txyy30a

**C**

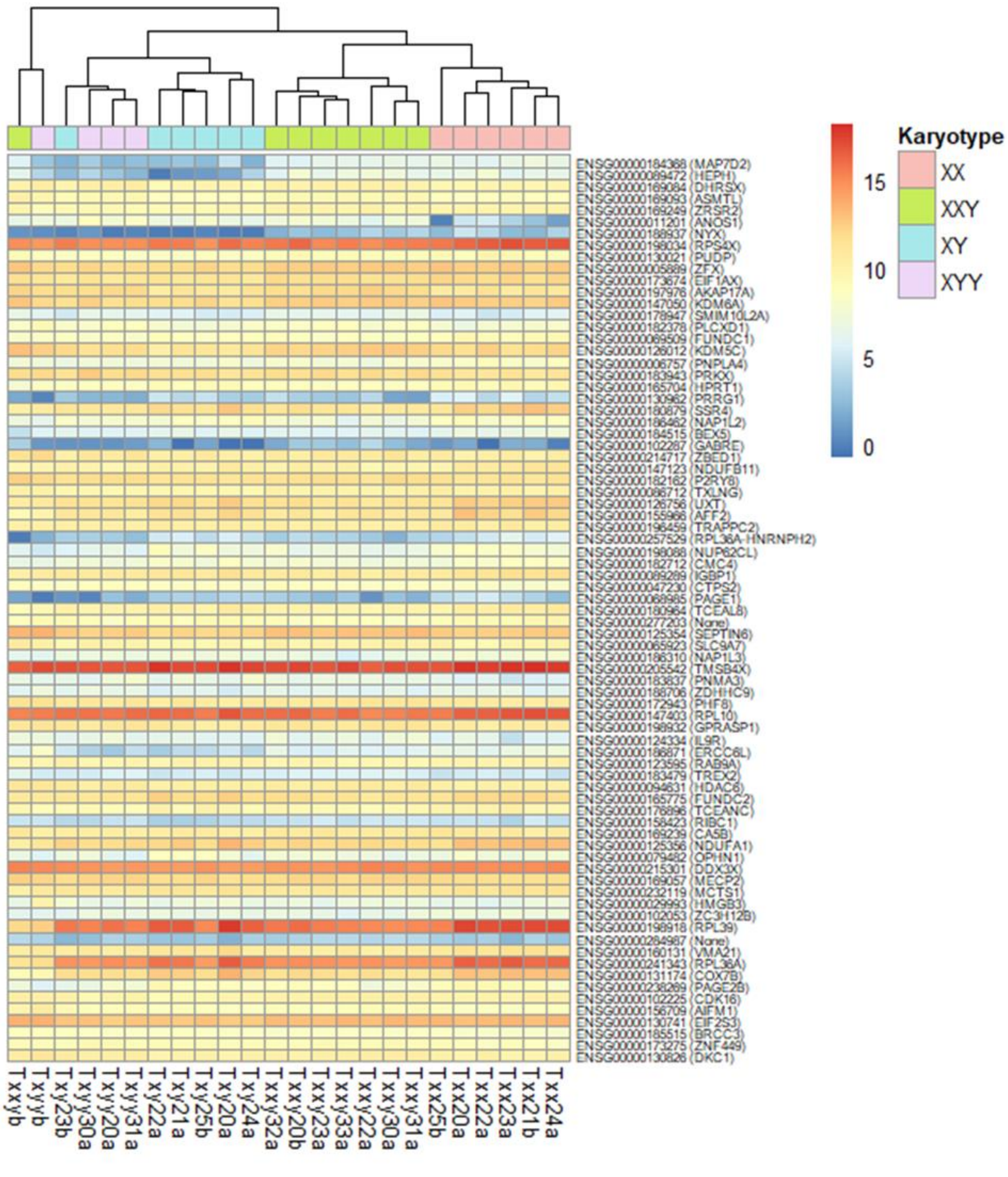
ENSG00000184368 (MAP7D2)
ENSG00000089472 (HEPH)
ENSG00000169084 (DHRSX)
ENSG00000169093 (ASMTL)
ENSG00000169249 (ZRSR2)
ENSG00000011201 (ANOS1)
ENSG00000188937 (NYX)
ENSG00000198034 (RPS4X)
ENSG00000130021 (PUDP)
ENSG00000005889 (ZFX)
ENSG00000173674 (EIF1AX)
ENSG00000197976 (AKAP17A)
ENSG00000147050 (KDM6A)
ENSG00000178947 (SMIM10L2A)
ENSG00000182378 (PLCXD1)
ENSG00000069509 (FUNDC1)
ENSG00000126012 (KDM5C)
ENSG00000006757 (PNPLA4)
ENSG00000183943 (PRKX)
ENSG00000165704 (HPRT1)
ENSG00000130962 (PRRG1)
ENSG00000180879 (SSR4)
ENSG00000186462 (NAP1L2)
ENSG00000184515 (BEX5)
ENSG00000102287 (GABRE)
ENSG00000214717 (ZBED1)
ENSG00000147123 (NDUFB11)
ENSG00000182162 (P2RY8)
ENSG00000086712 (TXLNG)
ENSG00000126756 (UXT)
ENSG00000155966 (AFF2)
ENSG00000196459 (TRAPPC2)
ENSG00000257529 (RPL36A-HNRNPH2)
ENSG00000198088 (NUP62CL)
ENSG00000182712 (CMC4)
ENSG00000089289 (IGBP1)
ENSG00000047230 (CTPS2)
ENSG00000068985 (PAGE1)
ENSG00000180964 (TCEAL8)
ENSG00000277203 (None)
ENSG00000125354 (SEPTIN6)
ENSG00000065923 (SLC9A7)
ENSG00000186310 (NAP1L3)
ENSG00000205542 (TMSB4X)
ENSG00000183837 (PNMA3)
ENSG00000188706 (ZDHHC9)
ENSG00000172943 (PHF8)
ENSG00000147403 (RPL10)
ENSG00000198932 (GPRASP1)
ENSG00000124334 (IL9R)
ENSG00000186871 (ERCC6L)
ENSG00000123595 (RAB9A)
ENSG00000183479 (TREX2)
ENSG00000094631 (HDAC6)
ENSG00000165775 (FUNDC2)
ENSG00000176896 (TCEANC)
ENSG00000158423 (RIBC1)
ENSG00000169239 (CA5B)
ENSG00000125356 (NDUFA1)
ENSG00000079482 (OPHN1)
ENSG00000215301 (DDX3X)
ENSG00000169057 (MECP2)
ENSG00000232119 (MCTS1)
ENSG00000029993 (HMGB3)
ENSG00000102053 (ZC3H12B)
ENSG00000198918 (RPL39)
ENSG00000284987 (None)
ENSG00000160131 (VMA21)
ENSG00000241343 (RPL36A)
ENSG00000131174 (COX7B)
ENSG00000238269 (PAGE2B)
ENSG00000102225 (CDK16)
ENSG00000156709 (AIFM1)
ENSG00000130741 (EIF2S3)
ENSG00000185515 (BRCC3)
ENSG00000173275 (ZNF449)
ENSG00000130826 (DKC1)
Txxyb
Txyyb
Txy23b
Txyy30a
Txyy20a
Txyy31a
Txy22a
Txy21a
Txy25b
Txy20a
Txy24a
Txxy32a
Txxy20b
Txxy23a
Txxy33a
Txxy22a
Txxy30a
Txxy31a
Txx25b
Txx20a
Txx22a
Txx23a
Txx21b
Txx24a
15
10
5
0
Karyotype
XX
XXY
XY
XYY

77

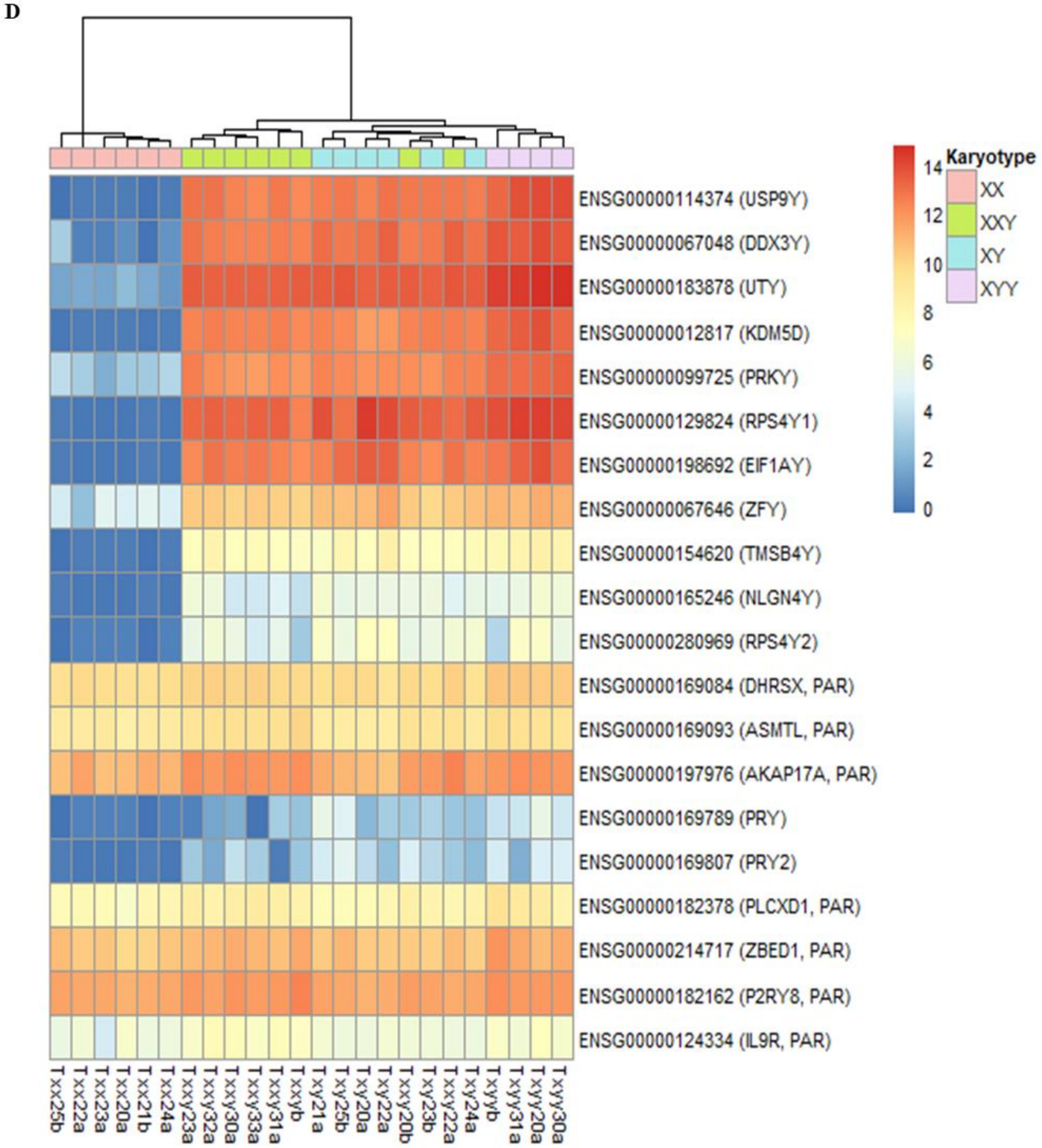


**Fig. S4. Heatmaps of the differentially expressed genes after batch effect was removed in the gonosome aneuploidy dataset, first considering all genes (A) on the X chromosome (number of genes = 103) and (B) on the Y chromosome (number of genes = 45), and then considering only protein-coding genes (C) on the X chromosome (number of genes = 77) and (D) on the Y chromosome (number of genes = 20).**

PAR indicates the genes located on the pseudo-autosomal region of the Y chromosome. Differentially expressed genes were obtained using the Likelihood-ratio test (see **Materials and Methods**) and the list of differentially expressed genes used in each heatmap is available in **Data S1**. Batch effect was removed before graphical display.

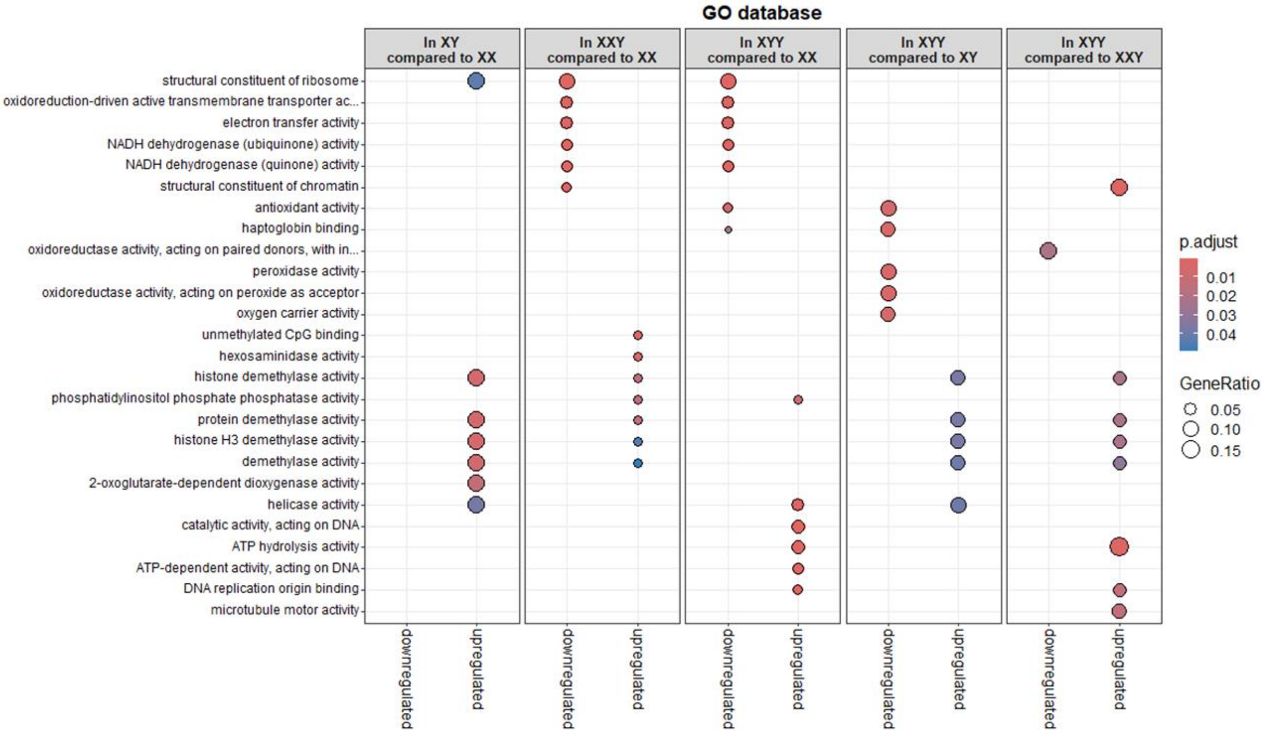


**Fig. S5. Gene ontology analysis performed using the differentially expressed genes (regardless if they are protein-coding or not) and the GO database.**

The plot shows the 5 most significant GO terms and shared GO terms considering downregulated genes and upregulated genes separately obtained from each pairwise karyotype comparison (when applicable).

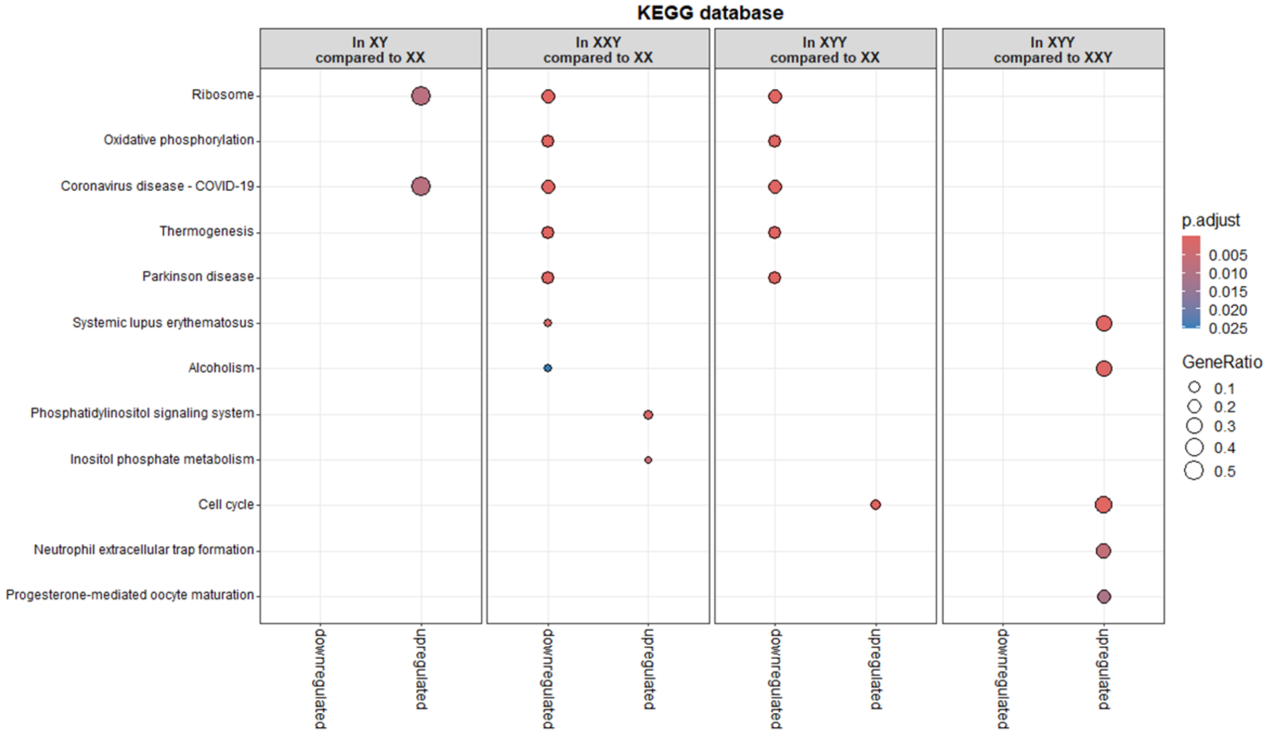


**Fig. S6. Pathway enrichment analysis performed using the differentially expressed genes (regardless if they are protein-coding or not) and the KEGG database.**

The plot shows the 5 most significant pathways and shared pathways considering downregulated genes and upregulated genes separately obtained from each pairwise karyotype comparison (when applicable).

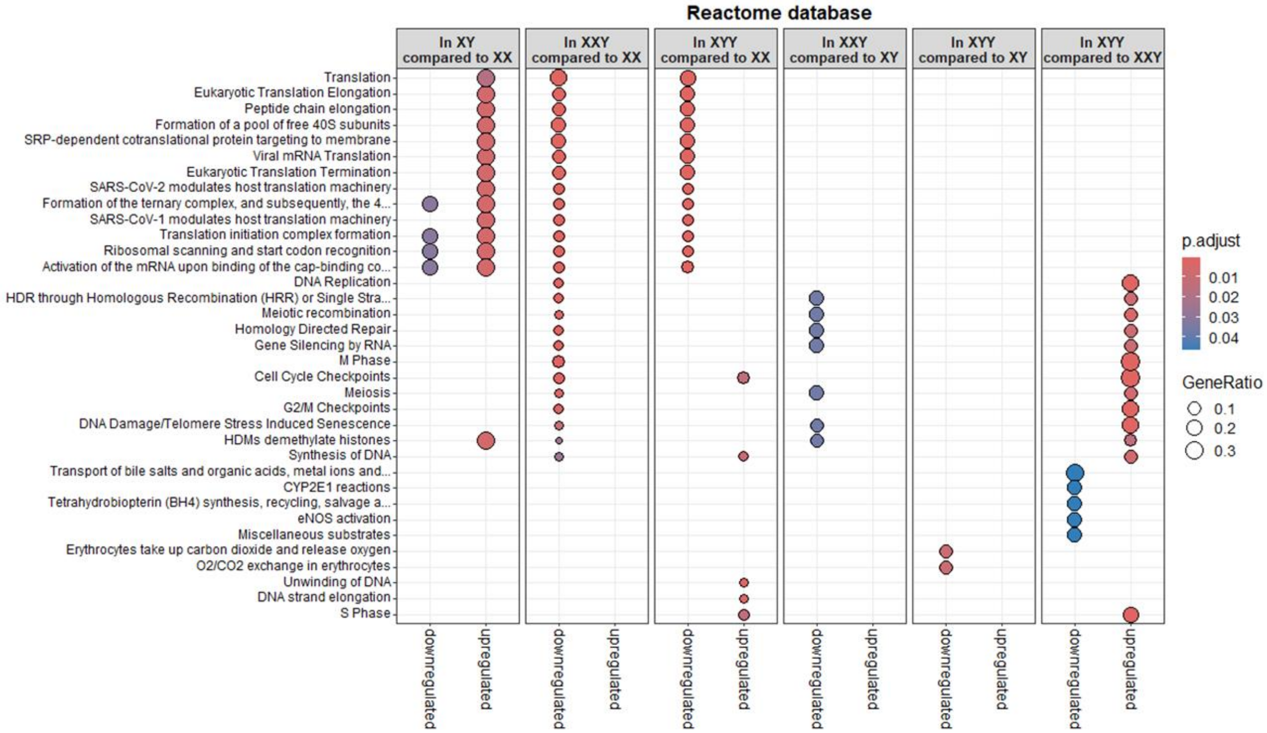


**Fig. S7. Pathway enrichment analysis performed using the differentially expressed genes (regardless if they are protein-coding or not) and the Reactome database.**

The plot shows the 5 most significant pathways and shared pathways considering downregulated genes and upregulated genes separately obtained from each pairwise karyotype comparison (when applicable).

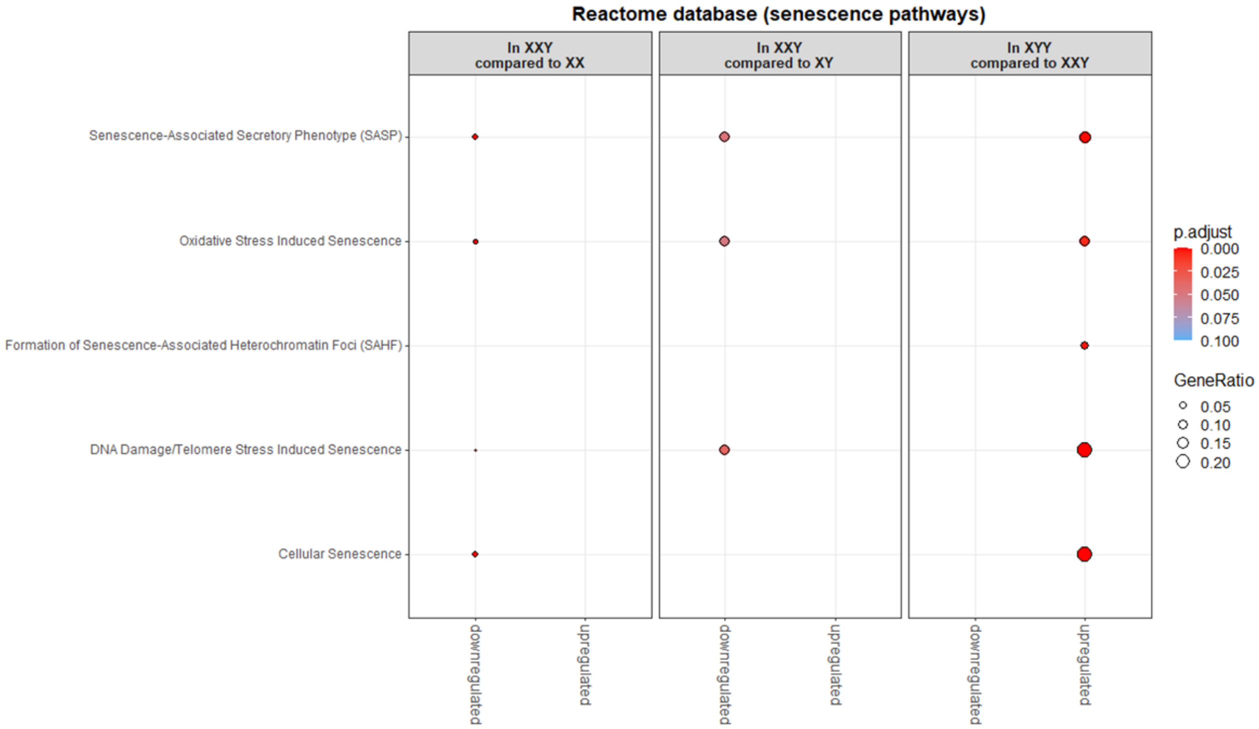


**Fig. S8. Pathway enrichment analysis performed using the differentially expressed genes (regardless if they are protein-coding or not) and focused on the significant terms from the Reactome database including the word "senescence" in their description.**

The plot shows the 5 pathways including the word "senescence" in their description if they were significant considering downregulated genes and upregulated genes separately obtained from each pairwise karyotype comparison (when applicable).

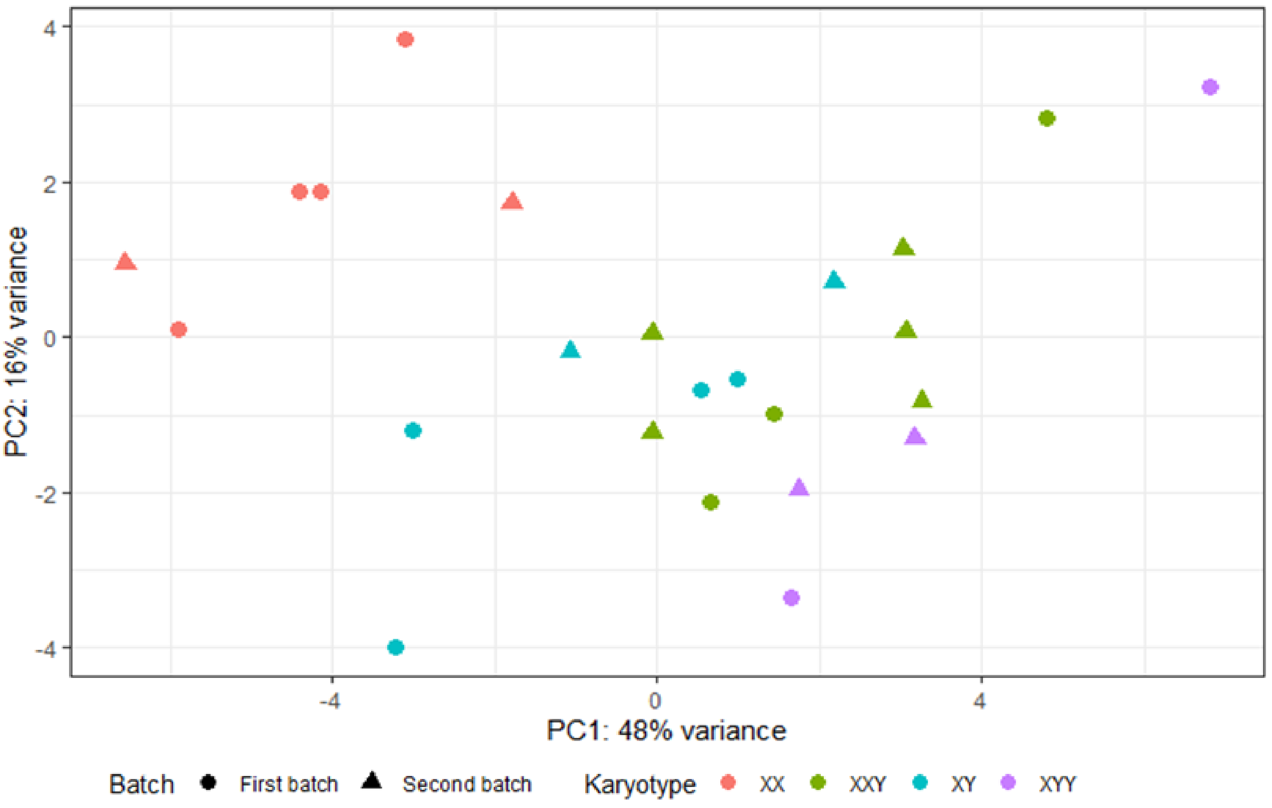


**Fig. S9. Principal component analysis on the 50 TEs explaining the most variance in the gonosome aneuploidy dataset after batch effect was removed.**

Each dot represents one individual. Dots are colored according to karyotype (four levels: XX, XY, XXY, XYY) and shaped according to batch (two levels: first batch, second batch). Batch effect was removed before graphical display. PC: Principal Component.

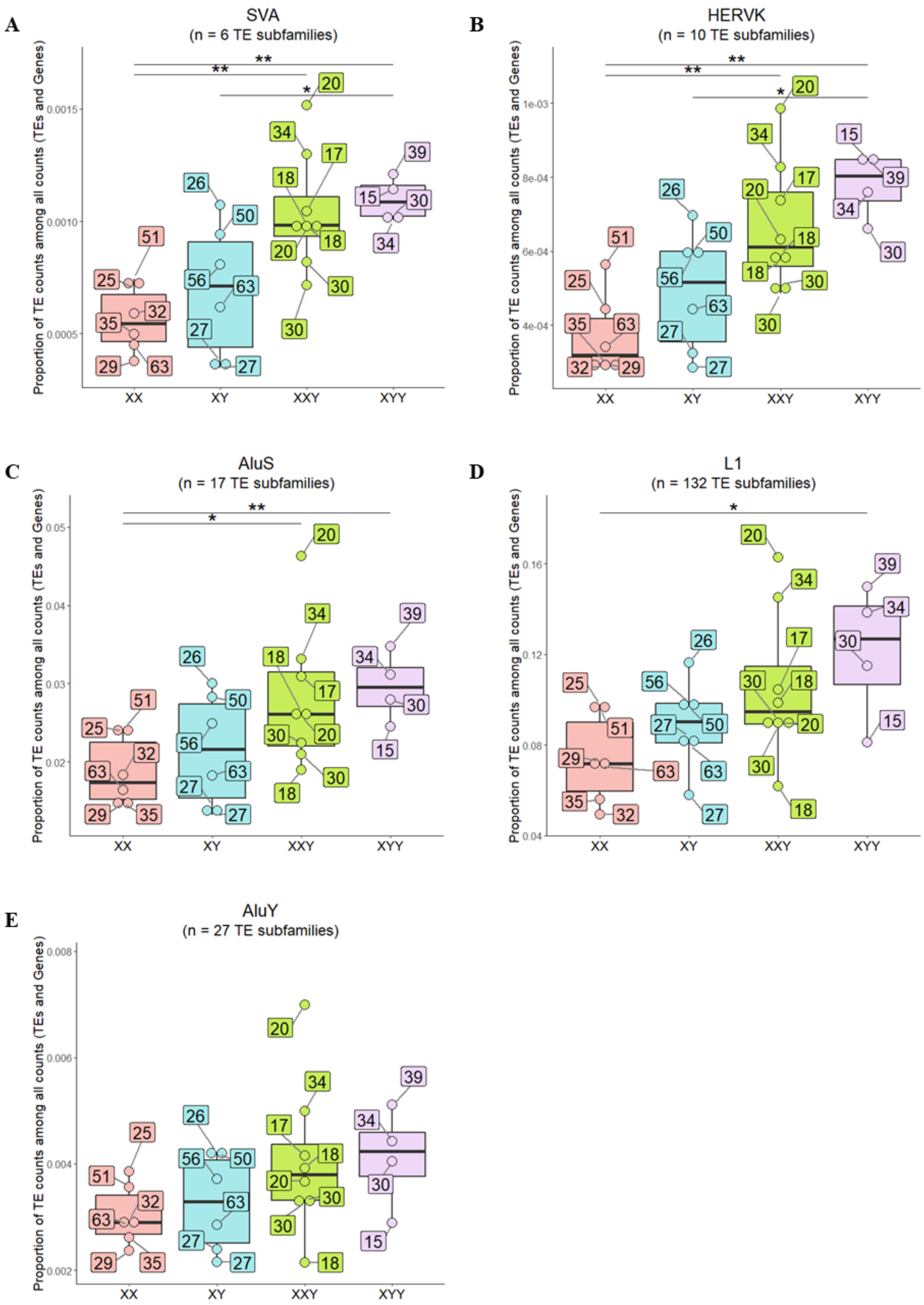
A
SVA
(n = 6 TE subfamilies)
B
HERVK
(n = 10 TE subfamilies)
C
AluS
(n = 17 TE subfamilies)
D
L1
(n = 132 TE subfamilies)
E
AluY
(n = 27 TE subfamilies)
Proportion of TE counts among all counts (TEs and Genes)
XX
XY
XXY
XYY
0.0005
0.0010
0.0015
4e-04
6e-04
8e-04
1e-03
0.02
0.03
0.04
0.05
0.04
0.08
0.12
0.16
0.002
0.004
0.006
0.008

**Fig. S10. TE expression in the different karyotypes after removing batch effect and considering TE subfamilies belonging to (A) SVA, (B) HERVK, (C) AluS, (D) L1, or (E) AluY elements.**

Global TE expression was measured in each karyotype (x-axis) as the proportion of TE read counts among all read counts (TEs and genes) (y-axis). In this calculation, read counts were used after DESeq2 normalization ("normalized counts") to remove any depth sequencing bias. Each dot represents one individual. The age of each individual is in a label box linked to each dot. Batch effect was removed before graphical display. P-values from Wilcoxon test comparing pairwise karyotypes: (*) < 0.05 and (**) < 0.01.

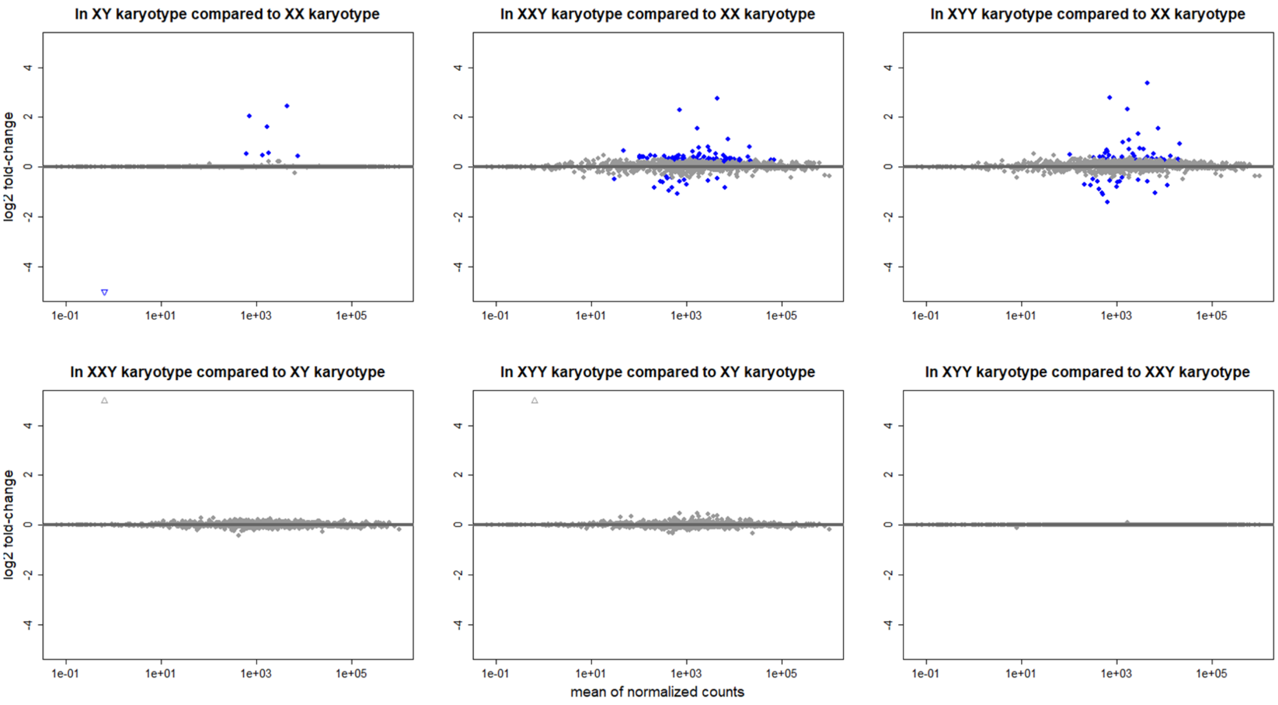


**Fig. S11. MA plots on the 1246 TE subfamilies in each pairwise karyotype comparison after batch adjustment in the gonosome aneuploidy dataset.**

Minus-Average plots (MA plots) represent, for each protein-coding gene, the $\log_2$(normalized counts in group 1) – $\log_2$(normalized counts in group 2) on the y-axis and the mean expression (i.e. mean of normalized counts) across all the samples on the x-axis. Each dot or triangle represents one TE subfamily. Blue dots show TE subfamily with an adjusted p-value < 0.05. TE subfamily for which log2 fold-change is greater than 4 or less than -4 are represented by an empty triangle. Positive log2 fold-change means that the TE subfamily is upregulated in the first karyotype (e.g. XYY in the “XYY karyotype vs XX karyotype” comparison) compared to the second (e.g. XX in the “XYY karyotype vs XX karyotype” comparison). Negative log2 fold-change means that the TE subfamily is downregulated in the first karyotype (e.g. XYY in the “XYY karyotype vs XX karyotype” comparison) compared to the second (e.g. XX in the “XYY karyotype vs XX karyotype” comparison).

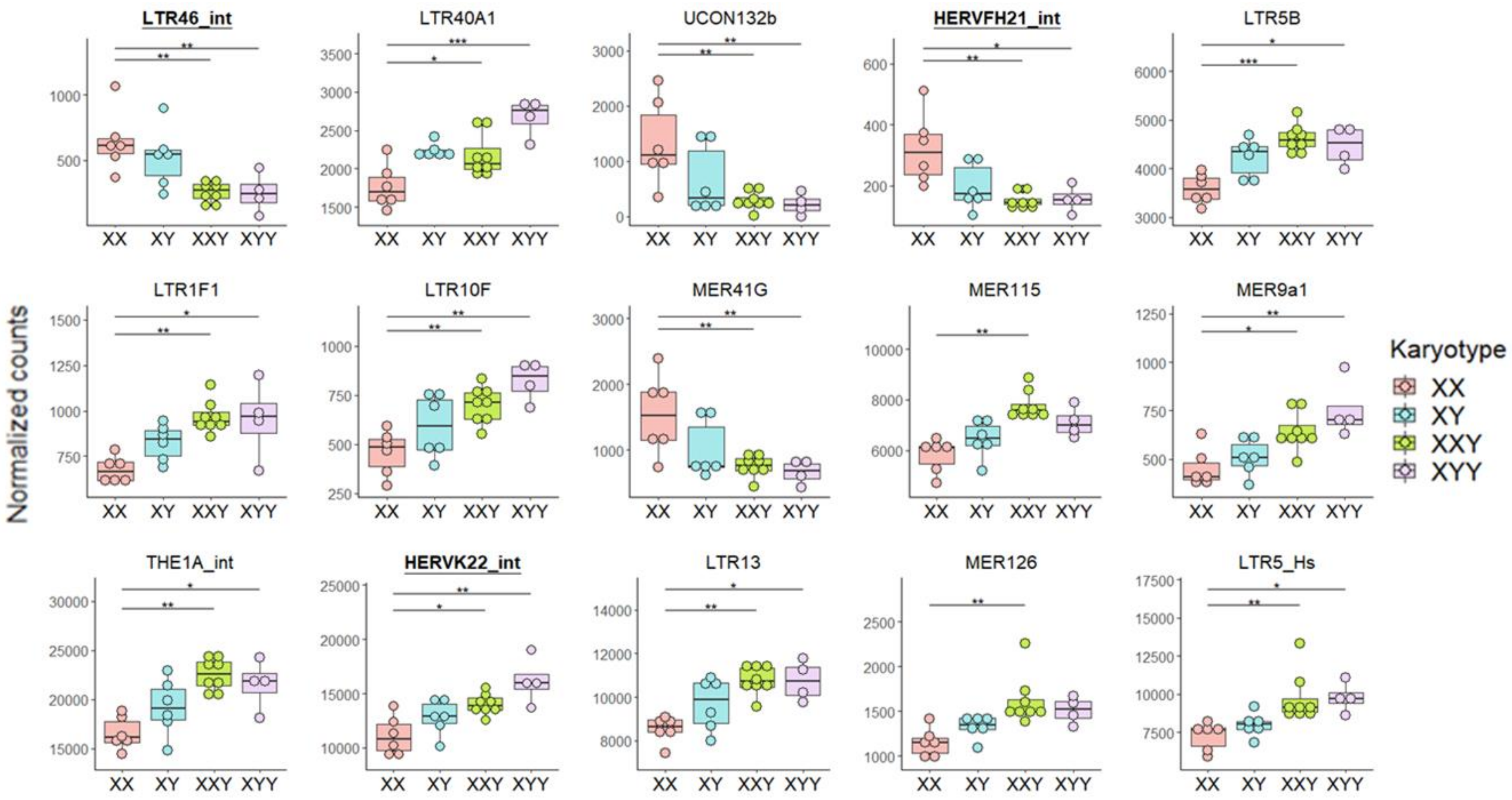

LTR46_int
LTR40A1
UCON132b
HERVFH21_int
LTR5B
LTR1F1
LTR10F
MER41G
MER115
MER9a1
THE1A_int
HERVK22_int
LTR13
MER126
LTR5_Hs
Normalized counts
XX
XY
XXY
XYY
Karyotype
XX
XY
XXY
XYY


135

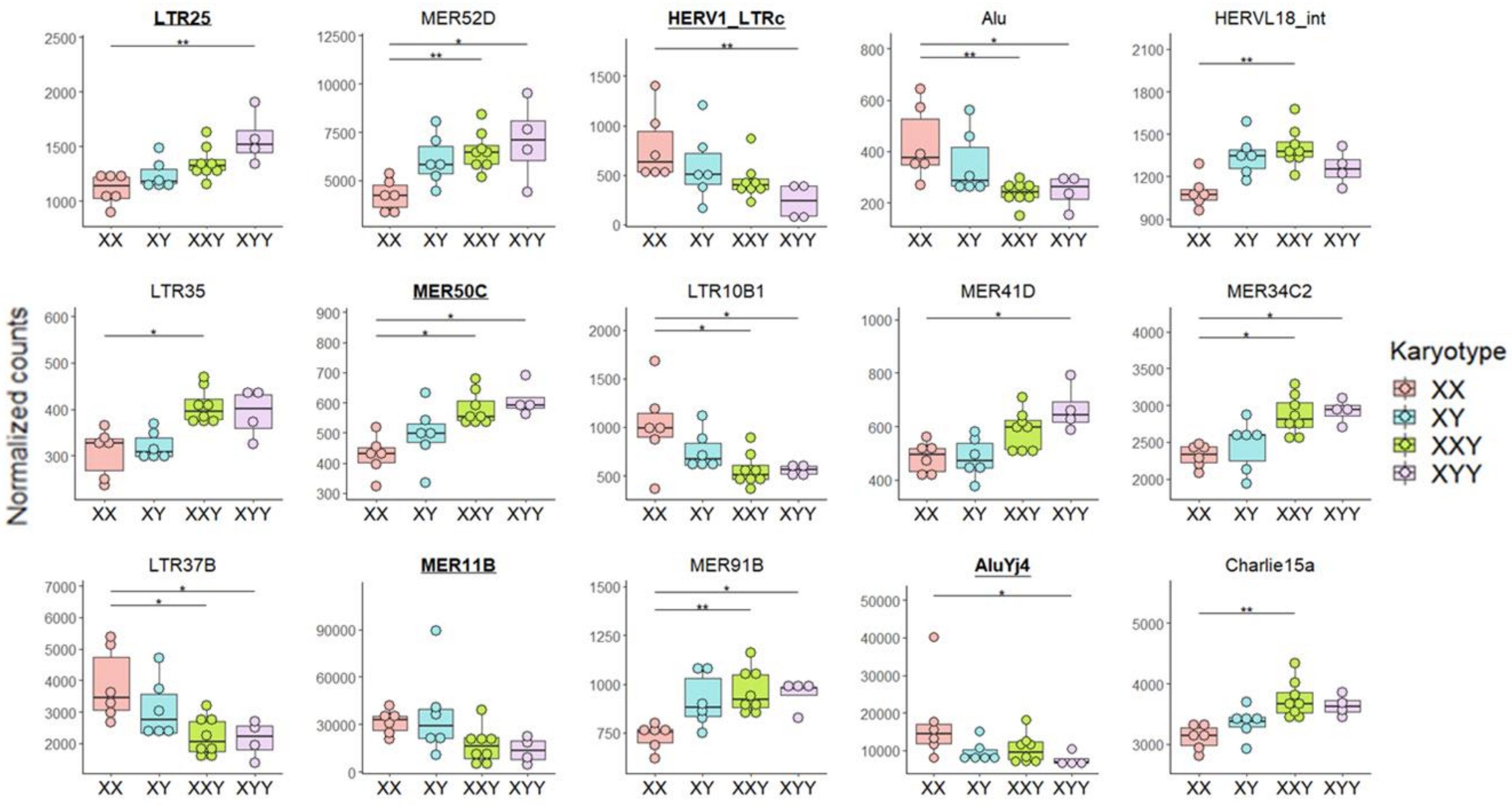
LTR25
MER52D
HERV1_LTRc
Alu
HERVL18_int
LTR35
MER50C
LTR10B1
MER41D
MER34C2
LTR37B
MER11B
MER91B
AluYj4
Charlie15a
Normalized counts
XX XY XXY XYY
Karyotype
XX
XY
XXY
XYY

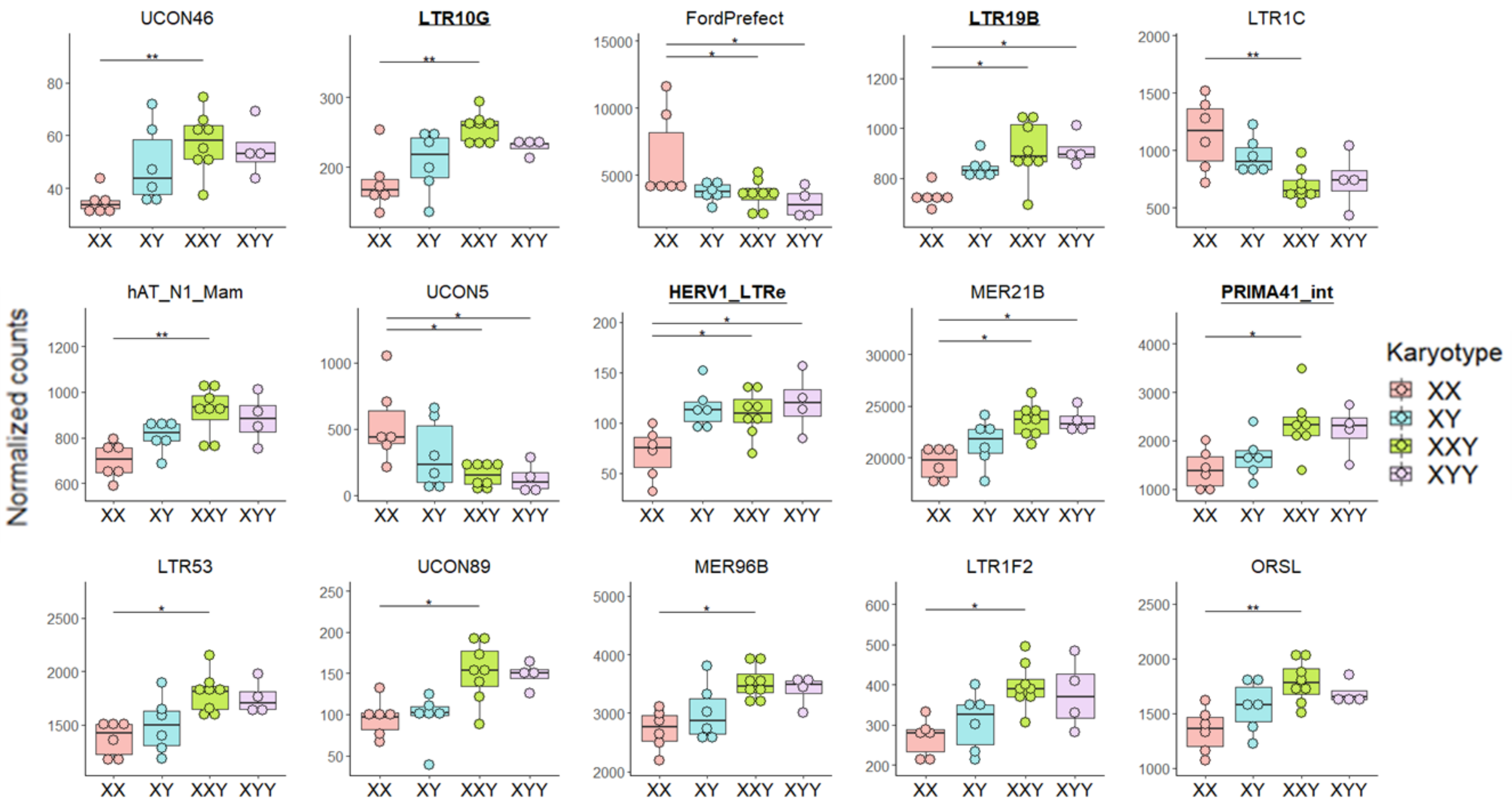


**Fig. S12. Boxplots for the 16th to 60th most significantly differentially expressed TE subfamilies according to karyotype using the likelihood-ratio test, after removing batch effect.**

Specific TE subfamily expression was estimated using the number of read counts after DESeq2 normalization ("normalized counts") to remove any depth sequencing bias. Each dot represents one individual. Dots are colored according to karyotype (four levels: XX, XY, XXY, XYY). Batch effect was removed before graphical display. Adjusted p-values from DESeq2: (*) < 0.05, (**) < 0.01, (***) < 0.001. Underlined bold TE subfamilies are enriched in the Y chromosome (binomial test adjusted p-value $< 0.05$ and number of observed copies located on Y chromosome > expected). See **Figure 2** for the 15 most significantly differentially expressed TE subfamilies according to karyotype.

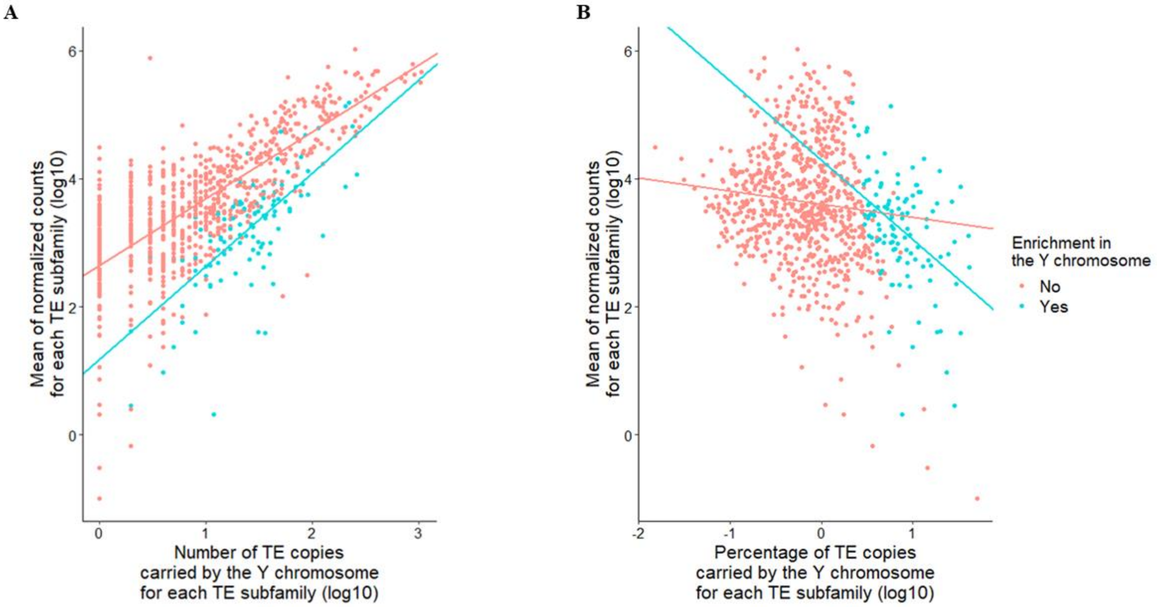


**Fig S13. TE subfamily expression according to (A) the number of their TE copies carried by the Y chromosome and (B) the proportion of their TE copies carried by the Y chromosome.**

(**A**) Regression lines were generated using a linear model using generalized least square estimation with an exponential variance function structure and that regressed the log10 mean of normalized counts across all individuals on two independent variables: the log10 number of TE copies carried by the Y chromosome (estimation = 1.046, p-value <0.001), the enrichment in the Y chromosome (factor variable : yes compared to no, estimation = -1.462, p-value < 0.001) and their interaction (estimation = 0.4098, p-value = 0.003). Note that TE subfamilies with number of TE copies carried by the Y chromosome equal to zero were removed before statistical analysis.

(**B**) Regression lines were generated using a linear model using generalized least square estimation with an exponential variance function structure and that regressed the log10 mean of normalized counts across all individuals on two independent variables : the log10 percentage of TE copies carried by the Y chromosome (estimation = -0.205, p-value = 0.001), the enrichment in the Y chromosome (factor variable : yes compared to no, estimation = 0.689, p-value = 0.005) and their interaction (estimation =-1.035, p-value < 0.001). Note that TE subfamilies with percentage of TE copies carried by the Y chromosome equal to zero were removed before statistical analysis.

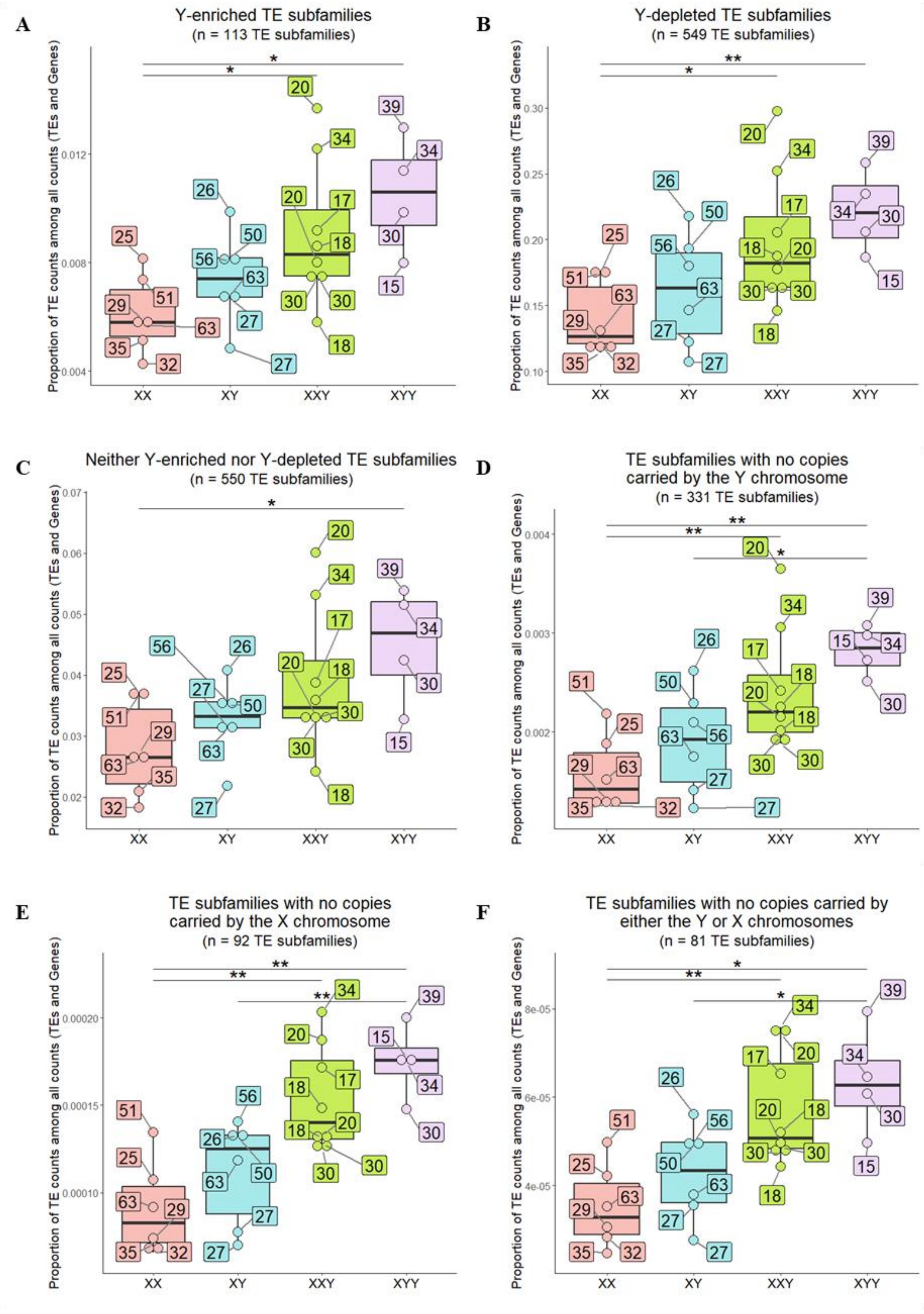
A
Y-enriched TE subfamilies
(n = 113 TE subfamilies)
Proportion of TE counts among all counts (TEs and Genes)
0.004
0.008
0.012
XX
XY
XXY
XYY
B
Y-depleted TE subfamilies
(n = 549 TE subfamilies)
Proportion of TE counts among all counts (TEs and Genes)
0.10
0.15
0.20
0.25
0.30
XX
XY
XXY
XYY
C
Neither Y-enriched nor Y-depleted TE subfamilies
(n = 550 TE subfamilies)
Proportion of TE counts among all counts (TEs and Genes)
0.02
0.03
0.04
0.05
0.06
0.07
XX
XY
XXY
XYY
D
TE subfamilies with no copies
carried by the Y chromosome
(n = 331 TE subfamilies)
Proportion of TE counts among all counts (TEs and Genes)
0.002
0.003
0.004
XX
XY
XXY
XYY
E
TE subfamilies with no copies
carried by the X chromosome
(n = 92 TE subfamilies)
Proportion of TE counts among all counts (TEs and Genes)
0.00010
0.00015
0.00020
XX
XY
XXY
XYY
F
TE subfamilies with no copies carried by
either the Y or X chromosomes
(n = 81 TE subfamilies)
Proportion of TE counts among all counts (TEs and Genes)
4e-05
6e-05
8e-05
XX
XY
XXY
XYY

**Fig. S14. TE expression in the different karyotypes after removing batch effect and considering all TE subfamilies which were (A) Y-enriched TE subfamilies, (B) Y-enriched TE subfamilies, (C) neither Y-enriched nor Y-depleted TE subfamilies, (D) TE subfamilies with no copies carried by the Y chromosome, (E) TE subfamilies with no copies carried by the X chromosome, or (F) TE subfamilies with no copies carried by either the Y or X chromosomes.**

Global TE expression was measured in each karyotype (x-axis) as the proportion of TE read counts among all read counts (TEs and genes) (y-axis). In this calculation, read counts were used after DESeq2 normalization ("normalized counts") to remove any depth sequencing bias. Each dot represents one individual. The age of each individual is in a label box linked to each dot. Batch effect was removed before graphical display. P-values from Wilcoxon test comparing pairwise karyotypes: (*) < 0.05 and (**) < 0.01.

See **Data S5** to know which TE subfamilies are Y-enriched (binomial test adjusted p-value < 0.05 and number of observed copies located on Y chromosome > expected)), Y-depleted (binomial test adjusted p-value < 0.05 and number of observed copies located on Y chromosome < expected) and to know the number of TE copy carried by the X and Y chromosomes according to the human reference sequence file of TE insertions.

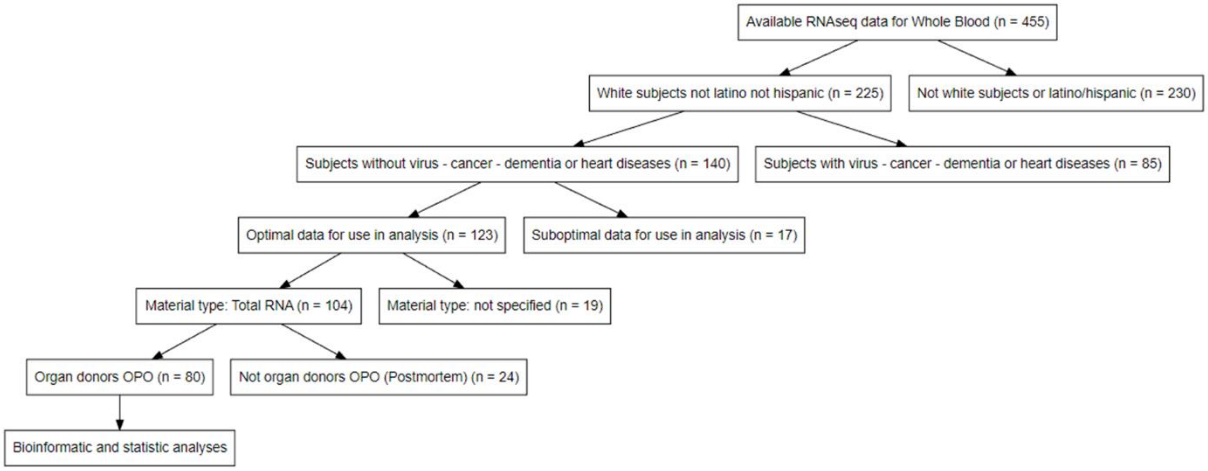


**Fig. S15. Flowchart showing the selection criteria used to constitute the filtered GTEx dataset (no disease group).**

OPO: Organ Procurement Organization.

**A** **In females**

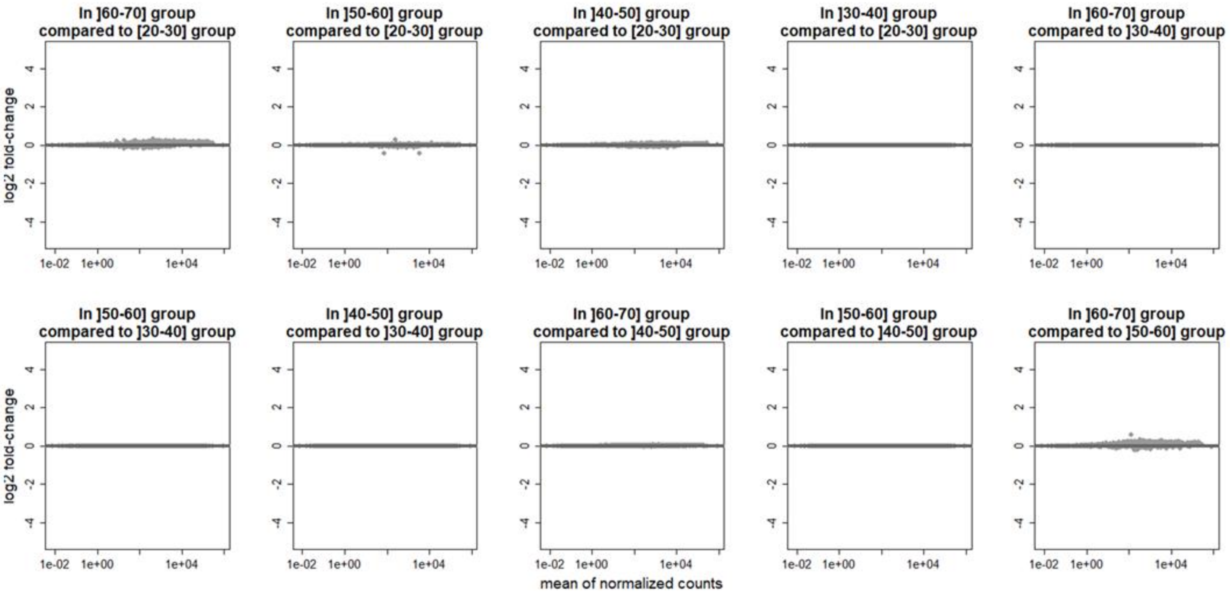


**B** **In males**

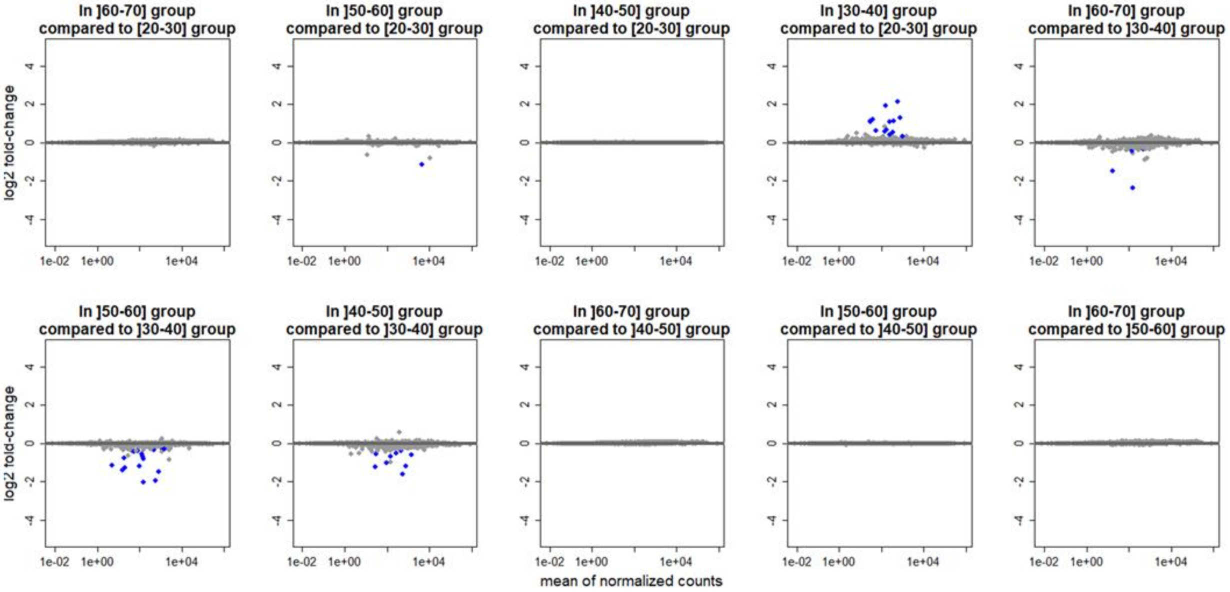

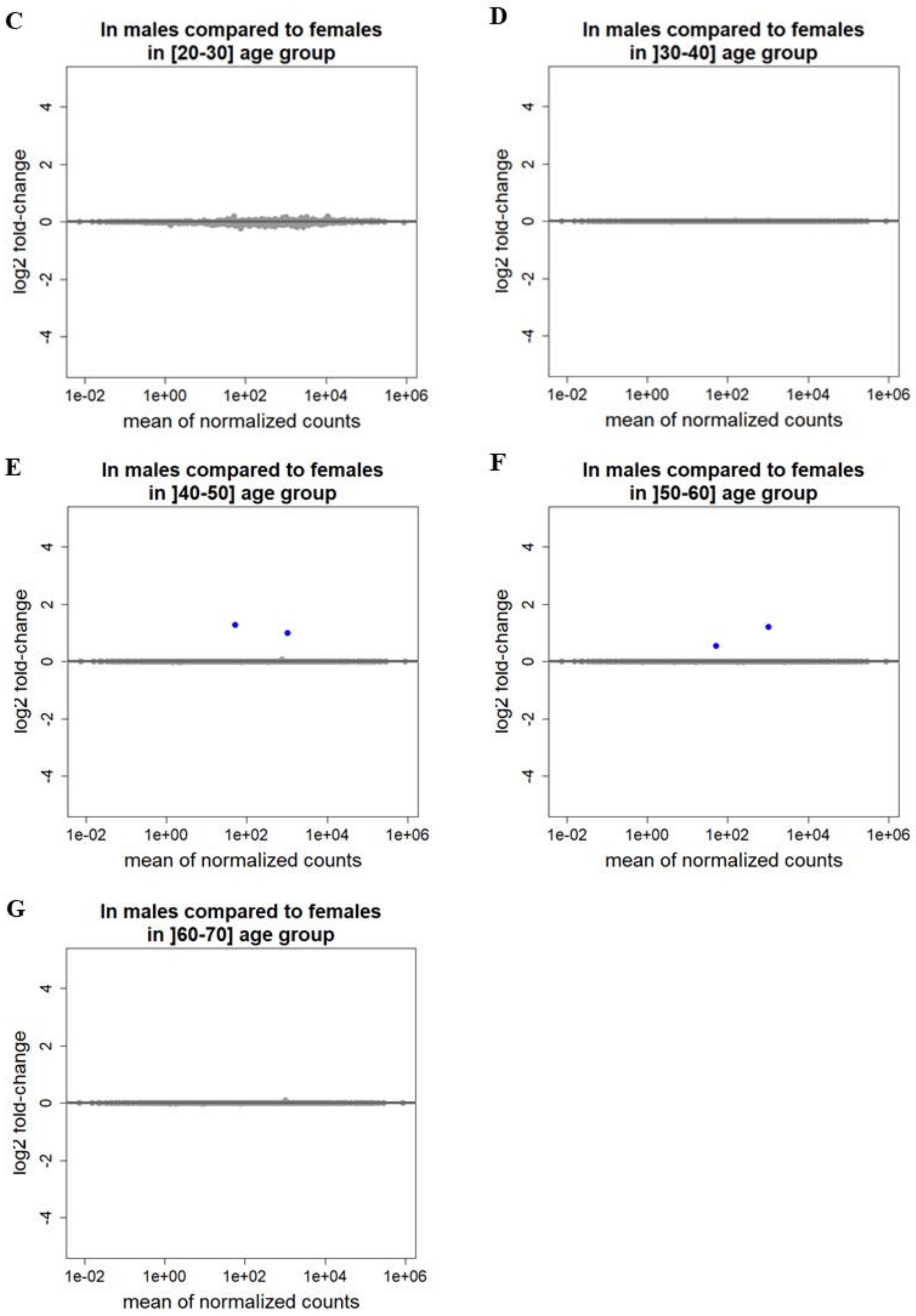
C
In males compared to females
in [20-30] age group
D
In males compared to females
in ]30-40] age group
E
In males compared to females
in ]40-50] age group
F
In males compared to females
in ]50-60] age group
G
In males compared to females
in ]60-70] age group
log2 fold-change
mean of normalized counts
-4
-2
0
2
4
1e-02
1e+00
1e+02
1e+04
1e+06

**Fig. S16. MA plots on the 1246 TE subfamilies using the filtered GTEx dataset (no disease group): (A) pairwise comparisons of age group in females, (B) pairwise comparisons of age group in males, (C) comparison between females and males in age group 20-30 years old, (D) comparison between females and males in age group 30-40 years old, (E) comparison between females and males in age group 40-50 years old, (F) comparison between females and males in age group 50-60 years old, and (G) comparison between females and males in age group 60-70 years old.**

Minus-Average plots (MA plots) represent, for each protein-coding gene, the $log_2$(normalized counts in group 1) – $log_2$(normalized counts in group 2) on the y-axis and the mean expression (i.e. mean of normalized counts) across all the samples on the x-axis. Each dot represents one TE subfamily. Blue dots show TE subfamilies with an adjusted p-value < 0.05. Positive log2 fold-change means that the TE subfamily is upregulated in the first group (e.g. Males in the “Males vs Females” comparison) compared to the second (e.g. Females in the “Males vs Females” comparison). Negative log2 fold-change means that the TE subfamily is downregulated in the first group (e.g. Males in the “Males vs Females” comparison) compared to the second (e.g. Females in the “Males vs Females” comparison).

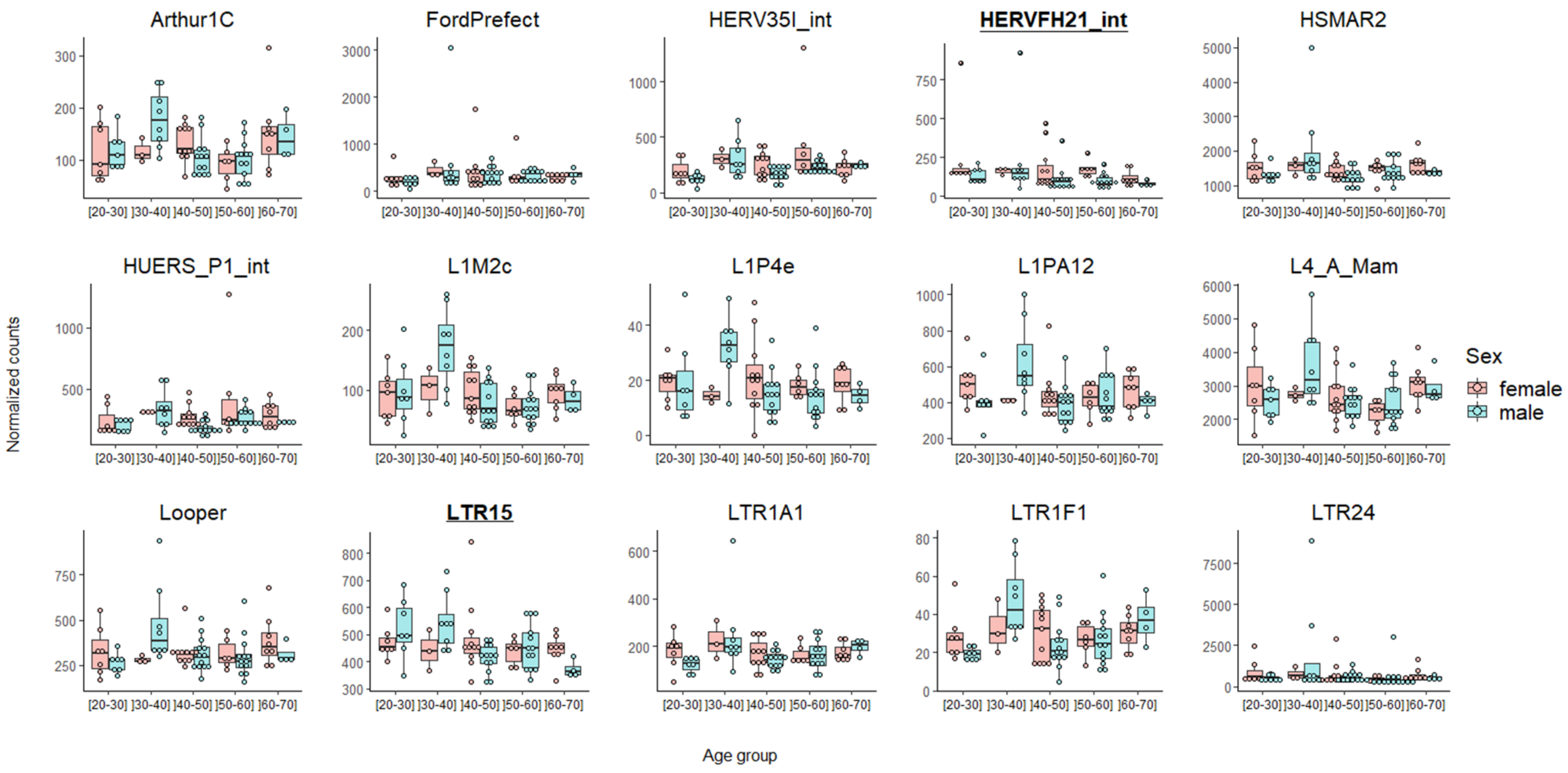
Arthur1C
FordPrefect
HERV35I_int
HERVFH21_int
HSMAR2
HUERS_P1_int
L1M2c
L1P4e
L1PA12
L4_A_Mam
Looper
LTR15
LTR1A1
LTR1F1
LTR24
[20-30] ]30-40] ]40-50] ]50-60] ]60-70]
Normalized counts
Age group
Sex
female
male

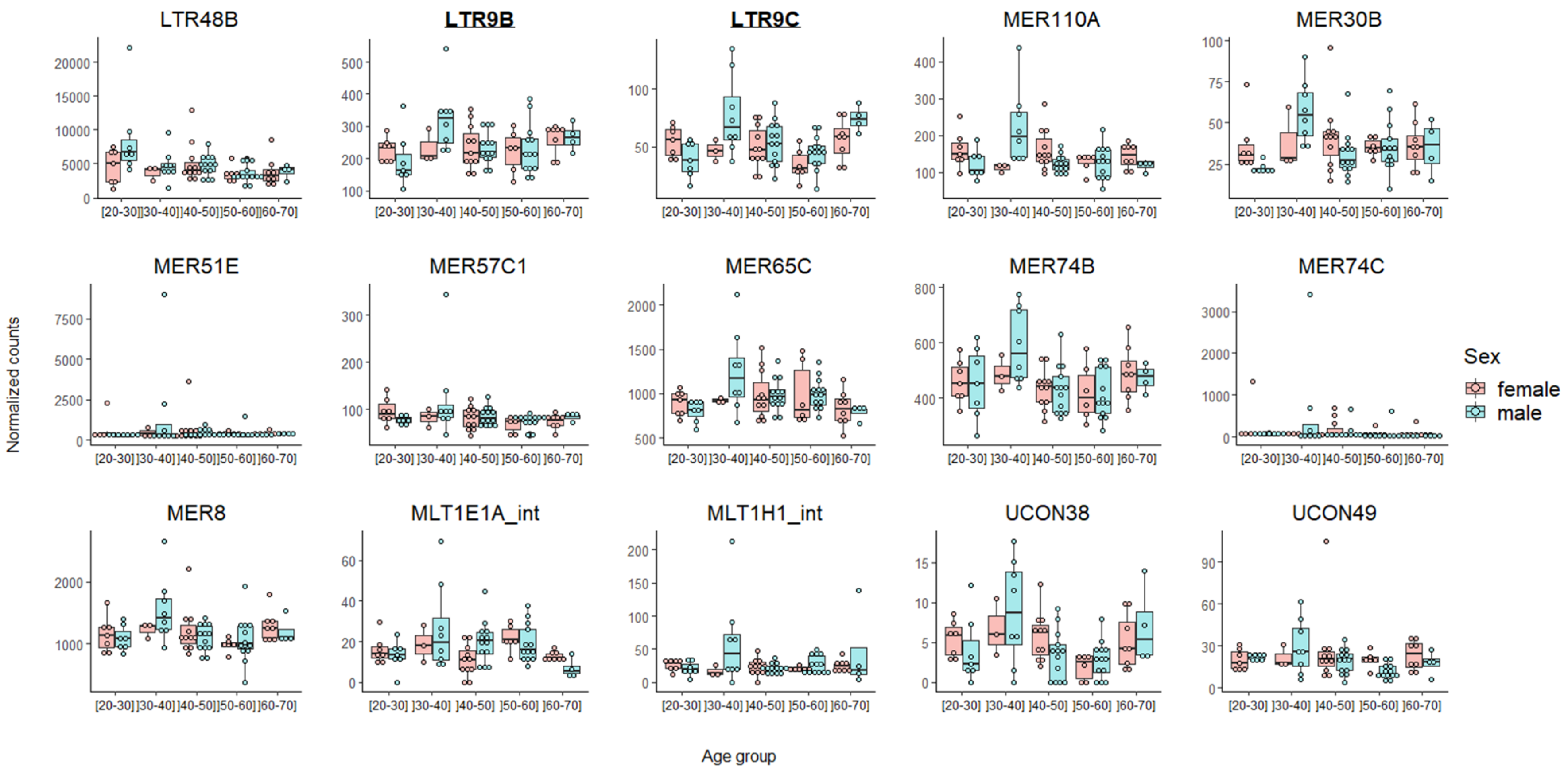
LTR48B
LTR9B
LTR9C
MER110A
MER30B
MER51E
MER57C1
MER65C
MER74B
MER74C
MER8
MLT1E1A_int
MLT1H1_int
UCON38
UCON49
Normalized counts
Age group
[20-30] ]30-40] ]40-50] ]50-60] ]60-70]
Sex
female
male

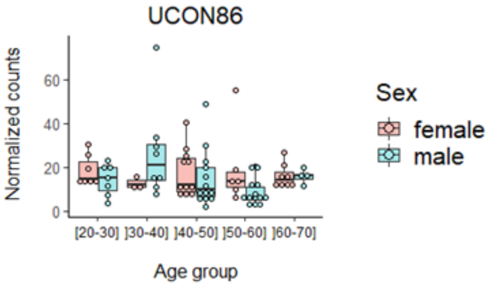


**Fig. S17. Boxplots of normalized counts according to sex and age group, for TE subfamilies that are significantly differentially expressed between two age groups in males or females in the filtered GTEx dataset (no disease group).**

Specific TE subfamily expression was estimated using the number of read counts after DESeq2 normalization ("normalized counts") to remove any depth sequencing bias. Each dot represents one individual. Dots are colored according to sex. Sex variable has two levels: female, male. Age group has five levels: [20-30], ]30-40], ]40-50], ]50-60], and ]60-70]. Underlined bold TE subfamilies are enriched in the Y chromosome (binomial test adjusted p-value < 0.05 and number of observed copies located on Y chromosome > expected).

For more details on the group comparisons in which TE subfamilies were significantly differentially expressed, see **Table S9**.

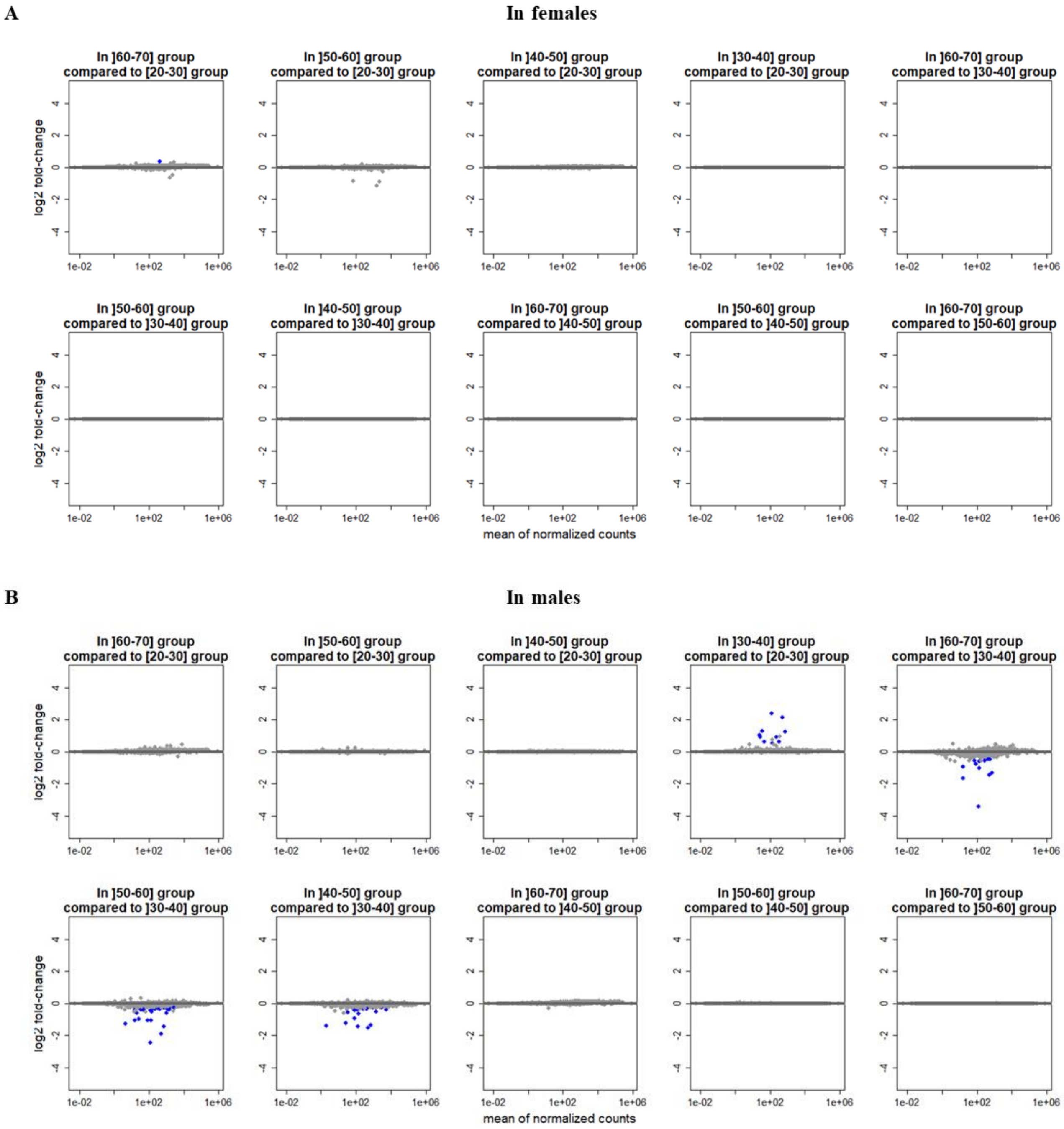
A
In females
In ]60-70] group compared to [20-30] group
In ]50-60] group compared to [20-30] group
In ]40-50] group compared to [20-30] group
In ]30-40] group compared to [20-30] group
In ]60-70] group compared to ]30-40] group
In ]50-60] group compared to ]30-40] group
In ]40-50] group compared to ]30-40] group
In ]60-70] group compared to ]40-50] group
In ]50-60] group compared to ]40-50] group
In ]60-70] group compared to ]50-60] group
log2 fold-change
mean of normalized counts
B
In males
In ]60-70] group compared to [20-30] group
In ]50-60] group compared to [20-30] group
In ]40-50] group compared to [20-30] group
In ]30-40] group compared to [20-30] group
In ]60-70] group compared to ]30-40] group
In ]50-60] group compared to ]30-40] group
In ]40-50] group compared to ]30-40] group
In ]60-70] group compared to ]40-50] group
In ]50-60] group compared to ]40-50] group
In ]60-70] group compared to ]50-60] group
log2 fold-change
mean of normalized counts

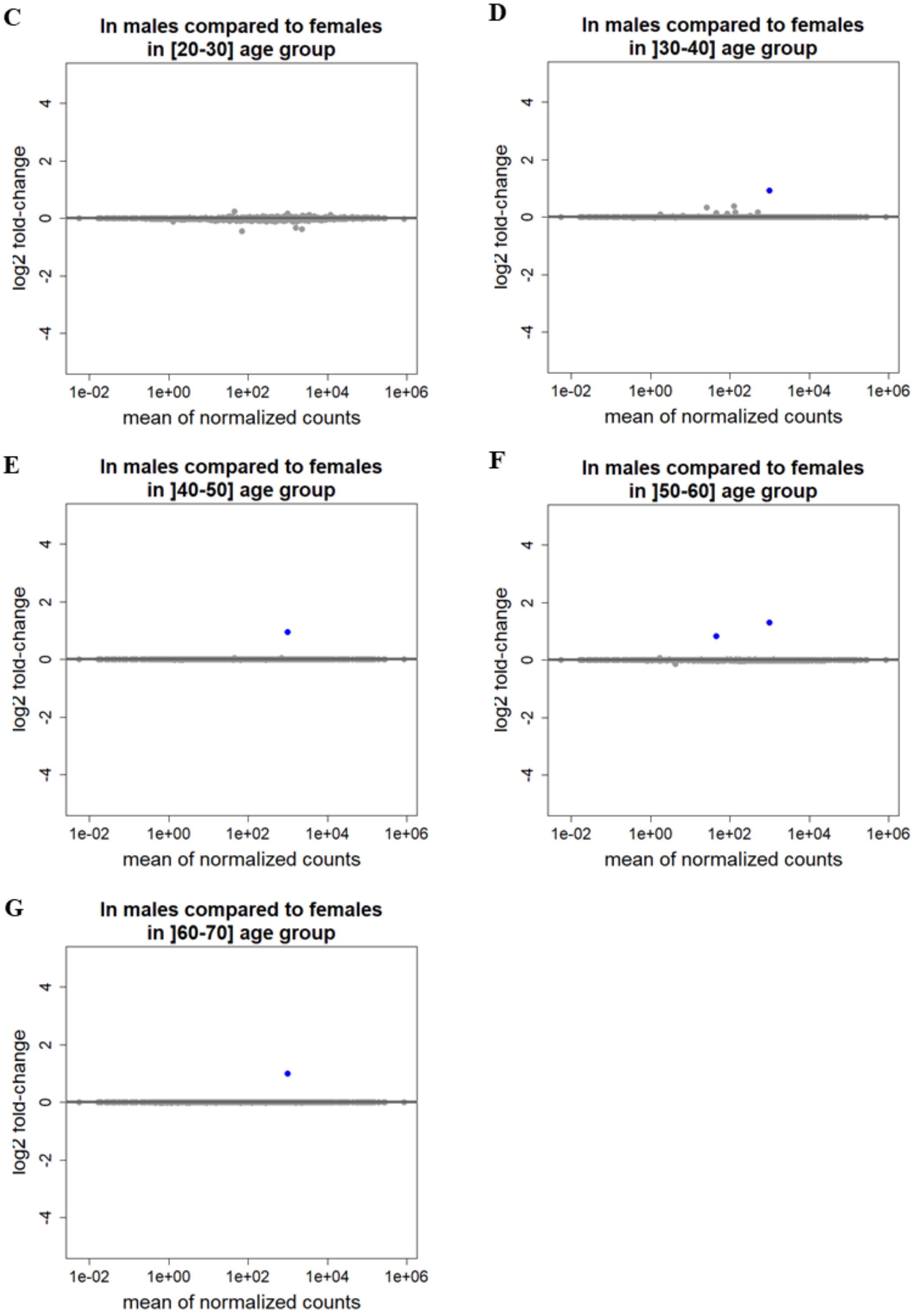


**Fig. S18. MA plots on the 1246 TE subfamilies using the filtered GTEx dataset (group including subjects with current and antecedents of cancer and cardiovascular diseases): (A) pairwise comparisons of age group in**

**females, (B) pairwise comparisons of age group in males, (C) comparison between females and males in age group 20-30 years old, (D) comparison between females and males in age group 30-40 years old, (E) comparison between females and males in age group 40-50 years old, (F) comparison between females and males in age group 50-60 years old, and (G) comparison between females and males in age group 60-70 years old.**

Minus-Average plots (MA plots) represent, for each protein-coding gene, the $\log_2$(normalized counts in group 1) – $\log_2$(normalized counts in group 2) on the y-axis and the mean expression (i.e. mean of normalized counts) across all the samples on the x-axis.  Each dot represents one TE subfamily. Blue dots show TE subfamilies with an adjusted p-value < 0.05. Positive log2 fold-change means that the TE subfamily is upregulated in the first group (e.g. Males in the “Males vs Females” comparison) compared to the second (e.g. Females in the “Males vs Females” comparison). Negative log2 fold-change means that the TE subfamily is downregulated in the first group (e.g. Males in the “Males vs Females” comparison) compared to the second (e.g. Females in the “Males vs Females” comparison).

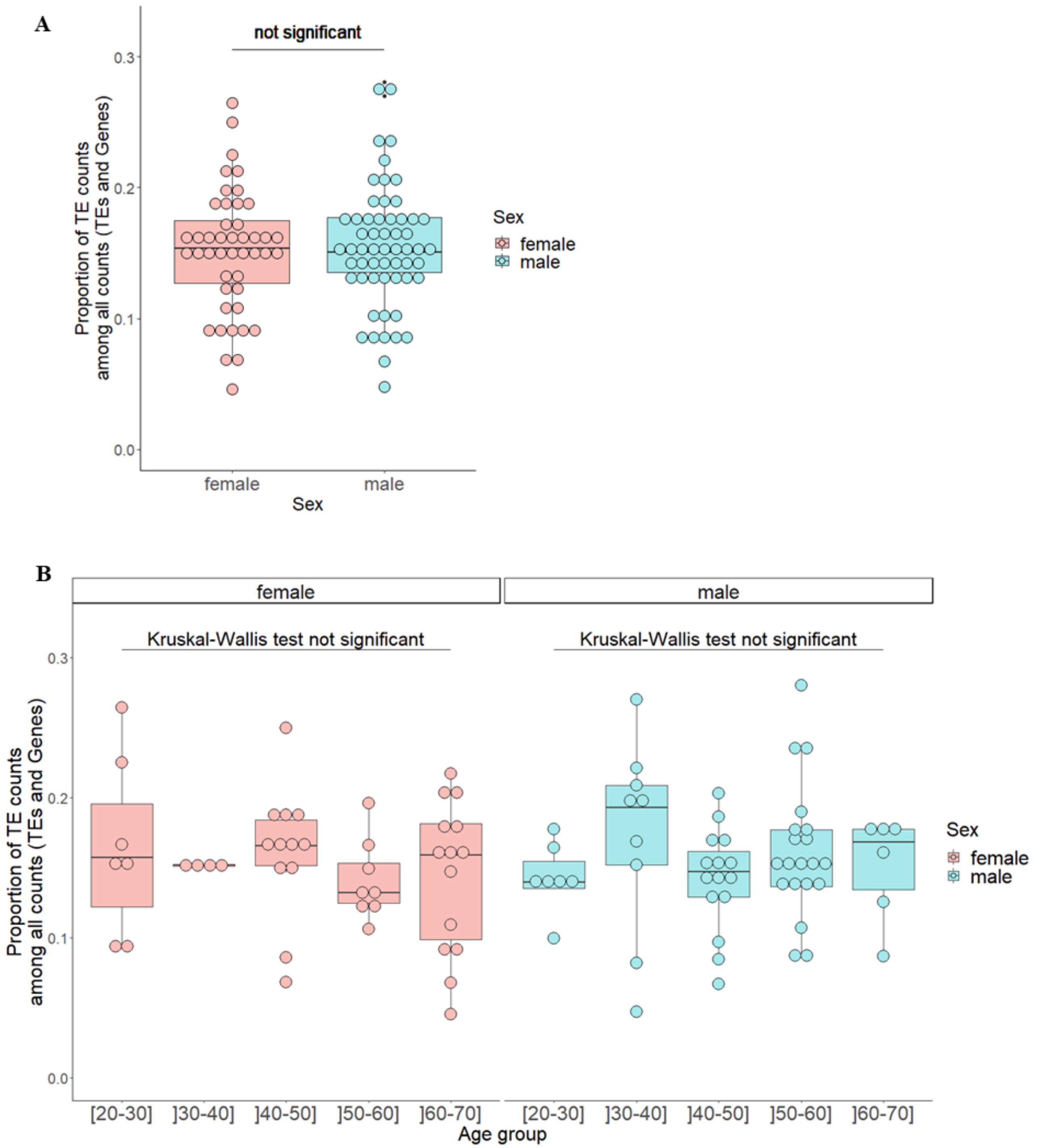


**Fig. S19. TE expression in the filtered GTEx dataset (in the group including subjects with current and antecedents of cancer and cardiovascular diseases).**

Global TE expression was measured in each karyotype (x-axis) as the proportion of TE read counts among all read counts (TEs and genes) (y-axis). In this calculation, read counts were used after DESeq2 normalization ("normalized counts") to remove any depth sequencing bias. Each dot represents one individual. Dots are colored according to sex. Sex variable has two levels: female, male. Age group has five levels: [20-30], ]30-40], ]40-50], ]50-60], and ]60-70].

(**A**) Proportion of TE read counts among all read counts was modelized using a linear model with sex and age group as independent variables. From this model, the p-value testing the effect of sex on proportion of TE counts among all counts adjusted on age group was not significant.

(**B**) A Kruskal-Wallis test comparing proportion of TE read counts among all read counts across age groups was used in females and then on males. This test was not significant in females as well as in males.

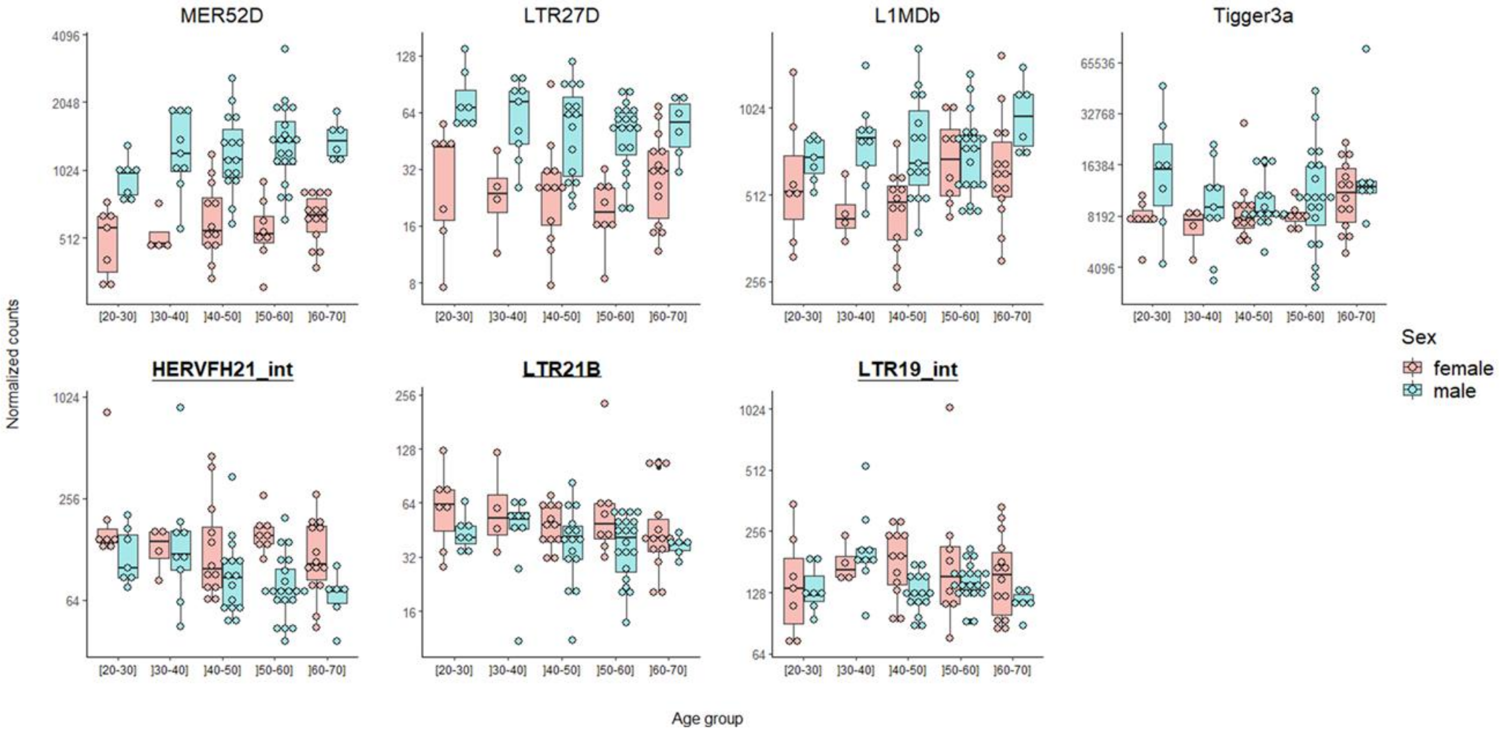


**Fig. S20. Boxplots of normalized counts according to sex and age group, for the TE subfamilies that are significantly differentially expressed between males and females in at least one age group or adjusted on age group in the filtered GTEx dataset (in the group including subjects with current and antecedents of cancer and cardiovascular diseases).**

Specific TE subfamily expression was estimated using the number of read counts after DESeq2 normalization ("normalized counts") to remove any depth sequencing bias. Each dot represents one individual. Dots are colored according to sex. Sex variable has two levels: female, male. Age group has five levels: [20-30], ]30-40], ]40-50], ]50-60], and ]60-70]. Y-axis is on a log2 scale.

All of these subfamilies had at least one copy on the Y chromosome and one copy on the X chromosome. Underlined bold TE subfamilies are enriched in the Y chromosome (binomial test adjusted p-value < 0.05 and number of observed copies located on Y chromosome > expected).

For more details on the group comparisons in which TE subfamilies were significantly differentially expressed, see **Table S12**.

A

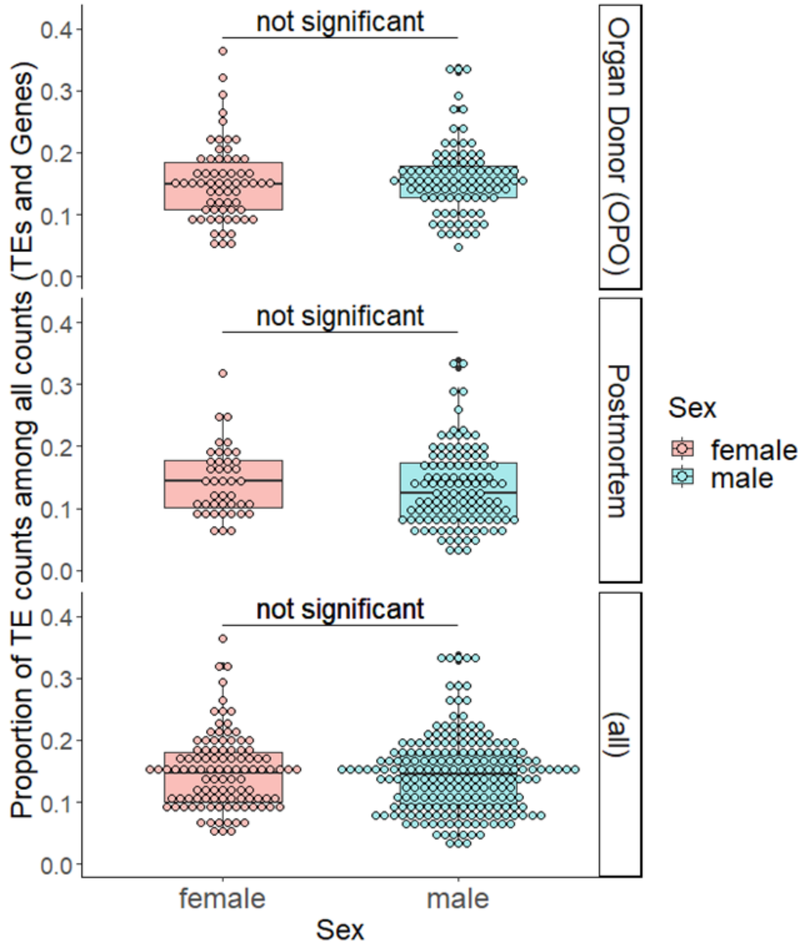

not significant
not significant
not significant
Proportion of TE counts among all counts (TEs and Genes)
0.0
0.1
0.2
0.3
0.4
Organ Donor (OPO)
Postmortem
(all)
female
male
Sex
Sex
female
male


256

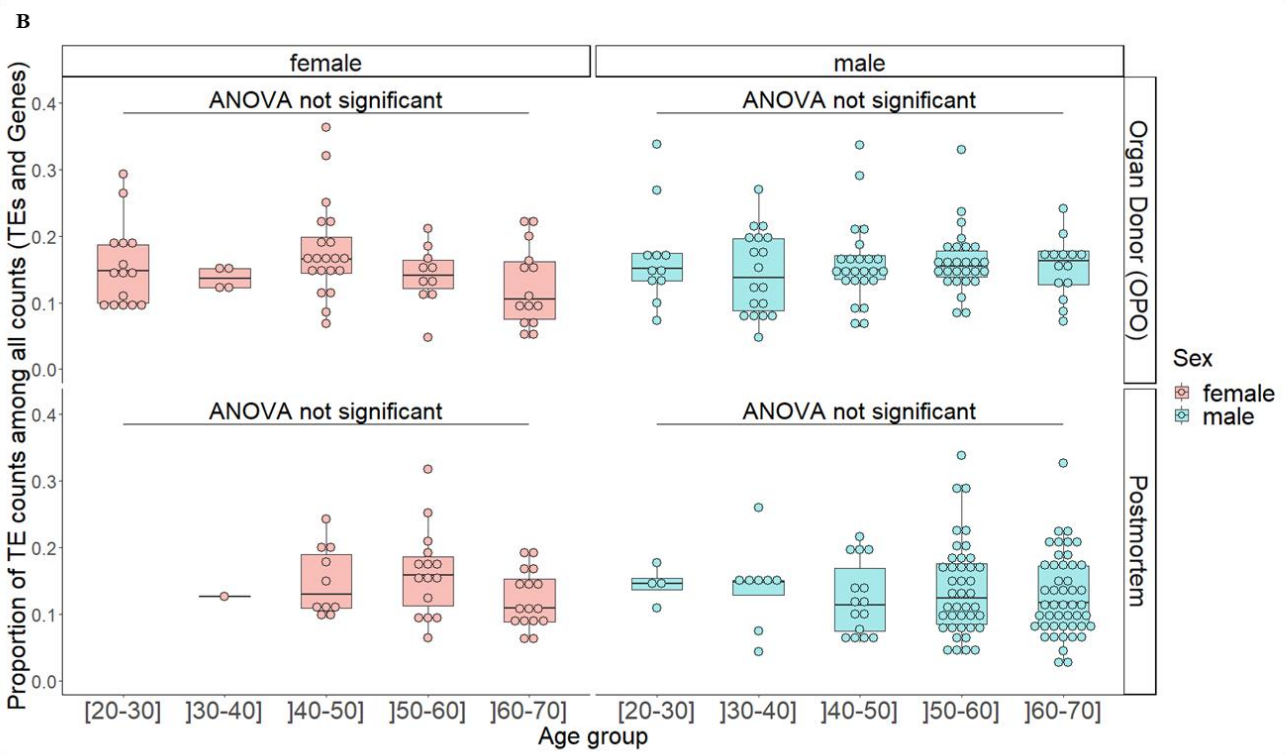


**Fig. S21. TE expression in the large GTEx dataset according to "sex" and "COHORT" variables (A) regardless of age group or (B) also according to age group.**

Global TE expression was measured in each karyotype (x-axis) as the proportion of TE read counts among all read counts (TEs and genes) (y-axis). In this calculation, read counts were used after DESeq2 normalization ("normalized counts") to remove any depth sequencing bias. Each dot represents one individual. Dots are colored according to sex. Sex variable has two levels: female, male. COHORT variable has two levels: Organ Donor (OPO), Postmortem.

(**A**) Proportion of TE read counts among all read counts was represented by sex for Organ Donor (OPO) individuals (top graph), for Postmortem individuals (middle graph), and without distinguishing them (bottom graph: "all").
Concerning the top and middle graphs, proportion of TE read counts among all read counts was modelized using a linear model with age group (5 levels: [20-30], ]30-40], ]40-50], ]50-60], and ]60-70]), sex, COHORT and interaction between the last two as independent variables. From this model, not significant p-values were obtained testing the age group adjusted effect of sex on proportion of TE read counts among all read counts in Organ Donor (OPO) individuals and in Postmortem individuals.
Concerning the bottom graph, proportion of TE read counts among all read counts was modelized using a linear model with age group, sex and COHORT as independent variables. From this model, the p-value testing the effect of sex on proportion of TE read counts among all read counts adjusted on age group and COHORT was not significant.

(**B**) A plot was drawn and an ANOVA model comparing proportion of TE read counts among all read counts across age groups was used in each sex*COHORT levels (female*Organ Donor (OPO), male*Organ Donor (OPO), female*Postmortem, and male*Postmortem). P-values testing the effect of the "age group" variable were not significant in each of the cases considered. P-value of age group effect adjusted on COHORT and sex was also not significant (p-value = 0.198) using an ANOVA model in the whole dataset.

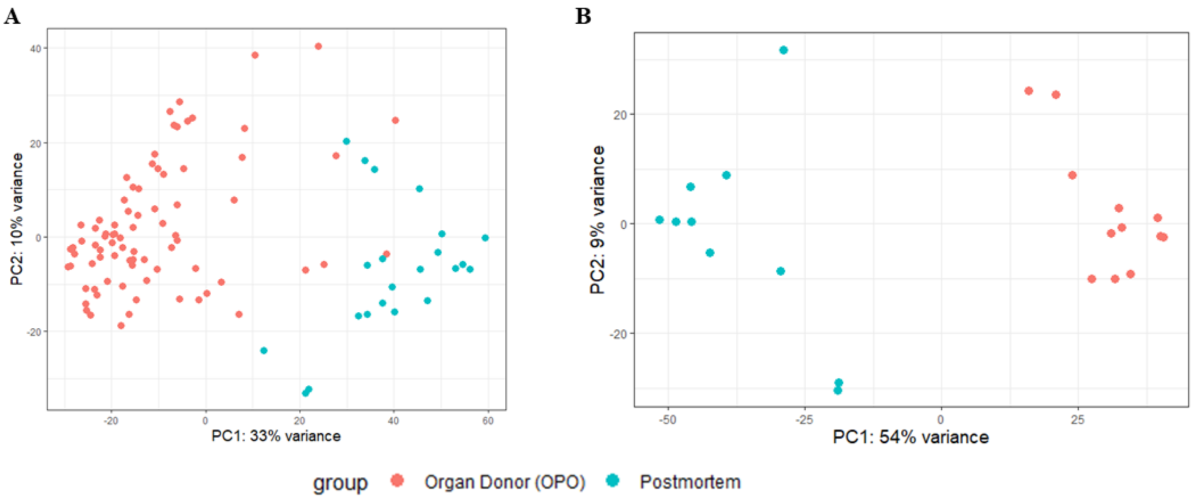


**Fig. S22. PCA analysis of GTEx expression data performed on the 500 genes explaining most of the variance for the identification of confounding factors considering (A) all of the 104 individuals with optimal data for use in the analysis after application of selection criteria and (B) only individuals in the 60-70 year-old group (n = 22/104).**

Each dot represents one individual. Dots are colored according to the two levels of the “COHORT” variable: “Organ Donor (OPO)” and “Postmortem”. PC: Principal Component.

### Supplementary tables

**Table S1. List of sequenced individuals and details on RNA-seq data in the gonosome aneuploidy dataset.**

| Identifier | Karyotype | Sex | Age (years) | Batch | Total number of read pairs | Pseudo-alignment rate Kallisto (genes) | Alignment rate TEcount (TE) |
|---|---|---|---|---|---|---|---|
| Txx20a | 46,XX | Female | 63 | 1 | 184223353 | 59.94% | 19.35% |
| Txx21b | 46,XX | Female | 35 | 2 | 132285589 | 60.39% | 15.65% |
| Txx22a | 46,XX | Female | 32 | 1 | 205687025 | 62.32% | 17.74% |
| Txx23a | 46,XX | Female | 29 | 1 | 169410086 | 63.82% | 17.52% |
| Txx24a | 46,XX | Female | 25 | 1 | 170479762 | 57.81% | 16.40% |
| Txx25b | 46,XX | Female | 51 | 2 | 167418680 | 48.30% | 16.08% |
| Txxy20b | 47,XXY* | Male | 30 | 2 | 153002185 | 53.52% | 12.36% |
| Txxy22a | 47,XXY | Male | 20 | 1 | 185535783 | 42.63% | 14.59% |
| Txxy23a | 47,XXY | Male | 34 | 1 | 183001944 | 47.65% | 14.72% |
| Txxy30a | 47,XXY | Male | 18 | 2 | 177627640 | 49.05% | 11.20% |
| Txxy31a | 47,XXY | Male | 17 | 2 | 149962790 | 45.56% | 11.36% |
| Txxy32a | 47,XXY | Male | 30 | 2 | 156062964 | 52.10% | 13.37% |
| Txxy33a | 47,XXY | Male | 20 | 2 | 171080372 | 50.14% | 11.57% |
| Txxyb | 47,XXY | Male | 18 | 1 | 173366155 | 58.29% | 14.32% |
| Txy20a | 46,XY | Male | 27 | 1 | 186765891 | 64.23% | 17.97% |
| Txy21a | 46,XY | Male | 56 | 1 | 185880218 | 56.05% | 14.16% |
| Txy22a | 46,XY | Male | 27 | 1 | 188136209 | 62.55% | 17.84% |
| Txy23b | 46,XY | Male | 26 | 2 | 143693391 | 44.53% | 11.26% |
| Txy24a | 46,XY | Male | 50 | 1 | 174258860 | 53.70% | 15.90% |
| Txy25b | 46,XY | Male | 63 | 2 | 134285944 | 53.05% | 17.38% |
| Txyy20a | 47,XYY | Male | 39 | 1 | 184316111 | 46.90% | 14.69% |
| Txyy30a | 47,XYY | Male | 34 | 2 | 174829021 | 40.65% | 12.61% |
| Txyy31a | 47,XYY* | Male | 30 | 2 | 161160366 | 46.24% | 10.91% |
| Txyyb | 47,XYY | Male | 15 | 1 | 188386148 | 54.03% | 14.27% |

TE: transposable elements.

*All 47,XXY males and 47,XYY males had homogenous 47,XXY or 47,XYY karyotype respectively except for Txxy20b individual (95% of mitoses were XXY, 3% XY, and 2% XXXY) and Txyy31a individual (97% of mitoses were XXY and 3% XY).

**Table S2. Percentages of significantly differentially expressed protein-coding genes after batch adjustment in the gonosome aneuploidy dataset in each of the 6 pairwise comparisons of karyotypes and according to the gene localization:total (n = 19947 genes), autosomes (n = 19035 genes), X chromosome (n = 851 genes), or Y chromosome (n = 66 genes including PAR, n = 48 genes excluding PAR)**

| Variation | In XY compared to XX | In XXY compared to XX | In XYY compared to XX | In XXY compared to XY | In XYY compared to XY | In XYY compared to XXY |
|---|---|---|---|---|---|---|
| Upregulated (%) – Total (19947 genes) | 19 (0.10%) | 726 (3.64%) | 518 (2.60%) | 104 (0.52%) | 30 (0.15%) | 35 (0.18%) |
| Downregulated (%) – Total (19947 genes) | 18 (0.09%) | 1167 (5.85%) | 992 (4.97%) | 37 (0.19%) | 28 (0.14%) | 15 (0.08%) |
| Upregulated (%) – Autosomes (19035 genes) | 5 (0.03%) | 692 (3.64%) | 480 (2.52%) | 87 (0.46%) | 19 (0.10%) | 29 (0.15%) |
| Downregulated (%) – Autosomes (19035 genes) | 4 (0.02%) | 1113 (5.85%) | 936 (4.92%) | 36 (0.19%) | 26 (0.14%) | 11 (0.06%) |
| Upregulated (%) – Xchr (851 genes) | 1 (0.12%) | 21 (2.47%) | 25 (2.94%) | 17 (2.00%) | 6 (0.71%) | 0 (0%) |
| Downregulated (%) – Xchr (851 genes) | 14 (1.65%) | 41 (4.82%) | 43 (5.05%) | 1 (0.12%) | 1 (0.12%) | 4 (0.47%) |
| Upregulated (%) – Ychr including PAR (66 genes) | 13 (19.70%) | 19 (28.79%) | 20 (30.30%) | 3 (4.55%) | 9 (13.64%) | 6 (9.09%) |
| Downregulated (%) – Ychr including PAR (66 genes) | 0 (0%) | 0 (0%) | 0 (0%) | 0 (0%) | 0 (0%) | 0 (0%) |
| Upregulated (%) – Ychr excluding PAR (48 genes) | 13 (27.08%) | 13 (27.08%) | 13 (27.08%) | 0 (0%) | 5 (10.42%) | 6 (12.50%) |
| Downregulated(%) – Ychr excluding PAR (48 genes) | 0 (0%) | 0 (0%) | 0 (0%) | 0 (0%) | 0 (0%) | 0 (0%) |

PAR: pseudo-autosomal region.

**Table S3. All the significantly differentially expressed protein-coding genes according to karyotype localized on the Y chromosome in the gonosome aneuploidy dataset likelihood-ratio test.**

| ENSG identifier | Gene name | Gene description | In PAR ? |
|---|---|---|---|
| ENSG00000114374 | *USP9Y* | ubiquitin specific peptidase 9 Y-linked | No |
| ENSG00000067048 | *DDX3Y* | DEAD-box helicase 3 Y-linked | No |
| ENSG00000183878 | *UTY* | ubiquitously transcribed tetratricopeptide repeat containing, Y-linked | No |
| ENSG00000012817 | *KDM5D* | lysine demethylase 5D | No |
| ENSG00000099725 | *PRKY* | protein kinase Y-linked (pseudogene) | No |
| ENSG00000129824 | *RPS4Y1* | ribosomal protein S4 Y-linked 1 | No |
| ENSG00000198692 | *EIF1AY* | eukaryotic translation initiation factor 1A Y-linked | No |
| ENSG00000067646 | *ZFY* | zinc finger protein Y-linked | No |
| ENSG00000154620 | *TMSB4Y* | thymosin beta 4 Y-linked | No |
| ENSG00000165246 | *NLGN4Y* | neuroligin 4 Y-linked | No |
| ENSG00000280969 | *RPS4Y2* | ribosomal protein S4 Y-linked 2 | No |
| ENSG00000169084 | *DHRSX* | dehydrogenase/reductase X-linked | Yes |
| ENSG00000169093 | *ASMTL* | acetylserotonin O-methyltransferase like | Yes |
| ENSG00000197976 | *AKAP17A* | A-kinase anchoring protein 17A | Yes |
| ENSG00000169789 | *PRY* | PTPN13 like Y-linked | No |
| ENSG00000169807 | *PRY2* | PTPN13 like Y-linked 2 | No |
| ENSG00000182378 | *PLCXD1* | phosphatidylinositol specific phospholipase C X domain containing 1 | Yes |
| ENSG00000214717 | *ZBED1* | zinc finger BED-type containing 1 | Yes |
| ENSG00000182162 | *P2RY8* | P2Y receptor family member 8 | Yes |
| ENSG00000124334 | *IL9R* | interleukin 9 receptor | Yes |

Top genes are those with the lowest adjusted p-values.
PAR: pseudo-autosomal region.

**Table S4. All the significantly differentially expressed protein-coding genes according to karyotype localized on the X chromosome in the gonosome aneuploidy dataset likelihood-ratio test.**

| ENSG identifier | Gene name | Gene description |
|---|---|---|
| ENSG00000184368 | MAP7D2 | MAP7 domain containing 2 |
| ENSG00000089472 | HEPH | hephaestin |
| ENSG00000169084 | DHRSX | dehydrogenase/reductase X-linked |
| ENSG00000169093 | ASMTL | acetylserotonin O-methyltransferase like |
| ENSG00000169249 | ZRSR2 | zinc finger CCCH-type, RNA binding motif and serine/arginine rich 2 |
| ENSG00000011201 | ANOS1 | anosmin 1 |
| ENSG00000188937 | NYX | nyctalopin |
| ENSG00000198034 | RPS4X | ribosomal protein S4 X-linked |
| ENSG00000130021 | PUDP | pseudouridine 5'-phosphatase |
| ENSG00000005889 | ZFX | zinc finger protein X-linked |
| ENSG00000173674 | EIF1AX | eukaryotic translation initiation factor 1A X-linked |
| ENSG00000197976 | AKAP17A | A-kinase anchoring protein 17A |
| ENSG00000147050 | KDM6A | lysine demethylase 6A |
| ENSG00000178947 | SMIM10L2A | small integral membrane protein 10 like 2A |
| ENSG00000182378 | PLCXD1 | phosphatidylinositol specific phospholipase C X domain containing 1 |
| ENSG00000069509 | FUNDC1 | FUN14 domain containing 1 |
| ENSG00000126012 | KDM5C | lysine demethylase 5C |
| ENSG00000006757 | PNPLA4 | patatin like phospholipase domain containing 4 |
| ENSG00000183943 | PRKX | protein kinase cAMP-dependent X-linked catalytic subunit |
| ENSG00000165704 | HPRT1 | hypoxanthine phosphoribosyltransferase 1 |
| ENSG00000130962 | PRRG1 | proline rich and Gla domain 1 |
| ENSG00000180879 | SSR4 | signal sequence receptor subunit 4 |
| ENSG00000186462 | NAP1L2 | nucleosome assembly protein 1 like 2 |
| ENSG00000184515 | BEX5 | brain expressed X-linked 5 |
| ENSG00000102287 | GABRE | gamma-aminobutyric acid type A receptor subunit epsilon |
| ENSG00000214717 | ZBED1 | zinc finger BED-type containing 1 |
| ENSG00000147123 | NDUFB11 | NADH:ubiquinone oxidoreductase subunit B11 |
| ENSG00000182162 | P2RY8 | P2Y receptor family member 8 |
| ENSG00000086712 | TXLNG | taxilin gamma |
| ENSG00000126756 | UXT | ubiquitously expressed prefoldin like chaperone |
| ENSG00000155966 | AFF2 | ALF transcription elongation factor 2 |
| ENSG00000196459 | TRAPPC2 | trafficking protein particle complex subunit 2 |
| ENSG00000257529 | RPL36A-HNRNPH2 | RPL36A-HNRNPH2 readthrough |
| ENSG00000198088 | NUP62CL | nucleoporin 62 C-terminal like |
| ENSG00000182712 | CMC4 | C-X9-C motif containing 4 |
| ENSG00000089289 | IGBP1 | immunoglobulin binding protein 1 |
| ENSG00000047230 | CTPS2 | CTP synthase 2 |
| ENSG00000068985 | PAGE1 | PAGE family member 1 |

| | | |
|---|---|---|
| ENSG00000180964 | TCEAL8 | transcription elongation factor A like 8 |
| ENSG00000277203 | None | novel transcript |
| ENSG00000125354 | SEPTIN6 | septin 6 |
| ENSG00000065923 | SLC9A7 | solute carrier family 9 member A7 |
| ENSG00000186310 | NAP1L3 | nucleosome assembly protein 1 like 3 |
| ENSG00000205542 | TMSB4X | thymosin beta 4 X-linked |
| ENSG00000183837 | PNMA3 | PNMA family member 3 |
| ENSG00000188706 | ZDHHC9 | zinc finger DHHC-type palmitoyltransferase 9 |
| ENSG00000172943 | PHF8 | PHD finger protein 8 |
| ENSG00000147403 | RPL10 | ribosomal protein L10 |
| ENSG00000198932 | GPRASP1 | G protein-coupled receptor associated sorting protein 1 |
| ENSG00000124334 | IL9R | interleukin 9 receptor |
| ENSG00000186871 | ERCC6L | ERCC excision repair 6 like, spindle assembly checkpoint helicase |
| ENSG00000123595 | RAB9A | RAB9A, member RAS oncogene family |
| ENSG00000183479 | TREX2 | three prime repair exonuclease 2 |
| ENSG00000094631 | HDAC6 | histone deacetylase 6 |
| ENSG00000165775 | FUNDC2 | FUN14 domain containing 2 |
| ENSG00000176896 | TCEANC | transcription elongation factor A N-terminal and central domain containing |
| ENSG00000158423 | RIBC1 | RIB43A domain with coiled-coils 1 |
| ENSG00000169239 | CA5B | carbonic anhydrase 5B |
| ENSG00000125356 | NDUFA1 | NADH:ubiquinone oxidoreductase subunit A1 |
| ENSG00000079482 | OPHN1 | oligophrenin 1 |
| ENSG00000215301 | DDX3X | DEAD-box helicase 3 X-linked |
| ENSG00000169057 | MECP2 | methyl-CpG binding protein 2 |
| ENSG00000232119 | MCTS1 | MCTS1 re-initiation and release factor |
| ENSG00000029993 | HMGB3 | high mobility group box 3 |
| ENSG00000102053 | ZC3H12B | zinc finger CCCH-type containing 12B |
| ENSG00000198918 | RPL39 | ribosomal protein L39 |
| ENSG00000284987 | None | novel transcript |
| ENSG00000160131 | VMA21 | vacuolar ATPase assembly factor VMA21 |
| ENSG00000241343 | RPL36A | ribosomal protein L36a |
| ENSG00000131174 | COX7B | cytochrome c oxidase subunit 7B |
| ENSG00000238269 | PAGE2B | PAGE family member 2B |
| ENSG00000102225 | CDK16 | cyclin dependent kinase 16 |
| ENSG00000156709 | AIFM1 | apoptosis inducing factor mitochondria associated 1 |
| ENSG00000130741 | EIF2S3 | eukaryotic translation initiation factor 2 subunit gamma |
| ENSG00000185515 | BRCC3 | BRCA1/BRCA2-containing complex subunit 3 |
| ENSG00000173275 | ZNF449 | zinc finger protein 449 |
| ENSG00000130826 | DKC1 | dyskerin pseudouridine synthase 1 |

Top genes are those with the lowest adjusted p-values.

**Table S5. Differentially expressed TE subfamilies in samples with different karyotypes among the 1246 explored in the gonosome aneuploidy dataset.**

| Variation | In XY compared to XX | In XXY compared to XX | In XYY compared to XX | In XXY compared to XY | In XYY compared to XY | In XYY compared to XXY |
|---|---|---|---|---|---|---|
| Upregulated TE subfamilies | 7 subfamilies (0.6%):<br><br>LTR22B2<br>MSTB1-int<br>LTR19-int*<br>MER70B<br>LTR19A<br>MER51C<br>LTR25-int | 79 subfamilies (6.3%):<br><br>LTR22B2<br>LTR19-int*<br>MSTB1-int<br>LTR19A<br>HERV9-int<br>MER51C<br>LTR25-int<br>MER61F<br>HERV3-int<br>LTR15*<br>LTR5B<br>LTR68<br>MER115<br>LTR1F1*<br>THE1A-int<br>MER70B<br>MER126<br>LTR13<br>**HERVL18-int**<br>LTR10G<br>**UCON46**<br>**hAT-N1_Mam**<br>Charlie15a<br>MER91B<br>LTR5_Hs<br>ORSL<br>MER52D*<br>LTR10F<br>LTR19B<br>PRIMA41-int<br>LTR1F2<br>MER96B<br>HERV15-int<br>LTR35<br>LTR14<br>MLT1G<br>MLT1M<br>MER50C<br>MER21B<br>LTR53<br>MER21A<br>ERVL-int<br>MER34C2<br>MLT1A-int<br>**LTR18A**<br>**MamRep38**<br>LTR22B<br>MER9a1<br>MER68-int<br>MER30B*<br>LTR12B<br>LTR4<br>ERV3-16A3_LTR<br>MamTip3<br>SVA_D | 43 subfamilies (3.5%):<br><br>LTR22B2<br>MSTB1-int<br>LTR19-int*<br>LTR19A<br>MER51C<br>MER61F<br>HERV9-int<br>LTR25-int<br>MER70B<br>LTR12B<br>LTR68<br>HERV3-int<br>LTR40A1<br>LTR15*<br>HERVK22-int<br>LTR10F<br>LTR25<br>MER9a1<br>MER52D*<br>LTR1F1*<br>LTR5B<br>MER50C<br>LTR13<br>MER41D*<br>HERV1_LTRe<br>LTR4<br>MamGypLTR3a<br>LTR5_Hs<br>LTR19B<br>MLT1N2<br>THE1A-int<br>MER92B<br>LTR66<br>**X7C_LINE**<br>MER4A1<br>MER34C2<br>Charlie22a<br>MER91B<br>LTR17<br>Ricksha_b<br>MER21B<br>HERVL66-int<br>MLT2B2 | 0 subfamily | 0 subfamily | 0 subfamily |

| | | | | | | |
|---|---|---|---|---|---|---|
| | | **MLT1C2-int**<br>MER52A<br>**UCON86***<br>LTR22C0*<br>LTR26D<br>MER74B*<br>**X7C_LINE**<br>LTR66<br>HERVK22-int<br>MER66B<br>MSTD-int<br>HERV1_LTRe<br>THE1B-int<br>HERVK3-int<br>MLT1N2<br>**MER51E***<br>MER117<br>**UCON89**<br>LTR23-int<br>MLT1E3<br>LTR40A1<br>LTR3B_<br>MER91A<br>MER92B | | | | |
| Downregulated TE subfamilies | 1 subfamily (0.1%):<br><br>**UCON75** | 16 subfamilies (1.3%):<br><br>HERVH-int<br>LTR46-int<br>**HERVH48-int**<br>HERVFH21-int*<br>UCON132b<br>**MER41G**<br>Alu<br>**LTR1C**<br>LTR10B1<br>LTR37B<br>**UCON5**<br>FordPrefect*<br>LTR7Y*<br>**UCON81**<br>LTR73<br>**UCON1** | 17 subfamilies (1.4%):<br><br>HERVH-int<br>**HERVH48-int**<br>UCON132b<br>HERV1_LTRc<br>LTR46-int<br>**MER41G**<br>HERVFH21-int*<br>AluYj4<br>FordPrefect*<br>AluYb9<br>**UCON5**<br>LTR10B1<br>LTR37B<br>**MER96**<br>**UCON1**<br>HERV1_I-int<br>Alu | 0 subfamily | 0 subfamily | 0 subfamily |

TE subfamily names are classified according to adjusted p-values in ascending order. The coloured names are names found in at least three comparisons.
Underlined TE subfamily names are those enriched in the Y chromosome (binomial test adjusted p-value < 0.05 and number of observed copies located on Y chromosome > expected). Bold TE subfamily names are TE family with no copy located in the Y chromosome in the TE sequence reference file.

* found significantly differentially expressed in any comparison in the filtered GTEx dataset.

Variation is upregulated if TE subfamily is upregulated in the first karyotype (e.g. XYY in the "XYY karyotype vs XX karyotype" comparison) compared to the second (e.g. XX in the "XYY karyotype vs XX karyotype" comparison). Variation is downregulated if TE subfamily is downregulated in the first karyotype (e.g. XYY in the "XYY karyotype vs XX karyotype" comparison) compared to the second (e.g. XX in the "XYY karyotype vs XX karyotype" comparison).

**Table S6. Overlap between intragenic regions of differentially expressed genes and differentially expressed TE subfamilies**

| Pairwise karyotype comparison | Overlap between | Expected proportion* | Observed proportion** | P-value (exact Binomial test) |
|---|---|---|---|---|
| XY are compared to XX | Upregulated TEs and upregulated genes | 0.00340 (206/60564) | 0.04762 (2/42) | 0.009 |
| XY are compared to XX | Downregulated TEs and downregulated genes | 0.00007 (4/60564) | 0 (0/35) | 1 |
| XXY are compared to XX | Upregulated TEs and upregulated genes | 0.13252 (8026/60564) | 0.20994 (228/1086) | <0.001 |
| XXY are compared to XX | Downregulated TEs and downregulated genes | 0.03537 (2142/60564) | 0.05473 (88/1608) | <0.001 |
| XYY are compared to XX | Upregulated TEs and upregulated genes | 0.06681 (4046/60564) | 0.09986 (71/711) | <0.001 |
| XYY are compared to XX | Downregulated TEs and downregulated genes | 0.05417 (3281/60564) | 0.09098 (123/1352) | <0.001 |

TE: transposable element.

*Expected proportion corresponds to genes containing at least one TE copy from one of the upregulated TE subfamilies among all the 60564 genes or to genes containing at least one TE copy from one of the downregulated TE subfamilies among all the 60564 genes according to what is indicated in the “overlap between” column (details of the calculation are in brackets).

**Observed proportion corresponds to upregulated genes containing at least one TE copy from one of the upregulated TE subfamilies among all the upregulated genes or to downregulated genes containing at least one TE copy from one of the downregulated TE subfamilies among all the downregulated genes according to what is indicated in the “overlap between” column (details of the calculation are in brackets).

**Table S7. Repartition of GTEx individuals according to age group and sex in the filtered GTEx dataset (no disease group).**

| Age group (years) | [20-30] | ]30-40] | ]40-50] | ]50-60] | ]60-70] | Total |
|---|---|---|---|---|---|---|
| Females | 7 | 3 | 11 | 6 | 8 | 35 |
| Males | 7 | 8 | 13 | 13 | 4 | 45 |
| Total | 14 | 11 | 24 | 19 | 12 | 80 |

**Table S8. Repartition of GTEx individuals according to age group and sex in the filtered GTEx dataset (group including individuals with current or antecedents of cancer or cardiovascular diseases).**

| Age group (years) | [20-30] | ]30-40] | ]40-50] | ]50-60] | ]60-70] | Total |
|---|---|---|---|---|---|---|
| Females | 7 | 4 | 12 | 8 | 14 | 45 |
| Males | 7 | 9 | 15 | 20 | 6 | 57 |
| Total | 14 | 13 | 27 | 28 | 20 | 102 |

**Table S9. Differentially expressed TE subfamilies among the 1246 explored in the filtered GTEx (no disease group) dataset between all the pairwise comparisons of age groups in each sex.**

| Age group (years) | Variation | In males | In females | Overall adjusted on sex |
|---|---|---|---|---|
| In ]30-40] compared to [20-30] | Upregulated | MER51E*<br>MER30B*<br>LTR1F1*<br>MER74C<br>HERV35I-int<br>MER110A<br>LTR24<br>FordPrefect*<br>MLT1H1-int<br>LTR1A1<br>Looper<br>LTR9C<br>MER65C<br>LTR9B | 0 | 0 |
| | Downregulated | 0 | 0 | 0 |
| In ]40-50] compared to [20-30] | Upregulated | 0 | 0 | 0 |
| | Downregulated | 0 | 0 | 0 |
| In ]50-60] compared to [20-30] | Upregulated | 0 | 0 | HERV35I-int |
| | Downregulated | LTR48B | 0 | LTR48B |
| In ]60-70] compared to [20-30] | Upregulated | 0 | 0 | LTR10E<br>MLT2B5<br>Tigger11a<br>MER121<br>SVA_A |
| | Downregulated | 0 | 0 | 0 |
| In ]40-50] compared to ]30-40] | Upregulated | 0 | 0 | LTR35B |
| | Downregulated | HSMAR2<br>MER110A<br>L1M2c<br>MER51E*<br>MLT1H1-int<br>L1PA12<br>LTR24<br>HUERS-P1-int<br>LTR1F1* | 0 | HSMAR2<br>L1M2c<br>MER41D* |
| In ]50-60] compared to ]30-40] | Upregulated | 0 | 0 | 0 |
| | Downregulated | MER51E*<br>L1M2c<br>UCON49<br>MER110A<br>UCON86*<br>LTR24<br>MER74C<br>MER74B*<br>Arthur1C<br>UCON38<br>L1P4e<br>MER57C1<br>HERVFH21-int*<br>HSMAR2<br>LTR9C<br>MER8<br>L4_A_Mam | 0 | L1M2c<br>MER51E*<br>LTR24<br>MER74C<br>UCON38<br>MER74B*<br>UCON49<br>Arthur1C<br>MER8<br>LTR86C<br>L4_A_Mam<br>MER57C1<br>LTR9C<br>MER110A<br>LTR22C0* |
| In ]60-70] compared to ]30- | Upregulated | 0 | 0 | MER99 |
| | Downregulated | MLT1E1A-int | 0 | MER74C |

| | | | | |
|---|---|---|---|---|
| 40] | | MER74C<br>LTR15*<br>MER110A | | MER51E*<br>HERVFH21-int*<br>LTR21A<br>MER65C<br>LTR24<br>LTR19-int*<br>LTR81 |
| In ]50-60] compared to ]40-50] | Upregulated | 0 | 0 | 0 |
| | Downregulated | 0 | 0 | 0 |
| In ]60-70] compared to ]40-50] | Upregulated | 0 | 0 | 0 |
| | Downregulated | 0 | 0 | 0 |
| In ]60-70] compared to ]50-60] | Upregulated | 0 | 0 | MER51-int<br>L1M3e<br>ORSL-2a<br>MamGyp-int |
| | Downregulated | 0 | 0 | 0 |

TE subfamily names are classified according to adjusted p-values in ascending order. The coloured names are names found in at least three comparisons. Note that none of these TE subfamilies were significantly differentially expressed according to the sex variable. Variation is upregulated if TE subfamily is upregulated in the first age group (e.g. ]30-40] in the "]30-40] vs [20-30]" comparison) compared to the second (e.g. [20-30] in the "]30-40] vs [20-30]" comparison). Variation is downregulated if TE subfamily is downregulated in the first age group (e.g. ]30-40] in the "]30-40] vs [20-30]" comparison) compared to the second (e.g. [20-30] in the "]30-40] vs [20,30]" comparison). Underlined TE subfamilies correspond to those for which we found a significant interaction indicating a log2-fold change in males compared to females that was higher in the ]30-40] age group than in the [20-30] age group: MER74C, MER110A.

* found significantly differentially expressed in some pairwise comparisons of karyotypes in the gonosome aneuploidy dataset.

**Table S10. Differentially expressed TE subfamilies among the 1246 explored in the filtered GTEx dataset (no disease group) between males and females in each age group.**

| Age group (years) | Upregulated TE subfamilies in males compared to females | Downregulated TE subfamilies in males compared to females |
|---|---|---|
| [20-30] | 0 | 0 |
| ]30-40] | 0 | 0 |
| ]40-50] | MER52D*<br>LTR27D | 0 |
| ]50-60] | MER52D*<br>LTR27D | 0 |
| ]60-70] | 0 | 0 |
| Overall adjusted on age group | MER52D*<br>LTR27D<br>L1MDb | 0 |

TE subfamily names are classified according to adjusted p-values in ascending order. The coloured names are names found in at least two comparisons.
Note that none of these TE subfamilies were significantly differentially expressed according to the age group variable. All of these 3 subfamilies had at least one copy on the Y chromosome and one copy on the X chromosome, but none are enriched on the Y chromosome.

* found significantly differentially expressed in some pairwise comparisons of karyotypes in the gonosome aneuploidy dataset.

**Table S11. Differentially expressed TE subfamilies among the 1246 explored in the filtered GTEx (group including subjects with current or antecedents of cancer or cardiovascular diseases) dataset between all the pairwise comparisons of age groups in each sex.**

| Age group (years) | Variation | In males | In females | Overall adjusted on sex |
|---|---|---|---|---|
| In ]30-40] compared to [20-30] | Upregulated | MER51E*<br>MER30B*<br>MER74C<br>MER110A<br>Looper<br>LTR1F1*<br>HERV35I-int<br>LTR9C<br>LTR24<br>MLT1H1-int | 0 | 0 |
| | Downregulated | 0 | 0 | 0 |
| In ]40-50] compared to [20-30] | Upregulated | 0 | 0 | 0 |
| | Downregulated | 0 | 0 | 0 |
| In ]50-60] compared to [20-30] | Upregulated | 0 | 0 | HERV35I-int<br>MamRep4096<br>LTR1C3<br>L1MD1 |
| | Downregulated | 0 | 0 | MER65-int |
| In ]60-70] compared to [20-30] | Upregulated | 0 | Tigger11a | LTR10E<br>Tigger11a<br>MLT2B5<br>SVA_A<br>HERV9N-int<br>MER121<br>HERVI-int<br>MER30 |
| | Downregulated | 0 | 0 | MER65-int |
| In ]40-50] compared to ]30-40] | Upregulated | 0 | 0 | 0 |
| | Downregulated | HSMAR2<br>MER110A<br>MER51E*<br>L1M2c<br>MLT1H1-int<br>LTR24<br>L1PA12<br>Eulor6E<br>MER74C<br>LTR15*<br>MER30B*<br>L1MA7<br>L1MD<br>LTR86A2<br>L1PB2<br>Arthur1C<br>L4_A_Mam<br>LTR37-int | 0 | HSMAR2 |
| In ]50-60] compared to ]30-40] | Upregulated | 0 | 0 | 0 |
| | Downregulated | MER51E*<br>MER74C<br>L1M2c<br>LTR24<br>MER74B*<br>HSMAR2<br>MER113B<br>HERVFH21-int*,† | 0 | L1M2c<br>UCON38<br>LTR24<br>Helitron1Nb_Mam<br>MER74B*<br>L1M3e<br>MER8<br>HSMAR2 |

| | | | | |
|---|---|---|---|---|
| | | MER8<br>UCON86*<br>L4_A_Mam<br>MER110A<br>MER41D*<br>MER53<br>MLT1H1-int<br>UCON38<br>L1ME5<br>Arthur1C<br>L1M3e<br>L1ME3C<br>L1PA12<br>Looper<br>L1P2<br>LTR9C<br>L1MA7<br>MARNA<br>MER30B*<br>UCON49<br>L5<br>Cheshire<br>L1PB2<br>LTR38 | | L5<br>L4_A_Mam<br>MER41D* |
| In ]60-70] compared to ]30-40] | Upregulated | 0 | 0 | 0 |
| | Downregulated | MER74C<br>MLT1E1A-int<br>MER110A<br>MER51E<br>LTR5A<br>LTR24<br>LTR38<br>HERVFH21-int*,†<br>L1M2c<br>LTR21A<br>L1PA12<br>ERV24_Prim-int<br>LTR15* | 0 | LTR21A<br>L1M2c<br>HERVFH21-int*,†<br>LTR47A2 |
| In ]50-60] compared to ]40-50] | Upregulated | 0 | 0 | 0 |
| | Downregulated | 0 | 0 | 0 |
| In ]60-70] compared to ]40-50] | Upregulated | 0 | 0 | 0 |
| | Downregulated | 0 | 0 | 0 |
| In ]60-70] compared to ]50-60] | Upregulated | 0 | 0 | 0 |
| | Downregulated | 0 | 0 | 0 |

TE subfamily names are classified according to adjusted p-values in ascending order. The coloured names are names found in at least three comparisons. Variation is upregulated if TE subfamily is upregulated in the first age group (e.g. ]30-40] in the "]30-40] vs [20-30]" comparison) compared to the second (e.g. [20-30] in the "]30-40] vs [20-30]" comparison). Variation is downregulated if TE subfamily is downregulated in the first age group (e.g. ]30-40] in the "]30-40] vs [20-30]" comparison) compared to the second (e.g. [20-30] in the "]30-40] vs [20-30]" comparison).

Underlined TE subfamilies correspond to those for which we found a significant interaction: MER74C, MER110A, and MER51E showed a log2-fold change in males compared to females that was higher in the ]30-40] age group than in the [20-30] age group; MER74C and MER110A showed a log2-fold change in males compared to females that was lower in the ]60-70] age group than in the ]30-40] age group; MER110A showed a log2-fold change in males compared to females that was lower in the ]40-50] age group than in the ]30-40] age group.

* found significantly differentially expressed in some pairwise comparisons of karyotypes in the gonosome aneuploidy dataset.
† found significantly differentially expressed in males vs females in some age groups of the filtered GTEx dataset in the group including subjects with current or antecedents of cancer of cardiovascular diseases.

**Table S12. Differentially expressed TE subfamilies among the 1246 explored in the filtered GTEx dataset between males on females in each age group, in the group including subjects with current or antecedents of cancer of cardiovascular diseases.**

| Age group (years) | Upregulated TE subfamilies in males compared to females | Downregulated TE subfamilies in males compared to females |
|---|---|---|
| [20-30] | 0 | 0 |
| ]30-40] | MER52D* | 0 |
| ]40-50] | MER52D* | 0 |
| ]50-60] | MER52D*<br>LTR27D | 0 |
| ]60-70] | MER52D* | 0 |
| Overall adjusted on age group | MER52D*<br>LTR27D<br>L1MDb<br>Tigger3a | HERVFH21-int*,†<br>LTR21B<br>LTR19-int* |

TE subfamily names are classified according to adjusted p-values in ascending order. The coloured names are names found in at least two comparisons.

* found significantly differentially expressed in some pairwise comparisons of karyotypes in the gonosome aneuploidy dataset.

† found significantly differentially expressed in some pairwise comparisons of age groups in the filtered GTEx dataset in the group including subjects with current or antecedents of cancer of cardiovascular diseases.

**Table S13. Repartition of GTEx individuals according to age group and sex in the large GTEx dataset.**

| Age group (years) | [20-30] | ]30-40] | ]40-50] | ]50-60] | ]60-70] | Total |
|---|---|---|---|---|---|---|
| Females | 15 | 5 | 30 | 25 | 30 | 105 |
| Males | 15 | 26 | 39 | 68 | 61 | 209 |
| Total | 30 | 31 | 69 | 93 | 91 | 314 |

#### Supplementary data

**Data S1. List of all differentially expressed genes on X chromosome or Y chromosome according to karyotype using the likelihood-ratio test in the gonosome aneuploidy dataset, first considering all genes and then considering protein-coding genes only.**

**Data S2. List of all differentially expressed genes (regardless if they are protein-coding or not) in each pairwise karyotype comparison after batch adjustment in the gonosome aneuploidy dataset.**

**Data S3. GO analysis in the gonosome aneuploidy dataset.**
GO analysis with gprofiler in the gonosome aneuploidy dataset after adjustment on batch effect. For each pairwise karyotype comparison, four files were generated (one for upregulated genes and one for downregulated genes using all genes first and then using only protein-coding genes).

**Data S4. Y chromosome enrichment test for TE subfamilies.**

**Data S5. Overlap between genomic positions of upregulated (downregulated, respectively) genes and positions of upregulated (downregulated, respectively) TEs for each pairwise karyotype comparison (when applicable).**

**Data S6. Rosetta file.**
